# Reconfigurable pH-Responsive DNA Origami Lattices

**DOI:** 10.1101/2023.02.03.526959

**Authors:** Sofia Julin, Veikko Linko, Mauri A. Kostiainen

## Abstract

DNA nanotechnology enables straightforward fabrication of user-defined and nano-meter-precise templates for a cornucopia of different uses. To date, most of these DNA assemblies have been static, but dynamic structures are increasingly coming into view. The programmability of DNA not only allows encoding of the DNA object shape, but it may be equally used in defining the mechanism of action and the type of stimuli-responsiveness of the dynamic structures. However, these "robotic" features of DNA nanostructures are usually demonstrated for only small, discrete and device-like objects rather than for collectively behaving higher-order systems. Here, we show how a large-scale, two-dimensional (2D) and pH-responsive DNA origami -based lattice can be assembled on a mica substrate and further reversibly switched between two distinct states upon the pH change of the surrounding solution. The control over these two configurations is achieved by equipping the arms of the lattice-forming DNA origami units with "pH-latches" that form Hoogsteen-type triplexes at low pH. In a nutshell, we demonstrate how the electrostatic control over the adhesion and mobility of the DNA origami units on the surface can be used both in the large lattice formation (with the help of directed polymerization) and in the conformational switching of the whole lattice on the substrate. To further emphasize the feasibility of the method, we also demonstrate the formation of reconfigurable 2D gold nanoparticle lattices. We believe this work serves as an important milestone in bridging the nanometer-precise DNA origami templates and higher-order large-scale systems with the stimuli-induced dynamicity.

---

Recent advances in the field of nanotechnology have enabled the fabrication of a variety of nanoobjects with intriguing geometries and properties. However, for many applications, more complex, structurally well-defined nanomaterials in which the individual building blocks could interact with each other in a predefined manner would be highly desirable.^1–3^ Owing to the highly specific and predictable Watson-Crick base-pairing, DNA-based nanostructures have proven to be feasible templates for constructing precise nanoscale arrangements.^4,5^ For this, particularly the DNA origami technique allows the production of a wide range of well-defined two- and three-dimensional (2D and 3D) DNA nanostructures with high complexity and addressability.^6–8^ The rapidly emerged DNA origami design software^9–11^ have further paved the way for numerous sophisticated applications in e.g. nanomedicine,^12–15^ nanophotonics, ^16,17^ nanoelectronics^18^ and bottom-up nanofabrication.^19,20^

DNA origami is a versatile method, and therefore, it has also been used to construct large-scale hierarchical assemblies. ^21–24^ These DNA origami-based lattices could also serve as templates for controlling and directing the spatial arrangements of other compounds as shown for example by creating increasingly complex metal nanoparticle lattices using DNA origami frameworks.^25,26^ Thus far, most of such research has been focused on static assemblies, however, getting inspired by nature, the interests are increasingly shifting toward dynamic structures that undergo conformational changes in response to external stimuli, such as pH, salt concentration, light or temperature.^27^ Apart from a very few examples,^28–30^ the use of DNA origami for the construction of dynamic 2D and 3D lattices has been rather limited. Nevertheless, the library of already demonstrated small dynamic DNA-based devices^31,32^ suggests the DNA origami method could also be harnessed in building larger dynamic lattices and other highly ordered assemblies.^33^

In this work, we created a dynamic 2D DNA origami lattice that changes its configuration in response to the pH of the surrounding solution. For that, we designed a pliers-like DNA origami unit that serves as the basic building block of the lattice and that can be readily switched between an open "+"-shaped and a closed "X"-shaped state upon a pH change. The controlled dynamicity is achieved using pH-sensitive "latches", whose counterparts are positioned at the opposite arms of the unit (**Figure 1a**). These particular "pH-latches" are based on the pH-dependent, Hoogsteen-type DNA triplex formation,^34,35^ but it is noteworthy to mention that there also exist other pH-responsive constructs that could be equally implemented, such as the i-motif.^36^ First, we characterized the plain unit and its dynamic behavior by agarose gel electrophoresis (AGE) and transmission electron microscopy (TEM). By introducing "connector oligonucleotides", the units were selectively linked together, and subsequently, we were able to increase the complexity of our system. This was shown by assembling DNA origami dimers, one-dimensional (1D) DNA origami arrays (chains), and ultimately reconfigurable pH-sensitive 2D DNA origami lattices. Furthermore, the developed lattices could also serve as templates for other nanoscale compounds, which we demonstrated here by assembling dynamic pH-responsive 2D gold nanoparticle (AuNP) lattices.

**Figure 1:**
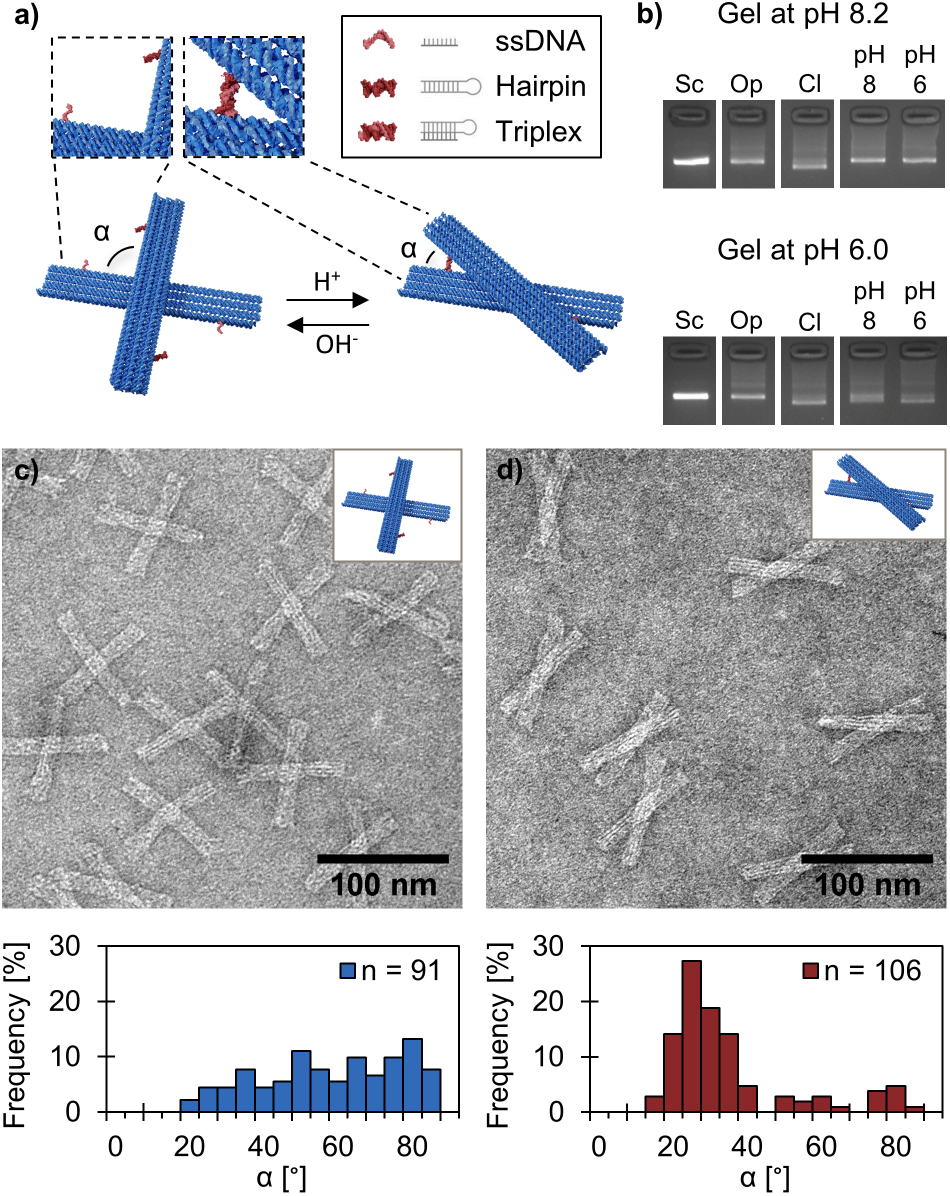
Design and characterization of the reconfigurable pH-sensitive DNA origami unit. (a) The DNA origami unit contains two bar-like arms (86 nm × 12 nm × 6 nm) that are connected through a pivot (two DNA scaffold crossover). The arms are equipped with two latches that consist of a 20-bp hairpin and a complementary 20-nt ssDNA counterpart. These two latch counterparts will form a DNA triplex when the solution pH is below the transition p*K* _a_ value (~7.2), and thus, the closing of the latches locks the arms of the unit at a fixed vertex angle, *α* ≈ 30°. Increasing the pH above the p*K* _a_ will open the latches and let the arms move freely between *α* ≈ 20–90°. (b) Analysis of different DNA origami units by agarose gel electrophoresis (AGE) at pH 8.2 (top panel) and pH 6.0 (bottom panel). The samples in the gel are scaffold (Sc), permanently open unit (Op), permanently closed unit (Cl) and unit with pH latches at pH 8.2 (pH 8) and pH 6.0 (pH 6). Transmission electron microscopy (TEM) images of the units with pH latches at (c) pH 8.2 and (d) pH 6.0. Both TEM images are negatively stained with 2 % (w/v) uranyl formate. The bottom panel shows the distribution of the angle, *α*, between the two arms of the unit. The number of individual structures analyzed for each sample is given by *n*.

## Results and Discussion

### Design and Characterization of the Reconfigurable pH-Responsive DNA Origami Unit

To assemble the pH-responsive and dynamic lattice, we first constructed and characterized the pH-sensitive DNA origami unit, the basic building block of the lattice. The pliers-like DNA origami unit consists of two bar-shaped arms (86 nm × 12 nm × 6 nm) that are connected to each other through the pivot which is two single-stranded DNA (ssDNA) scaffold crossovers (analogous to the Holliday junction) (**Figure 1a**). The unit is designed with two rationally engineered pH-sensitive "latches". Therefore, depending on the pH of the surrounding solution, the unit may adopt either an open (vertex angle between the arms varies freely from *α* ≈ 20° to *α* ≈ 90°) or a closed configuration (vertex angle *α* ≈ 30°). In more detail, the latches are staple-strand extensions and consist of two counterparts positioned on different arms of the unit: a hairpin with a 20-base pair (bp) double-stranded DNA (dsDNA) region and a complementary 20-nucleotide (nt) ssDNA sequence. At high pH, the hairpin and the ssDNA do not interact with each other thus allowing free rotation of the arms. At low pH, for one, these two counterparts can form a parallel DNA triplex through Hoogsteen interactions, which locks the two arms at a fixed position. Both pH latches have unique base sequences, but an identical T-A·T base content of 60%, which ensures that both latches have a transition pH value of p*K* _a_ ~ 7.2 and thus they will open/close at the same pH.^37^ However, the pH range at which the opening/closing takes place can be rationally tuned by adjusting the T-A·T base content of the latch sequences.^34,38^ In addition to the pH-sensitive unit, we also designed and prepared two control units: a permanently open unit with no latch sequences (Op) and a permanently closed unit (Cl), in which the pH-sensitive latch sequences have been replaced with complementary ssDNA overhangs.

To confirm both the correct folding of the units and the functionality of the pH-sensitive latches, poly-T passivated DNA origami units (8-nt polythymine extensions at each helix to avoid end-to-end stacking) were first analyzed by agarose gel electrophoresis (AGE) (**Figures 1b, S2**). The closed unit is more compact than the open unit and therefore the closed unit exhibits a higher electrophoretic mobility in the gel. This allows separation of these two configurations by AGE. The first gel was run at pH 8.2 (**Figure 1b, top panel**), which is above the p*K* _a_ value, and thus, it was also expected that samples prepared at pH 8.2 (initially open) and at pH 6.0 (initially closed) will both adapt the open configuration. This is indeed the case, as the both samples exhibit equal mobility which further matches the mobility of the permanently open (Op) control sample. The second gel (**Figure 1b, bottom panel**), was run at pH 6.0, which is well below the p*K* _a_ value. Here, the (initially closed) sample at pH 6.0 remains predominantly in its closed configuration, while the (initially open) sample prepared at pH 8.2 shows a slightly broader band. This indicates that the sample is a blend of both open and closed configurations due to the slow closing kinetics of the initially open unit.^37,38^ The opening kinetics is faster and therefore the initially closed unit will swiftly open in the pH 8.2 gel resulting in a clear and narrow band.

In addition to AGE, we also used TEM to characterize the DNA origami units at both pH 8.2 and pH 6.0. In both cases, TEM reveals distinct, correctly folded units (**Figures 1c,d, S3, S4**). At pH 8.2, the unit equipped with pH-sensitive latches adapted the open configuration with a wide (*α* ≈ 20–90°) and flat vertex angle distribution (**Figure 1c**). At pH 6.0, on the other hand, most of the units adapted the closed configuration with an vertex angle of *α* ≈ 30° (~ 75% of the units have vertex angles between 20–40°) (**Figure 1d**). Despite this pronounced and narrow vertex angle distribution, both TEM and AGE analysis additionally reveal that a small fraction of the pH-sensitive units still remains at the open configuration at pH 6.0. The same trend was also observed for the permanently closed control unit (at both pH 8.2 and pH 6.0, see **Figures S5, S6**) and even more pronounced at low cation concentrations (see **Figure S1**), indicating that the electrostatic repulsion between the two arms is strong enough to prevent some units from closing. Nevertheless, the observed closing yield is in good agreement with previously reported closing efficiencies for similar pH-responsive DNA origami structures.^39^

### Selective Assembly of DNA Origami Dimers

For the lattice formation it is crucial that the units are connected together in a programmable fashion without undesired interconnection of the top arm and the bottom arm that are located in different planes. In order to selectively connect only specific ends of the arms, we designed "connector oligonucleotides" for seven helices in each of the two arms (**Figures 2a, S36**). To further minimize the undesired interactions between the two arms, the connector oligonucleotides were arranged in different patterns for the top and the bottom arms of the unit, while the rest of the helix-ends remained untouched (blue helices in **Figure 2a** cross-section). In total 14 strands of the connector oligonucleotides (7 per each arm, 4 at one end and 3 at the other) contain a 3-nt long protruding 3’-end-overhang. Each overhang is complementary to a 3-nt long scaffold sequence, which is located in the same helix but at the opposite site of the arm (3 and 4 recession sites at the opposite edges of the arm). Therefore, these interlocking complementary sequences can efficiently bridge the side scaffold loops of these two adjacent DNA origami units.^40^ The combination of short hybridizing sequences and shape complementarity provides the needed specificity for correctly joining the units together, however, the interactions are still weak enough to allow rearrangements between the units and thus to help avoiding misaligned lattice formation.^41^

**Figure 2:**
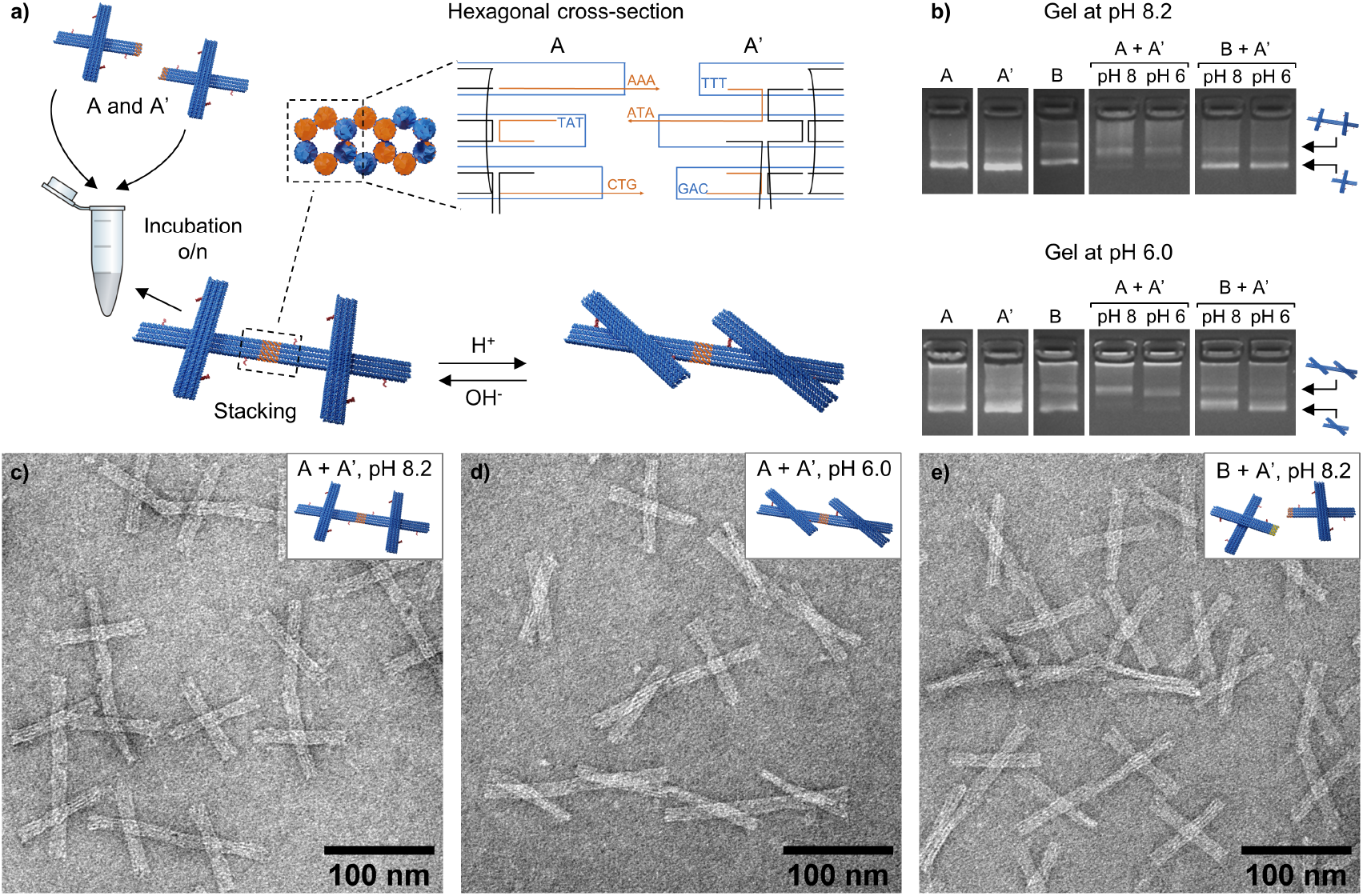
Formation of dynamic DNA origami dimers. (a) Dimers are formed by mixing equimolar amounts of both units (folded separately). The DNA origami units are selectively linked together by bridging the side scaffold loops with connector oligonucleotides. To connect the scaffold loops, seven of the connector oligonucleotides have a 3-nt overhang (in the 3’ end) complementary to the scaffold sequence on the opposite end of the arm. (b) Characterization of the dimer formation by AGE at pH 8.2 (top panel) and pH 6.0 (bottom panel). If not otherwise specified, the pH of the samples are 8.2 in the top gel and 6.0 in the bottom gel. TEM images of (c) dimers formed at pH 8.2 by combining A and A’ units (*c* = 5.7 nM) (d) the same dimer solution as in c) (A and A’ units, *c* = 5.4 nM) after the pH has been decreased to 6.0 with acetic acid. (e) a mixture of B and A’ units (*c* = 5.4 nM). These units do not have matching connector oligonucleotides and therefore no dimers are formed. The samples in TEM images are negatively stained with 2 % (w/v) uranyl formate.

To demonstrate the selectivity of the connector oligonucleotides, we, as a proof of concept, prepared different versions of pH-responsive DNA origami dimers (**Figures 2a, S9–S17**). The two units (marked with A and A’ if the connector oligonucleotides are in the bottom arm) were folded and purified from excess staple strands in separate batches, after which the dimers were formed by mixing equimolar amounts of both units. In order to prevent multimerization of the units, the interfaces of the arm ends not involved in dimer formation were poly-T-passivated (8-nt long polythymine overhangs). AGE revealed that the band corresponding to the single units almost completely vanished in the dimer mixture, whereas another band with lower electrophoretic mobility appeared in the gel, indicating a successful dimerization (**Figures 2b, S9, S13**). Importantly, a control sample with mismatching units (unit B with connectors in top arm combined with unit A’) did not form any dimers, demonstrating that our strategy to connect the units is indeed highly selective. To further confirm that the two units interact with each other correctly, we used TEM to visualize the formed dimers. The TEM images of the dimers that were assembled at pH 8.2 show, as expected, perfectly aligned and well-defined DNA origami dimers with the arms open (**Figures 2c, S10, S11**). Furthermore, by adding acetic acid to this dimer solution, the arms of the dimer could be locked into the closed configuration (**Figures 2d, S10, S11**). Equally, the dimers could be formed from the units initially at the closed state at pH 6.0, after which the arms could be released again by increasing the pH with sodium hydroxide (**Figures S13–S15**). As indicated above, B and A’ units neither have the required shape complementarity nor the matching sequences, and therefore, only discrete, unconnected DNA origami units were observed in TEM (**Figures 2e, S12, S16**).

### Formation of 1D Arrays Using the DNA Origami Unit

To further explore the possibility of using the DNA origami unit for the construction of large-scale lattices, we formed 1D arrays using the DNA origami unit. To this end, we prepared a unit with the polymerizing connector oligonucleotides on the bottom arm (A and A’ interactions) and fully poly-T-passivated interfaces on the top arm. To avoid undesired multimerization and formation of kinetically trapped configurations during the folding, the unit was prepared without the connector oligonucleotides. The polymerization of the units into linear arrays was initiated in a subsequent step by adding connector oligonucleotides in 10-fold excess to units that were earlier purified from the excess staple strands used in folding (**Figure 3a**, step 1). Initially, the assembly was carried out in solution by incubating the sample mixture at room temperature for at least 24 h. Although we recognized correctly formed linear chains when imaging the sample by TEM (**Figures 3b, S18, S19**), the tendency of the sample to form highly entangled structures set limitations to the analysis of the chain formation.

**Figure 3:**
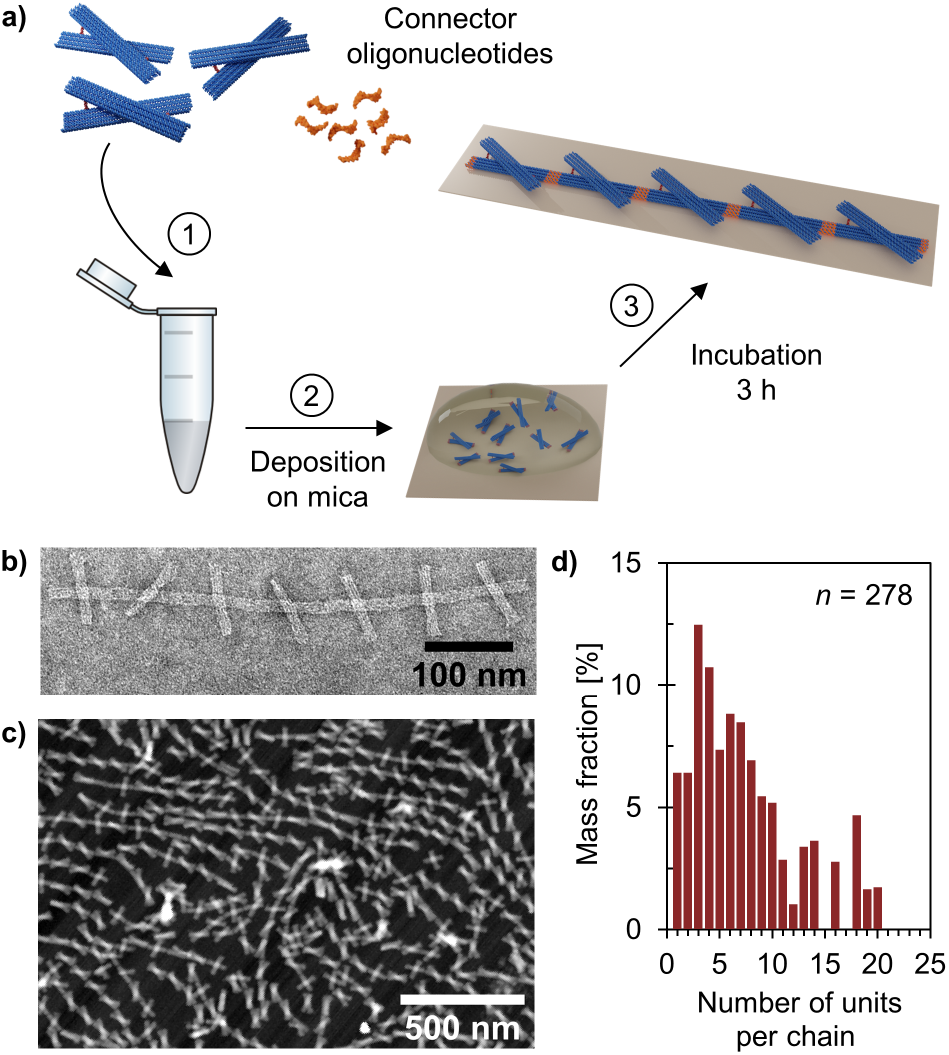
Formation of one-dimensional (1D) arrays using the DNA origami unit. (a) The polymerization of the units into chains is initiated by the addition of connector oligonucleotides. For the surface-mediated assembly, the mixture is immediately deposited onto a mica substrate. (b) TEM image of a negatively stained linear 1D array formed in solution at pH 8.2 (25 h incubation at room temperature, *c*_unit_ = 10.0 nM, but sample diluted 1:2 in 1 × FOB (1 × TAE, 20 mM MgCl_2_, 5 mM NaCl) before deposited onto the TEM grid). (c) Atomic force microscopy (AFM) image of DNA origami chains formed on a mica substrate at pH 6.0 (3 h incubation) (d) Observed chain length distribution for the 1D arrays assembled on a mica substrate at pH 6.0 (determined from AFM images).

As an alternative to the solution-phase formation, we also assembled the DNA origami chains on a mica substrate at the solid-liquid interface. The interface restricts the movement of the units to the 2D plane and may thus provide additional control of the lattice formation and growth.^33,42^ For the surface-assisted assembly, the units and the connector oligonucleotides were mixed together in a buffer supplemented with MgCl_2_ and NaCl, and immediately after that deposited onto a mica substrate (**Figure 3a**). Linear arrays were grown at both pH 6.0 (**Figure 3c, S20**) and pH 8.2 (**Figure S21, S22**), and in both cases discrete chains of various lengths were formed. 19% of the units assembled into >1 μm long chains (>11 units), while the majority of them formed chains of 3–10 units (pH 6.0, *n* = 278) (**Figure 3d**). This is also in line with the previously reported chain lengths for similar linear DNA origami arrays.^43^

### Assembly of pH-Responsive and Reconfigurable 2D DNA Origami Lattices

By introducing connector oligonucleotides on both the bottom and the top arms of the unit (A and A’ interactions as well as B and B’ interactions), we constructed a dynamic 2D lattice (**Figure 4a**). The two pH-sensitive conformations of the unit allow the lattice to adopt either an open or a closed configuration depending on the pH of the assembly solution. The 2D lattices were assembled directly onto the mica substrate by employing a previously established protocol^24^ that we further developed and optimized for our system. For successful formation of large hierarchical DNA origami assemblies on mica, the electrostatic interactions between the DNA origami and the surface have to be carefully controlled, which is usually accomplished by tuning the relative amounts of Na^+^ and Mg^2+^ in the assembly buffer.^42,44^ The divalent Mg^2+^ ions mediate the DNA origami adsorption onto mica by forming salt bridges, whereas the competitive Na^+^ ions weaken these interactions and enhance the DNA origami mobility on the surface. Depending on the assembly pH, we observed a clear difference in the DNA origami adsorption, which also affected the lattice growth. Therefore, we investigated the influence of the Mg^2+^ concentration on lattice formation on the mica substrate during 3 h by keeping Na^+^ concentration constant at 75 mM (**Figures 4b, S23, S24**). At pH 8.2, the optimum Mg^2+^ concentration was found to be 10 mM, which is well in agreement with previously optimized conditions.^45^ At pH 6.0, for one, the electrostatic interactions were noticeably weaker and a Mg^2+^ concentration of 12.5 mM was needed to obtain sufficient DNA origami adsorption for the subsequent lattice growth. The observed pH-dependent difference in the required Mg^2+^ concentration may be explained by the contribution of acetic acid that is used to decrease the pH of the assembly solution. Acetic acid is a known chelating agent that binds to Mg^2+^ ions and could thus reduce the availability of free Mg^2+^ ions in the solution. In addition, increasing the Mg^2+^ concentrations of the assembly solution beyond these optimized values results in high DNA origami adsorption and low DNA origami mobility on the surface, which remarkably decrease the lattice order.

**Figure 4:**
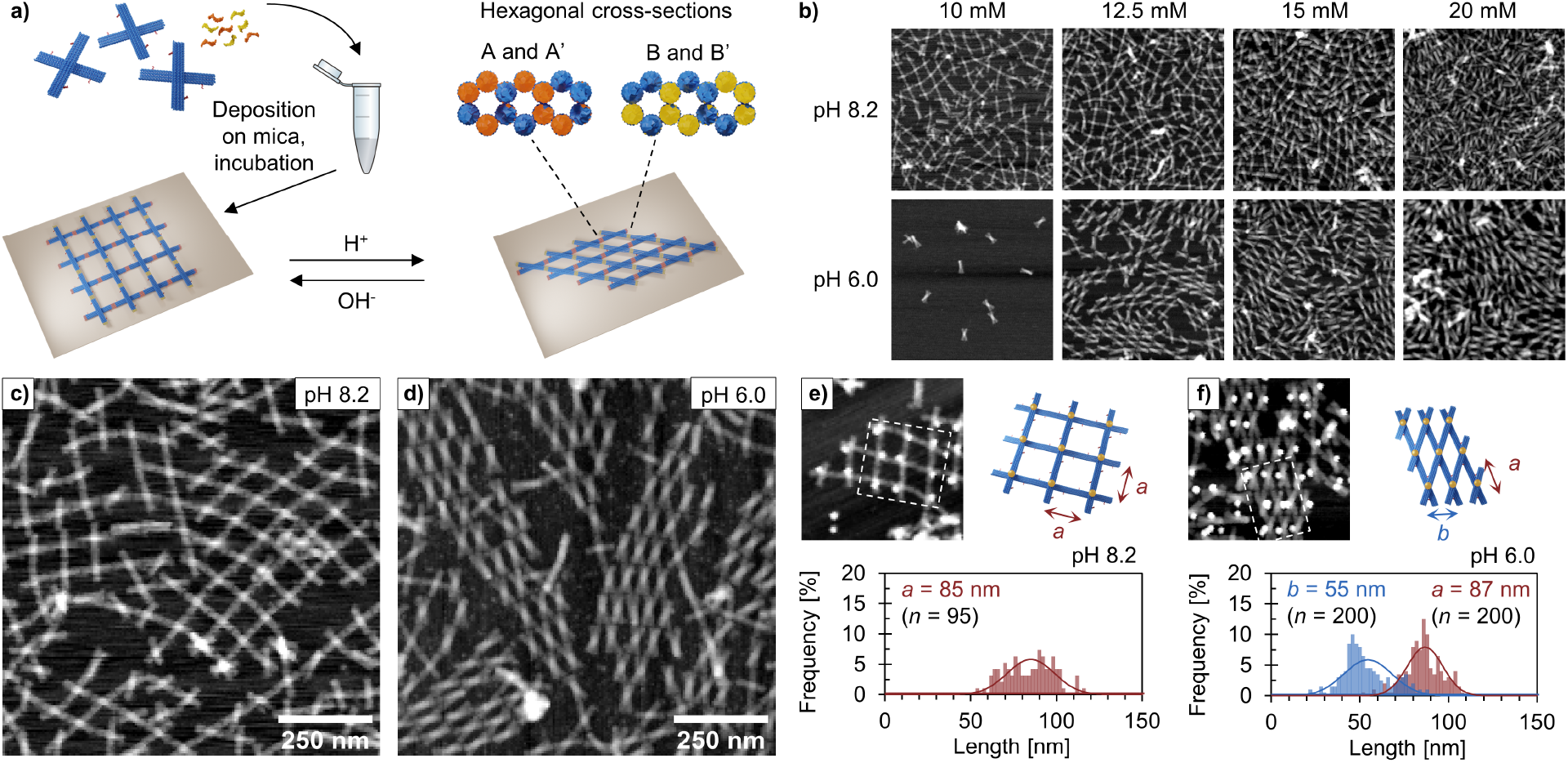
Assembly of pH-responsive and reconfigurable two-dimensional (2D) DNA origami lattices. (a) Connector oligonucleotides for both arms of the unit initiate the assembly of a 2D lattice on a mica substrate. The formed lattice could adapt either an open or a closed configuration depending on the pH of the surrounding solution. (b) AFM images (1 μm × 1 μm) of the 2D lattice formation at different Mg^2+^ concentrations. The Na^+^ concentration is kept constant at 75 mM and the assembly time is 3 h. AFM images of lattices assembled at (c) pH 8.2 and (d) pH 6.0 during 24 h. The DNA origami lattice could guide gold nanoparticles (AuNPs) into either (e) a square lattice at pH 8.2 or (f) a oblique lattice at pH 6.0. The top panel shows an AFM image of the AuNP lattice and the area marked with dotted lines is schematically presented next to the image. The bottom panel show the observed lattice constant distributions for the formed AuNP lattice (determined from the AFM images). The AuNP lattices are assembled during 3 h. In (c)–(f), the Mg^2+^ concentration is 10 mM for lattices at pH 8.2 and 12.5 mM for lattices at pH 6.0.

Depending on the assembly pH, the DNA origami lattice has two clearly distinguishable configurations (**Figures 4c,d**). At pH 8.2, the unit will adapt the open configuration and the formed lattice will be in an expanded state. At pH 6.0, on the other hand, the units are predominantly in the closed configuration and therefore a more compact lattice is formed. Nonetheless, in both cases the obtained lattice is polycrystalline and comprised of smaller crystalline domains of various sizes in close proximity to each other. The order and the size of the crystal domains correlate with the assembly duration, and therefore the crystal growth could be considerably improved by increasing the assembly time (**Figures S25–S28**). The crystal domains are generally also larger at pH 6.0, which could be explained by the enhanced rigidity of the unit when the arms are tied together and not able to rotate freely.

Thus far, most of the reported DNA origami-based frameworks have been static meaning that their lattice parameters have been fixed once they have been assembled. However, approaches allowing a stimuli-induced dynamic symmetry conversion after the assembly would be highly desirable. Therefore, we next studied whether our assembled pH-responsive and reconfigurable lattice could be readily expanded and squeezed by simply increasing or decreasing the pH. For these experiments, we first assembled the lattice on the mica surface for 3 h at pH 8.2 or 6.0, washed away weakly interacting and unbound assemblies, deposited a new buffer solution with lower/higher pH and incubated for additional 2 h (pH increase from 6.0 to 8.2) or 20 h (pH decrease from 8.2 to 6.0). When the pH was increased from 6.0 to to 8.2, a clear change from the closed state toward the open lattice configuration was observed (**Figure S30**), indicating that the formed lattice is rather mobile on the surface. Closing of the lattice after assembly, (pH decrease from 8.2 to 6.0), for one, required much longer time, and the overall change in the lattice configuration was not as pronounced as in the case of opening the lattice (**Figure S31**).

### Assembly of DNA-Templated, pH-Responsive, and Reconfigurable 2D AuNP Lattices

It is known that spatially well-defined arrangements of metal nanoparticles possess intriguing optical, plasmonic, electronic, and magnetic properties,^2^ but fabrication of highly ordered dynamic nanoparticle lattices is rather challenging. As already mentioned, programmable and modular DNA-based structures are suitable templates for guiding nanoparticles into complex, mostly static lattices using either DNA hybridization^21,25,26^ or electrostatic interactions.^46^ In order to demonstrate that our pH-sensitive lattice could be used as a template to create reconfigurable nanoparticle lattices, we modified the DNA origami unit by adding an anchoring site for an oligonucleotide-coated gold nanoparticle (AuNP, 10 nm in diameter) in the middle of the unit (**Figure S7**). AFM images of the prepared lattices show, as expected, two distinct lattice configurations depending on the assembly pH; a 2D square lattice at pH 8.2 (**Figure 4e, S32**) and a 2D oblique lattice at pH 6.0 (**Figure 4f, S33**). Furthermore, the average lattice constants determined by AFM are *a* = 85 ± 14 nm for the square lattice and *a* = 87 ± 10 nm, *b* = 55 ± 14 nm for the oblique lattice. The highest frequency was observed for *a* = 90–92 nm for the square lattice and *a* = 86–88 nm and *b* = 44–46 nm for the oblique lattice. The DNA origami unit is rather flexible at pH 8.2 and taking that into account, the observed lattice constants are well in agreement with the theoretical ones (*a* = 86 nm (both for square and oblique lattices) and *b* = 45 nm, assuming a vertex angle of 30°)

## Conclusions

In this work, we have presented a strategy for constructing pH-responsive and dynamically reconfigurable lattices using DNA origami as the building block. The pH-responsiveness of the lattice is achieved by equipping the arms of the pliers-like, lattice-forming DNA origami unit with pH latches that form Hoogsteen-type of triplexes in low pH. Therefore the unit could rapidly switch between an open "+"-shaped and a closed "X"-shaped configuration upon a pH change. Nevertheless, the high level of programmability of the DNA origami would equally enable other stimuli-responsive elements, such as e.g. photoresponsive molecules^47^ and thermoresponsive polymers,^48^ to be implemented into the basic building block of the lattice, thus allowing reconfigurable lattices that undergo conformational changes in response to different external stimuli. Furthermore, the unprecedented addressability of DNA origami allows not only AuNPs (as demonstrated here) but also a wide variety of other compounds to be precisely positioned onto DNA origami frameworks. Therefore, we believe that our demonstrated system as well as other recently reported reconfigurable DNA-based lattices^28–30,49^ will play an important role for the development of more sophisticated stimuli-responsive and functional materials in future.

## Methods

### Design and Preparation of the pH-Responsive DNA Origami Unit

The pH-responsive DNA origami unit was designed on a honeycomb lattice using caDNAno v 2.2.0,^50^ and its three-dimensional shape was predicted using the CanDo software. ^51,52^ The caDNAno design for the unit is shown in **Figures S35–S36** and the staple strands needed for the different versions of the unit are listed in **Tables S2–S9**.

The DNA origami units were folded in a one-pot reaction in either 50 μL or 100 μL quantities by mixing the circular p7249 scaffold (final concentration of 20 nM) with 7.5× excess of staple strands in a folding buffer (FOB) containing 1× Tris-acetate-EDTA (TAE) buffer, 20 mM MgCl_2_ and 5 mM NaCl. The folding reaction mixture was thermally annealed from 75 °C to 27 °C degree in a ProFlex PCR system or a G-storm G1 Thermal Cycler using the following annealing program: (1) Cooling from 75 °C to 70 °C at a rate of −0.2 °C / 8 s; (2) Cooling from 70 °C to 60 °C at a rate of −0.1 °C / 8 s; (3) Cooling from 60 °C to 27 °C at a rate of −0.1 °C / 2 min; (4) Cooled down to 20 °C or 12 °C and stored at this temperature until the program was manually stopped.

The excess staple strands were removed from the folded DNA origami structures using a polyethylene glycol (PEG) precipitation method.^53^ First, the DNA origami solution was diluted four-fold with 1× FOB, after which the solution was thoroughly mixed 1:1 with PEG precipitation buffer (15% (w/v) PEG 8000, 1× TAE, 505 mM NaCl). The mixture was centrifuged at 14 000 *g* for 30 min at room temperature using and Eppendorf 5424R microcentrifuge, the supernatant was carefully removed and the DNA origami pellet resuspendended in 1× FOB (either at pH 8.2 or 6.0) to the original reaction volume. To dissolve the pellet, the DNA origami solution was incubated at 30 °C overnight under continuous shaking at 600 rpm using an Eppendorf Thermomixer C. The DNA origami concentration was estimated as described in Section 2.1 in the Supporting Information.

### Dimer Formation

The DNA origami dimers were formed by mixing equimolar amounts of PEG-purified DNA origami units (to a final concentration of 5.7 nM) in 1× FOB (either at pH 8.2 or 6.0). To allow the formation of dimers, the samples were incubated at room temperature for at least 22 h. The pH of the dimer solution was decreased / increased by adding 1.5 μL of 0.5 M acetic acid or 0.5 M sodium hydroxide to 30 μL of dimer solution and incubating at room temperature for at least additional 23 h.

### 1D Array Formation in Solution

For the assembly of DNA origami chains in solution, PEG-purified DNA origami units (final concentrations of 2.0, 5.0 or 10.0 nM) were mixed with 10-fold excess of connector oligonucleotides in 1× FOB (either at pH 8.2 or 6.0). To allow polymerization, the samples were incubated at room temperature for at least 24 h.

### 1D and 2D Lattice Assembly on Mica

The lattice assembly on mica was mainly carried out following a procedure previously described by Xin et al.^24^ For the deposition, PEG purified DNA origami units (final concentration of 2.0 nM) were mixed with 10-fold excess of connector oligonucleotides in a buffer (at either pH 8.2 or 6.0) containing 1× TAE supplemented with MgCl_2_ (10–20 mM depending on the sample) and 75 mM NaCl. 120 μL of the DNA origami sample mixture was evenly deposited onto a freshly cleaved mica surface (15 mm × 15 mm, grade V1, Electron Microscopy Sciences) and incubated covered at room temperature for 3–24 h. After the incubation, the mica surface was rinsed 5 times with 100 μL of 1× TAE supplemented with MgCl_2_ (same MgCl_2_ concentration and pH as in the sample solution). Immediately after the washing step, 120 μL of 1× TAE containing 10 mM NiCl_2_ was deposited on the mica surface and incubated covered for 1 h. After the incubation, the mica surface was rinsed 6 times with 100 μL of deionized water, after which the sample was dried thoroughly using a nitrogen gas stream.

### Preparation of DNA-Functionalized AuNPs and AuNP-Conjugated DNA Origami Units

The DNA-functionalized AuNPs were mainly prepared as described previously.^37^ If not stated otherwise, all the steps of the DNA-functionalization of the AuNPs were carried out at 40 °C under constant shaking at 600 rpm using an Eppendorf Thermomixer C. First, 80 μL of citrate-stabilized AuNPs (10 nm in diameter, upconcentrated to 50 nM) was incubated with 1.6 μL of 1% (w/v) sodium dodecyl sulfate (SDS) solution for 20 min. Next, 16 μL of thiolated oligonucleotides (*c* = 100 μM, see **Table S7** for the sequence) was added and the mixture was incubated for additional 30 min, after which a salt-aging process was carried out. First, 0.8 μL of 2.5 M NaCl was added every 5 min (6 times), followed by 1.6 μL of 2.5 M NaCl every 5 min (6 times), 3.2 μL of 2.5 M NaCl every 5 min (5 times) and 2.0 μL of 2.5 M NaCl (once). After the salt-aging, 120 μL of 1× FOB (1× TAE buffer, 20 mM MgCl_2_, 5 mM NaCl) supplemented with 0.02% (w/v) SDS was added and the mixture incubated for 60 minutes. The temperature was decreased to 20 °C after which the incubation continued overnight (constant shaking at 600 rpm).

Before the conjugation to the DNA origami unit, the DNA-functionalized AuNPs were purified from excesss thiolated oligonucleotides using spin-filtration at room temperature (Amicon Ultra 100 kDa MWCO centrifugal filter, EMD Millipore). The filter was washed with 200 μL of 1× FOB with 0.02% (w/v) SDS (14 000 *g*, 5 min) before use. DNA-functionalized AuNPs (260–400 μL per addition, in total 1260 μL) were added to the filter unit and after each addition, the unit was centrifuged at 14 000 *g* for 10 min using an Eppendorf micro-centrifuge 5424R. Finally, the DNA-functionalized AuNPs were washed 3 times by adding 200 μL of 1× FOB with 0.02% (w/v) SDS and centrifuging at 14 000 *g* for 10 min. The DNA-functionalized AuNPs were recovered by inverting the filter unit and centrifuging at 2000 *g* for 2.5 min.

The DNA origami unit has a position for AuNP attachment in the middle of the unit (see **Table S7** for the sequences of the attachment strands). For the conjugation, 7.5× excess of DNA-functionalized AuNPs were mixed with PEG purified DNA origami units (final concentration of 7.5 nM in the conjugation mixture) in 1× FOB supplemented with 0.01% (w/v) SDS. To increase the attachment yield, the mixture was thermally annealed from 40 to 20 °C at a rate of −0.1 °C/min using a ProFlex PCR system.

Before the lattice assembly on mica, PEG precipitation^54^ was used to remove the SDS and some of the free AuNPs from the solution with AuNP-conjugated DNA origami units. 50 μL of AuNP-conjugated DNA origami units were mixed with 12.5 μL of PEG precipitation buffer (17.5% (w/v) PEG 8000, 1× TAE, 10 mM MgCl_2_, 500 mM NaCl). The mixture was incubated at 4 °C for 10 min before centrifuged at 12 600 *g* at 4 °C for 30 min. The supernatant was carefully removed after which the pellet was resuspended in 50 μL of 1× FOB at either pH 8.2 or 6.0. The solution was incubated at room temperature overnight before deposited on mica. The lattice assembly was done as described above and the composition of the used buffer was the same, but the DNA origami concentration was slightly higher (2.0–3.0 nM, calculated based on the concentration in the conjugation step, assuming no loss during the PEG precipitation).

### Agarose Gel Electrophoresis

Agarose gel electrophoresis was used to analyze the folding of the DNA origami unit as well as the formation of DNA origami dimers. A 2% (w/v) agarose gel was prepared in 1× TAE buffer containing 11 mM MgCl_2_ for the gel at pH 8.2, whereas the 2% (w/v) agarose gel was prepared in 45 mM MES and 25 mM Tris containing 11 mM MgCl_2_ for the gel at pH 6.0. Both gels were stained with ethidium bromide (final concentration of 0.46 μg mL^−1^). Depending on the sample and the type of gel, the sample volume was 10–18 μL and the DNA origami concentration 11.1/15.0 nM (DNA origami units) or 5.4/5.7 nM (DNA origami dimers). A gel loading dye solution was added to the samples at a ratio of 1:5 before loading the samples in the gel pockets. The gel was run for 45 minutes at a constant voltage of 95 V using a BioRad Wide Mini-Sub Cell GT System and a BioRad PowerPac Basic power supply while keeping the gel electrophoresis chamber on an ice bath. For the gel at pH 8.2, the running buffer was 1× TAE buffer supplemented with 11 mM MgCl_2_, whereas 45 mM MES and 25 mM Tris supplemented with 11 mM MgCl_2_ was used as running buffer for the gel at pH 6.0. After the run, the gel was visualized by ultraviolet light using a BioRad Gel Doc XR+ documentation system.

### Transmission Electron Microscopy

The TEM samples were prepared on glow-charged (20 s oxygen plasma flash) Formvar carbon-coated copper grids (FCF400-Cu, Electron Microscopy Sciences) according to the protocol previously described by Castro et al. ^51^ 3 μL of DNA origami solution (*c* = 7.5 nM for DNA origami units, *c* = 5.4/5.7 nM for DNA origami dimers and *c* = 2.0–5.0 nM) was applied onto the carbon-coated side of the grid and incubated for 3 min before excess sample solution was blotted away with filter paper. After that, the sample was negatively stained with 2% (w/v) aqueous uranyl formate solution containing 25 mM NaOH (added to increase the pH of the stain solution) in two subsequent steps. First, the sample was immersed into a 5-μL droplet of stain solution, after which the stain was immediately removed using filter paper. Next, the sample was immersed into a 20-μL droplet of stain solution for 45 s before the solution was blotted away with a filter paper. The samples were left to dry under ambient conditions for at least 15 minutes before imaging. All TEM images were obtained using a FEI Tecnai 12 Bio-Twin electron microscope operated at an acceleration voltage of 120 kV. The images were processed and analyzed (vertex angle measurements) using ImageJ.

### Atomic Force Microscopy

The atomic foce microscopy (AFM) images were obtained using a Dimension Icon AFM (Bruker). The samples were imaged in air using ScanAsyst in Air Mode and ScanAsyst-Air probes (Bruker). The AFM images were recorded with a resolution of 512 pxl × 512 pxl and a scan rate of 0.5 Hz or 0.75 Hz depending on the scan size (5 μm × 5 μm, 3 μm × 3 μm, or 2 μm × 2 μm). The images were processed (row alignment, correction of horizontal scars and height scale adjustment) using NanoScope Analysis (v. 1.90, Bruker) and/or Gwyddion open source software (v. 2.58).^55^

## Supporting Information Available

The following files are available free of charge.

- Supporting Information: Material sources; Supplementary methods including molar extinction coefficients for different versions of the DNA origami unit; Additional characterization data (AGE, TEM, AFM) of the DNA origami unit; AGE and TEM images demonstrating the selective assembly of DNA origami dimers; Additional TEM and AFM images of 1D array formation both in solution and on mica; AFM images of 2D lattice formation on mica using different parameters (pH, MgCl_2_, assembly time, unit type); AFM images demonstrating the pH-responsiveness of already assembled lattices, Additional AFM images of AuNP lattices; DNA origami unit design and full list of staple strands for the DNA origami units (PDF)

## Notes

The authors declare no competing financial interest.

## Acknowledgement

This work was supported by the Academy of Finland (project numbers 308578 and 314671), European Research Council (ERC) and ERA Chair MATTER under the European Union’s Horizon 2020 research and innovation programme (grant agreement numbers 101002258 and 856705, respectively), Aalto University School of Chemical Engineering, Victoriastiftelsen, Finnish Cultural Foundation (Maili Autio Fund), Jane and Aatos Erkko Foundation, Sigrid Jusélius Foundation, Emil Aaltonen Foundation, and Vilho, Yrjö and Kalle Väisälä Foundation of the Finnish Academy of Science and Letters. The work was carried out under the Academy of Finland Centers of Excellence Programme (2022-2029) in Life-Inspired Hybrid Materials (LIBER), project number 346110. The authors thank E. Kaipia for assisting in preparing the DNA-functionalized AuNPs, J. V. I. Timonen for technical assistance, as well as A. Keller and H. Ijäs for fruitful discussions. We also acknowledge the provision of facilities and technical support by Aalto University Bioeconomy Facilities, OtaNano - Nanomicroscopy Center (Aalto-NMC) and Micronova Nanofabrication Center.

## TOC Graphic

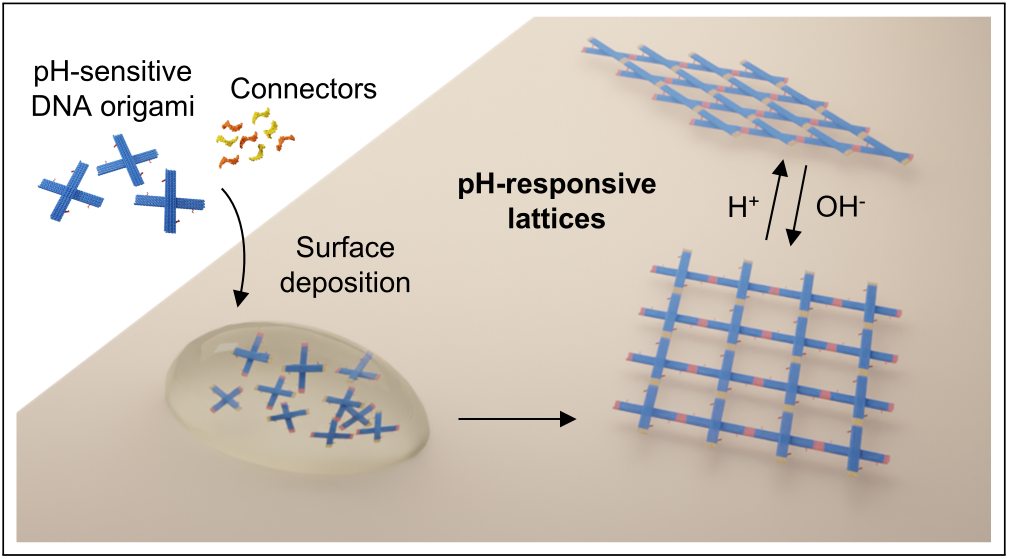

