## Supporting Information for "Reconfigurable pH-Responsive DNA Origami Lattices"

### Contents


| | | Page |
|---|---|---|
| 1 | **Materials** | **S3** |
| 2 | **Supplementary Methods** | **S4** |
| | 2.1 DNA origami concentration estimation | S4 |
| | 2.2 AFM imaging of the DNA origami unit | S4 |
| | 2.3 pH responsiveness of assembled 2D DNA origami lattice | S4 |
| 3 | **pH-Responsive DNA Origami Unit** | **S6** |
| 4 | **pH-Responsive DNA Origami Dimers** | **S14** |
| 5 | **pH-Responsive 1D DNA Origami Arrays** | **S23** |
| 6 | **pH-Responsive and Reconfigurable 2D DNA Origami Lattices** | **S28** |
| 7 | **DNA-Templated, pH-Responsive, and Reconfigurable 2D AuNP Lattices** | **S37** |
| 8 | **DNA origami Unit Design** | **S40** |
| 9 | **References** | **S49** |

#### 1. Materials

All chemicals were purchased from commercial suppliers and used as received unless otherwise noted. For preparation of the DNA origami unit, the circular single-stranded p7249 scaffold ($c$ = 100 nM) was purchased from Tilibit Nanosystems and the single-stranded staple strands from Integrated DNA Technologies. 50× TAE buffer (2 M tris(hydroxymethyl)aminomethane (Tris), 1 M acetic acid, 50 mM ethylenediaminetetraacetic acid (EDTA), pH 8.4) was purchased from Thermo Fischer Scientific. For the agarose gel electrophoresis, the ethidium bromide and gel loading dye solution were purchased from Sigma Aldrich, whereas the agarose was purchased from Meridian Bioscience. The gel loading dye solution (0.25% bromophenol blue, 0.25% xylene cyanol, 40% sucrose) was diluted 1:30 in 40% (w/v) sucrose before used. The uranyl formate for staining the TEM samples were purchased from Electron Microscopy Sciences. The citrate stabilized gold nanoparticles (10 nm in diameter) were purchased from either Sigma Aldrich or nanoComposix. For all of the experiments, deionized water (Milli-Q grade) was used.

#### 2. Supplementary Methods

##### 2.1. DNA origami concentration estimation

The DNA origami concentration after PEG purification was estimated from the DNA origami absorbance at 260 nm using Beer-Lambert law:

$$A_{260} = \epsilon_{260} c l \quad \textbf{(S1)}$$

where $A_{260}$ is the absorbance at a wavelength of 260 nm, $\epsilon_{260}$ is the approximated molar extinction coefficient of the DNA origami structure at 260 nm and $l$ is the path lenght through the solution in centimeters (0.05 cm in the used set-up). The absorbance at 260 nm was measured using a BioTek Eon Microplate Spectrophotometer and a Take3 microvolume plate. A sample size of 2 µL was used for the measurements and the concentration was obtained as the average of three measurements. The molar extinction extinction coefficients, $\epsilon_{260}$, where calculated as

$$\epsilon_{260} = 6700 \times N_{\mathrm{ds}} + 10000 \times N_{\mathrm{ss}} \quad \textbf{(S2)}$$

where $N_{\mathrm{ds}}$ is the number of hybridized nucleotides and $N_{\mathrm{ss}}$ is the number of non-hybridized, single-stranded nucleotides in the DNA origami unit.(1)

**Table S1.** Molar extinction coefficients for different versions of the DNA origami unit.

| DNA origami unit | Extinction coefficient ($M^{-1}cm^{-1}$) |
|---|---|
| DNA origami unit with pH latches, polyT-passivated | $1.02 \times 10^8$ |
| Permanently closed DNA origami unit, polyT-passivated | $1.02 \times 10^8$ |
| Permanently open DNA origami unit, polyT-passivated | $1.01 \times 10^8$ |
| DNA origami unit with pH latches for dimer formation | $1.01 \times 10^8$ |
| DNA origami unit with pH latches, polyT-passivated on the top arm (used for 1D arrays) | $0.98 \times 10^8$ |
| DNA origami unit with pH latches and scaffold loops (used for 2D lattices) | $0.94 \times 10^8$ |
| DNA origami unit with pH latches, AuNP attachment strands and scaffold loops (used for 2D lattices) | $0.95 \times 10^8$ |
| Permanently closed DNA origami unit with scaffold loops (used for 2D lattices) | $0.94 \times 10^8$ |
| Permanently closed DNA origami unit with AuNP attachment strands and scaffold loops (used for 2D lattices) | $0.94 \times 10^8$ |

##### 2.2. AFM imaging of the DNA origami unit

10 µL of poly-T passivated DNA origami units with pH latches (PEG-purified, $c$ = 1 nM for the sample at pH 8.2 and $c$ = 1 nM for the sample at pH 6.0) was deposited onto a freshly cleaved mica surface (15 mm × 15 mm, grade V1, Electron Microscopy Sciences) and incubated covered at room temperature for 1 min. After the incubation, the mica surface was rinsed 3 times with 100 µL of deionized water, after which the sample was dried thorouhly using a nitrogen gas stream.

The atomic foce microscopy (AFM) images were obtained using a Dimension Icon AFM (Bruker). The samples were imaged in air using ScanAsyst in Air Mode and ScanAsyst-Air probes (Bruker). The AFM images were recorded with a resolution of 512 pxl × 512 pxl, a scan rate of 0.5 Hz and a scan size of 2 µm × 2 µm. The images were processed using Gwyddion open source software (v. 2.58). (2)

##### 2.3. pH responsiveness of assembled 2D DNA origami lattice

In order to demonstrate that the lattice is pH-responsive and reconfigurable also after the initial assembly, the following approach was used. PEG-purified DNA origami units (final concentration of 2.0 nM) were mixed with 10-fold excess of connector oligonucleotides in 1× TAE supplemented with 10 mM $MgCl_2$ and 75 mM NaCl at pH 8.2 (lattice assembled at pH 8.2) or in 1× TAE supplemented with 12.5 mM $MgCl_2$ and 75 mM NaCl at pH 6.0 (lattice assembled at pH 6.0). 120 µL of the DNA origami sample mixture was evenly deposited onto a freshly cleaved mica surface (15 mm × 15 mm, grade V1, Electron Microscopy Sciences) and incubated covered at room temperature for 3 h. After the incubation, the mica surface was rinsed 5 times with 100 µL of 1× TAE, 10 mM $MgCl_2$ at pH 8.2 (for the lattice assembled at pH 8.2) or 100 µL of 1× TAE, 12.5 mM $MgCl_2$ at pH 6.0 (for the lattice assembled at pH 6.0). In order to decrease the pH, 120 µL of 1× TAE, 12.5 mM $MgCl_2$ and 75 mM NaCl at pH 6.0 was deposited on the lattice assembled at pH 8.2 and incubated covered for 20 h. Similarly, in order to increase the pH, 120 µL of 1× TAE, 10 mM $MgCl_2$ and 75 mM NaCl at pH 8.2 was deposited on the lattice assembled at pH 6.0 and incubated covered for 2 h. After the incubation, the mica surface was rinsed 5 times with 100

$\mu$L of 1$\times$ TAE, 12.5 mM $MgCl_2$ at pH 6.0 (for the lattice changed to pH 6.0) or 100 $\mu$L of 1$\times$ TAE, 10 mM $MgCl_2$ at pH 8.2 (for the lattice changed to pH 8.2). Immediately after the second washing step, 120 $\mu$L of 1$\times$ TAE, 10 mM $NiCl_2$ was deposited on the mica surface and incubated covered for 1 h. After the incubation, the mica surface was rinsed 6 times with 100 $\mu$L of deionized water, after which the sample was dried thoroughly using a nitrogen gas stream.

The atomic foce microscopy (AFM) images were obtained using a Dimension Icon AFM (Bruker). The samples were imaged in air using ScanAsyst in Air Mode and ScanAsyst-Air probes (Bruker). The AFM images were recorded with a resolution of 512 pxl $\times$ 512 pxl, a scan rate of 0.75 Hz and a scan size of 3 $\mu$m $\times$ 3 $\mu$m. The images were processed using Gwyddion open source software (v. 2.58). (2)

#### 3. pH-Responsive DNA Origami Unit

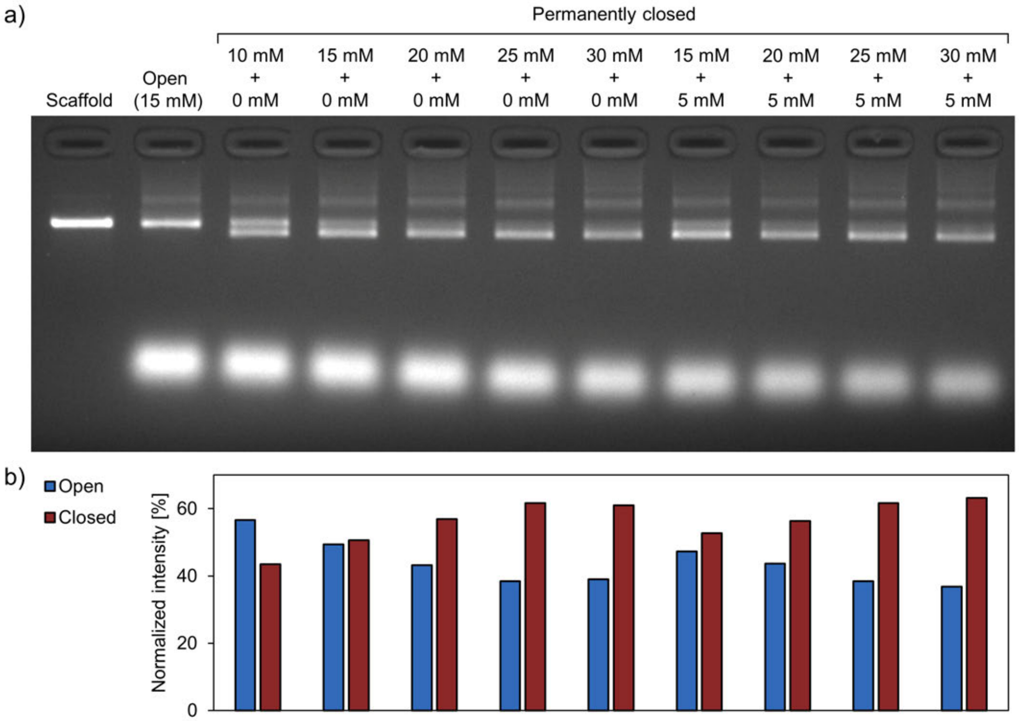


**Figure S1.** Characterization of permanently closed DNA origami units folded at different $MgCl_2$ and NaCl concentrations. (a) Agarose gel electrophoresis of folded, unpurified structures. The top concentration is the $MgCl_2$ concentration and the bottom the NaCl concentration. The DNA origami concentration in the gel is 11.1 nM and the gel was run at pH 8.2. (b) Ethidium bromide intensity of the bands corresponding to the closed and open configurations. The total intensity of the two bands corresponding to the open and the closed configuration were always set to 100 %.

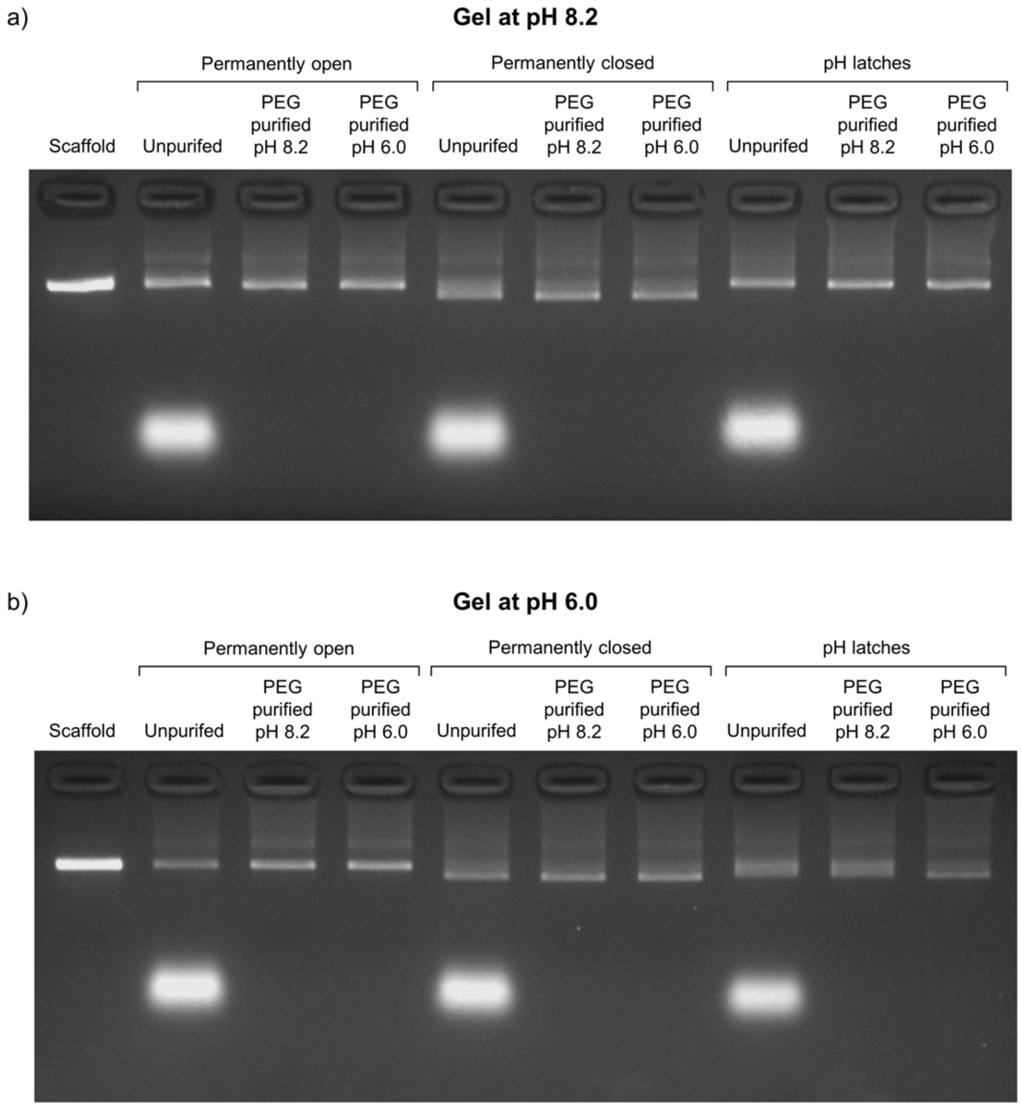


**Figure S2.** Characterization of the DNA origami units by agarose gel electrophoresis at (a) pH 8.2 and (b) pH 6.0. Selected parts of the gels are also shown in the article in Figure 1b. The DNA origami concentration in the gels is 15.0 nM.

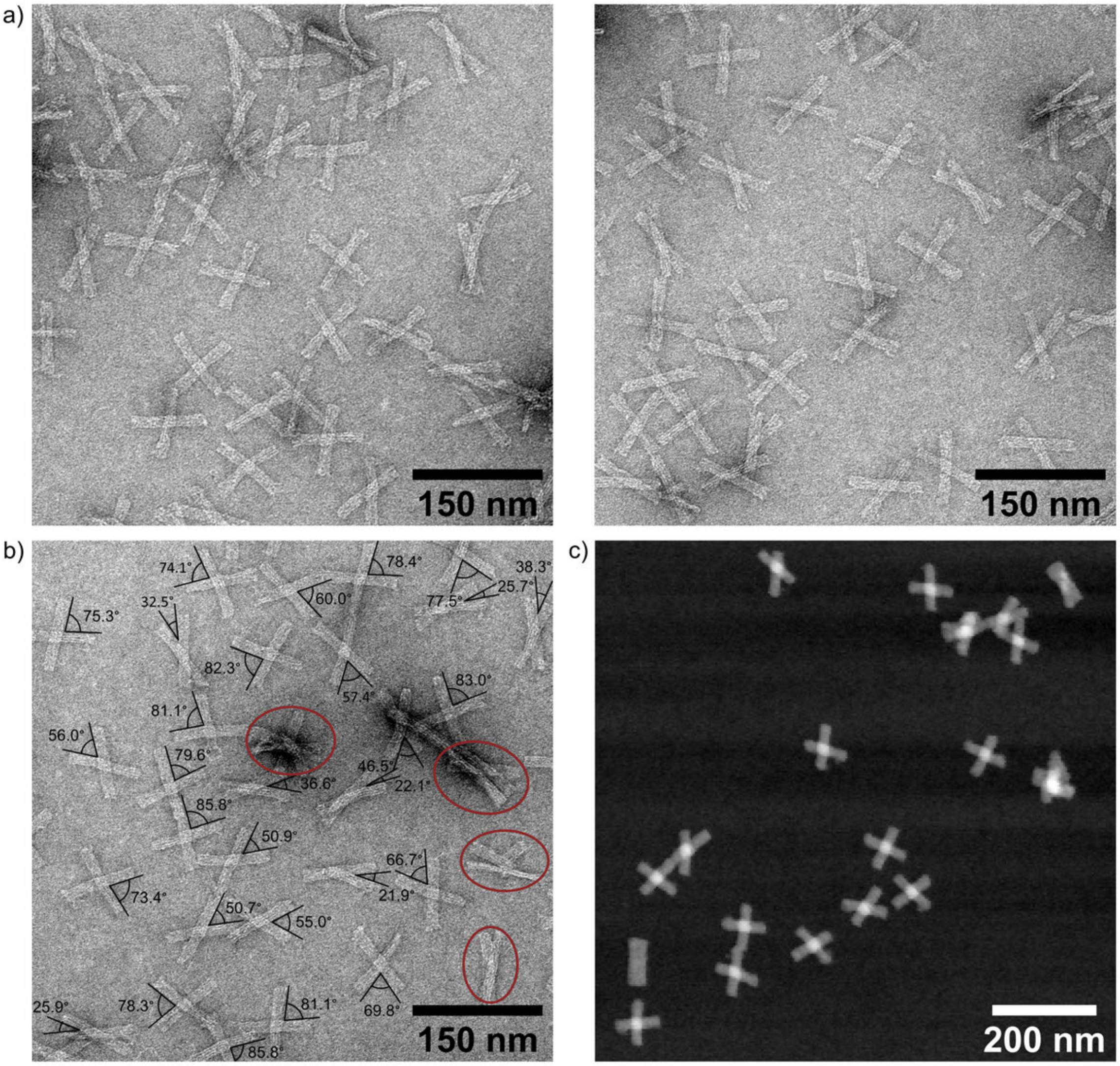


**Figure S3.** Characterization of the DNA origami unit with pH latches (poly-T passivated) at pH 8.2. (a) TEM images of the unit ($c$ = 5.0 nM) in 1× FOB (1× TAE, 20 mM $MgCl_2$, 5 mM NaCl) at pH 8.2. (b) Examples of determined angles, $\alpha$, between the two arms of the unit. The encircled structures were excluded from the analysis. (c) AFM image of the unit ($c$ = 1 nM) in 1× FOB at pH 8.2. The TEM images are negatively stained with uranyl formate (2 % (w/v)).

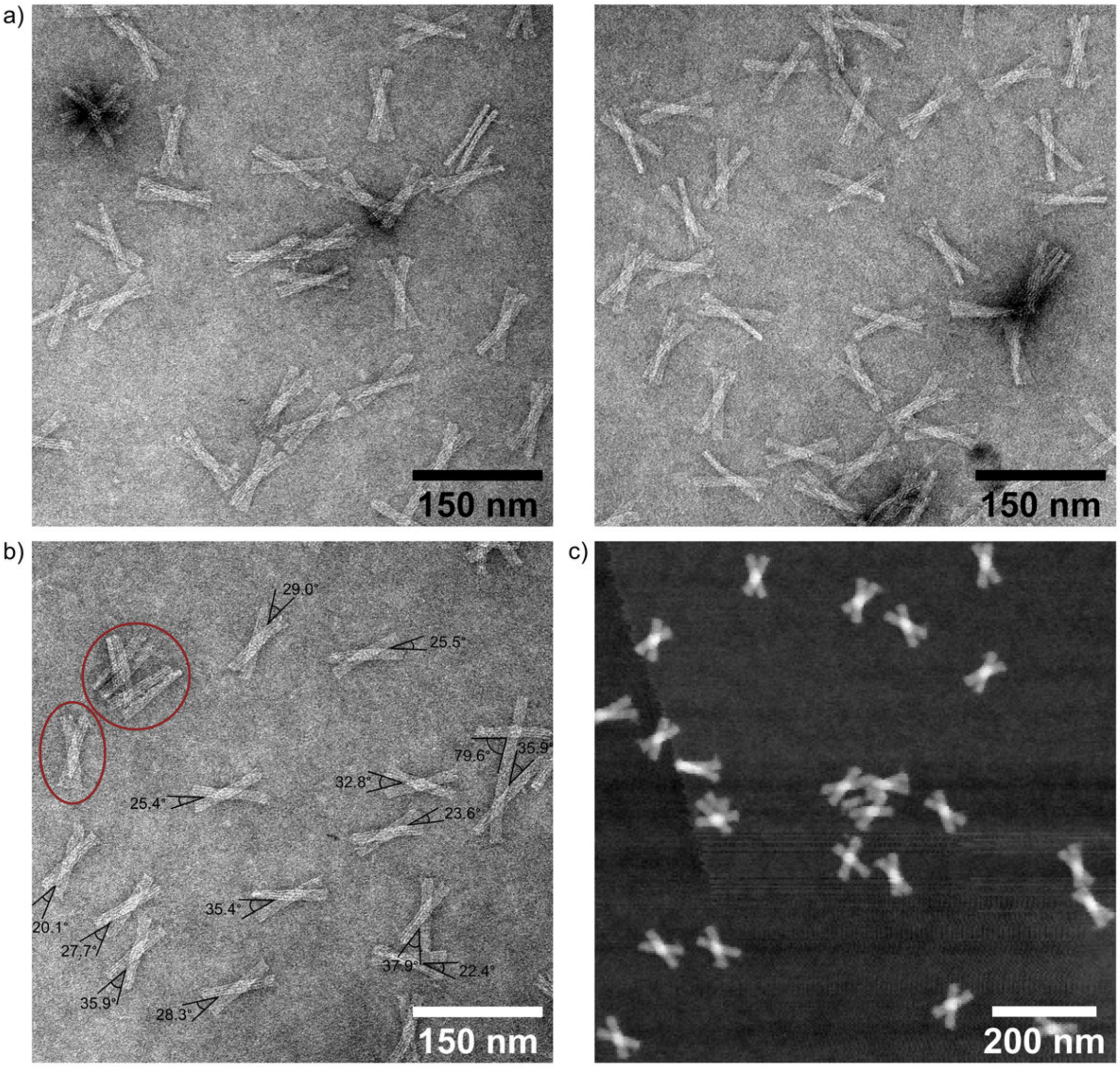


**Figure S4.** Characterization of the DNA origami unit with pH latches (poly-T passivated) at pH 6.0. (a) TEM images of the unit ($c$ = 5.0 nM) in 1× FOB (1× TAE, 20 mM $MgCl_2$, 5 mM NaCl) at pH 6.0. (b) Examples of determined angles, $\alpha$, between the two arms of the unit. The encircled structures were excluded from the analysis. (c) AFM image of the unit ($c$ = 2 nM) in 1× FOB at pH 6.0. The TEM images are negatively stained with uranyl formate (2 % (w/v)).

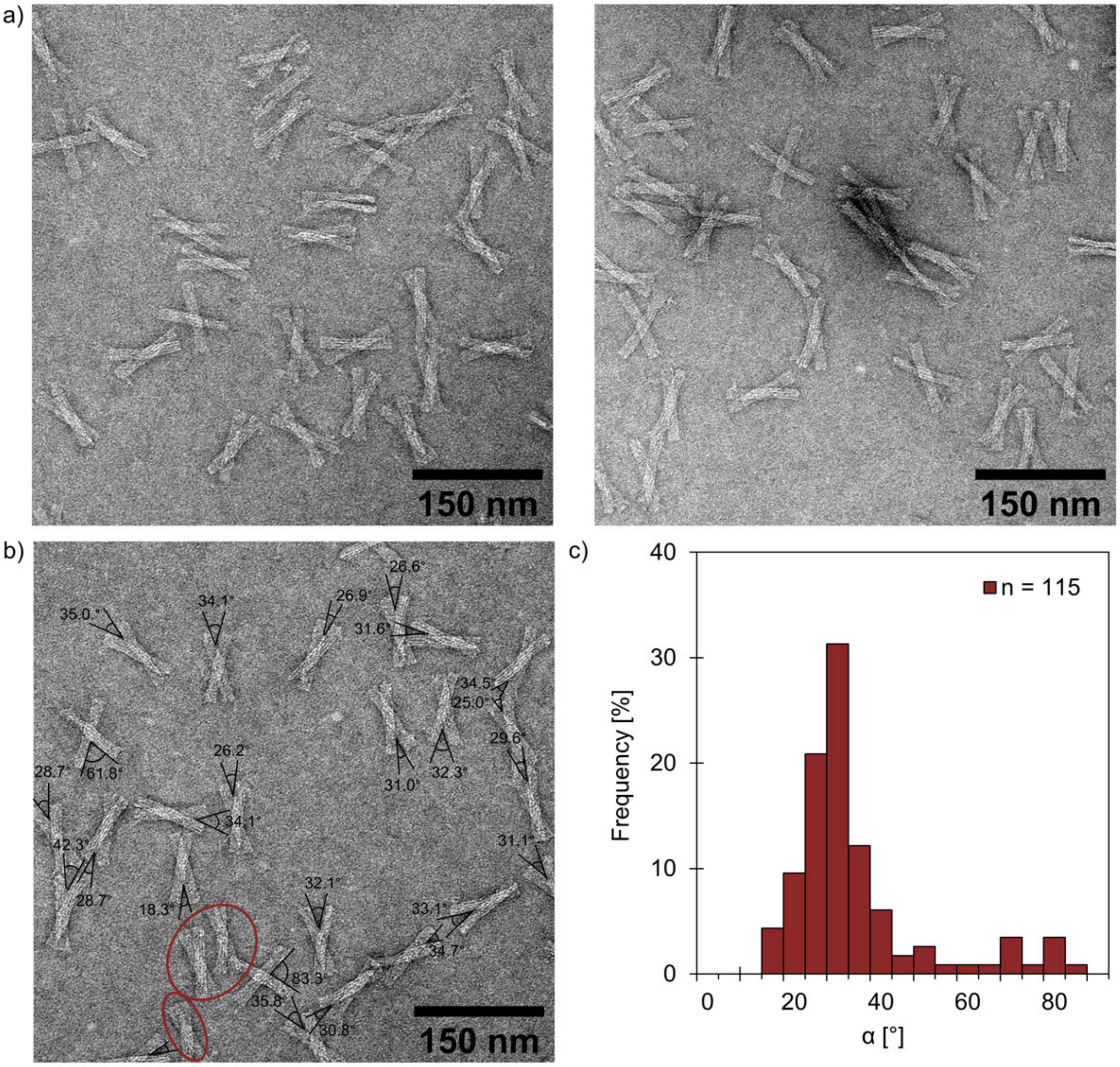


**Figure S5.** Characterization of the permanently closed DNA origami unit (poly-T passivated) at pH 8.2. (a) TEM images of the unit ($c$ = 5.0 nM) in $1\times$ FOB ($1\times$ TAE, 20 mM $MgCl_2$, 5 mM NaCl) at pH 6.0. (b) Examples of determined angles, $\alpha$, between the two arms of the unit. The encircled structures were excluded from the analysis. (c) The distribution of the angle, $\alpha$, between the two arms of the unit. The number of individual structures analyzed for the sample is $n$ = 115. The TEM images are negatively stained with uranyl formate (2 % (w/v)).

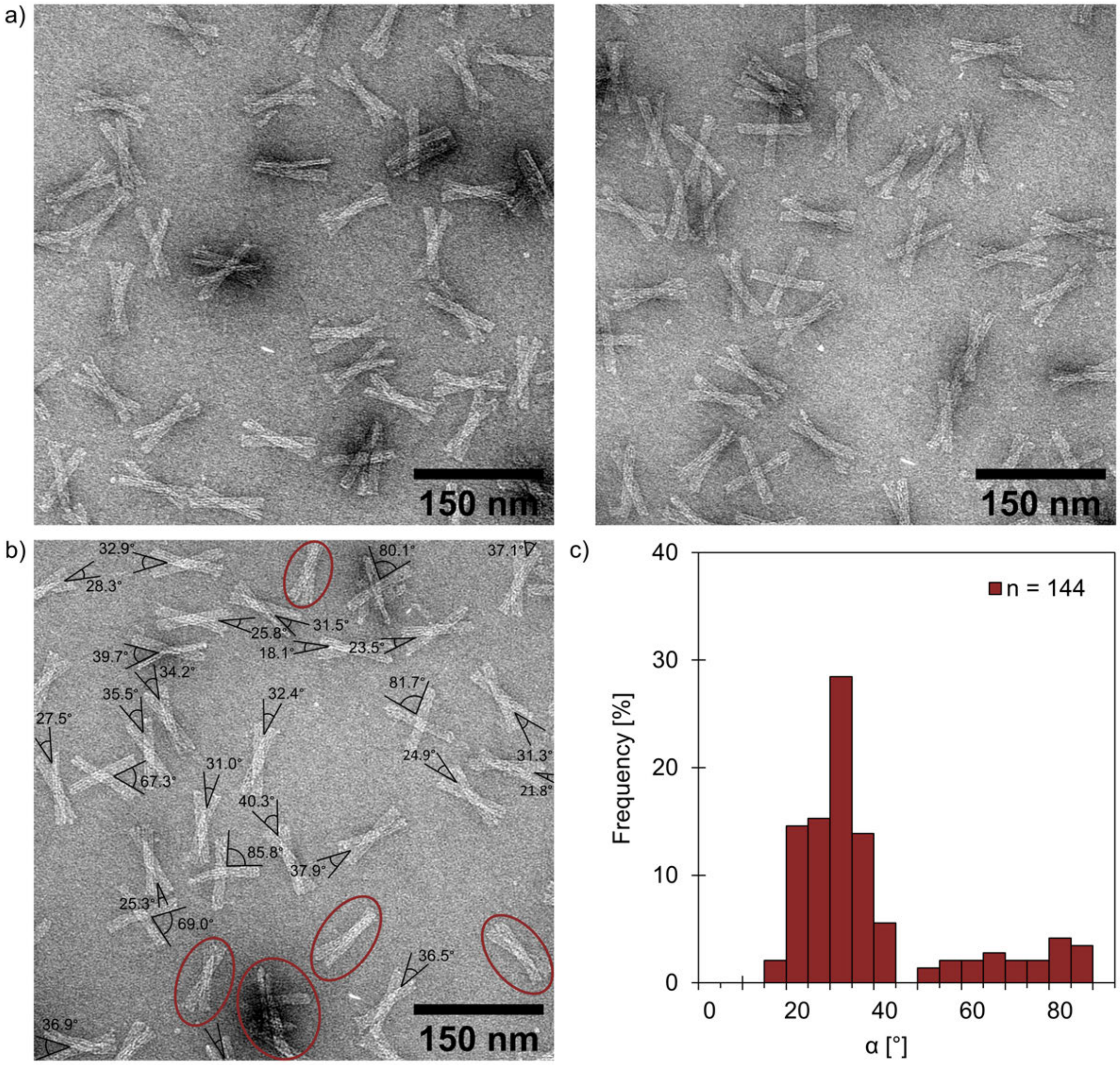


**Figure S6.** Characterization of the permanently closed DNA origami unit (poly-T passivated) at pH 6.0. (a) TEM images of the unit ($c$ = 5.0 nM) in 1× FOB (1× TAE, 20 mM $MgCl_2$, 5 mM NaCl) at pH 6.0. (b) Examples of determined angles, $\alpha$, between the two arms of the unit. The encircled structures were excluded from the analysis. (c) The distribution of the angle, $\alpha$, between the two arms of the unit. The number of individual structures analyzed for the sample is $n$ = 144. The TEM images are negatively stained with uranyl formate (2 % (w/v)).

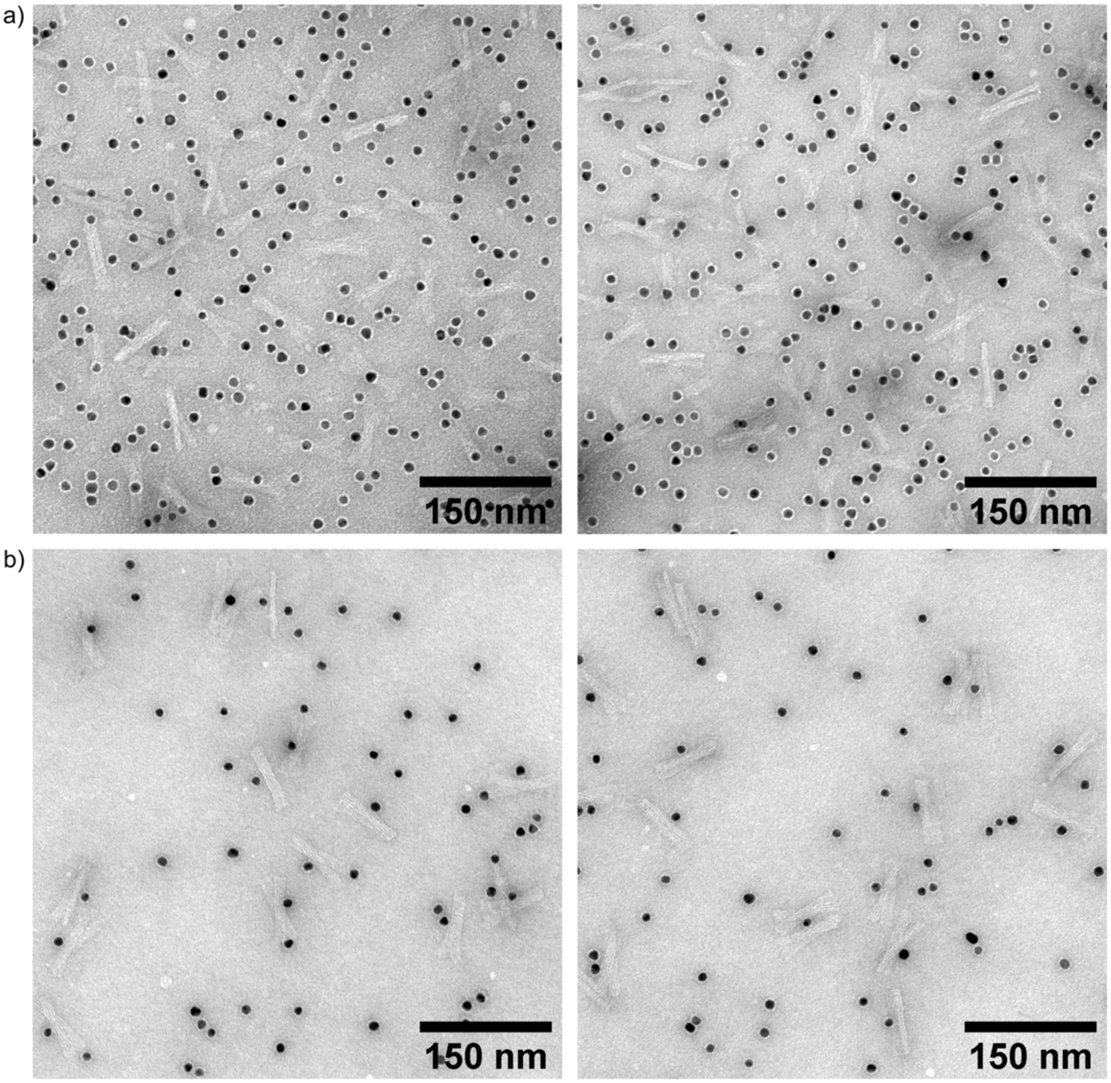


**Figure S7.** Characterization of the permanently closed DNA origami unit with scaffold loops and an anchored AuNP in the middle. TEM images of (a) unpurified units ($c$ = 7.6 nM).(b) units that have been PEG-purified twice ($c$ = 5.4 nM before the PEG purifications). The units are in 1× FOB (1× TAE, 20 mM $MgCl_2$, 5 mM NaCl) at pH 8.2 and the TEM images are negatively stained with uranyl formate (2 % (w/v)).

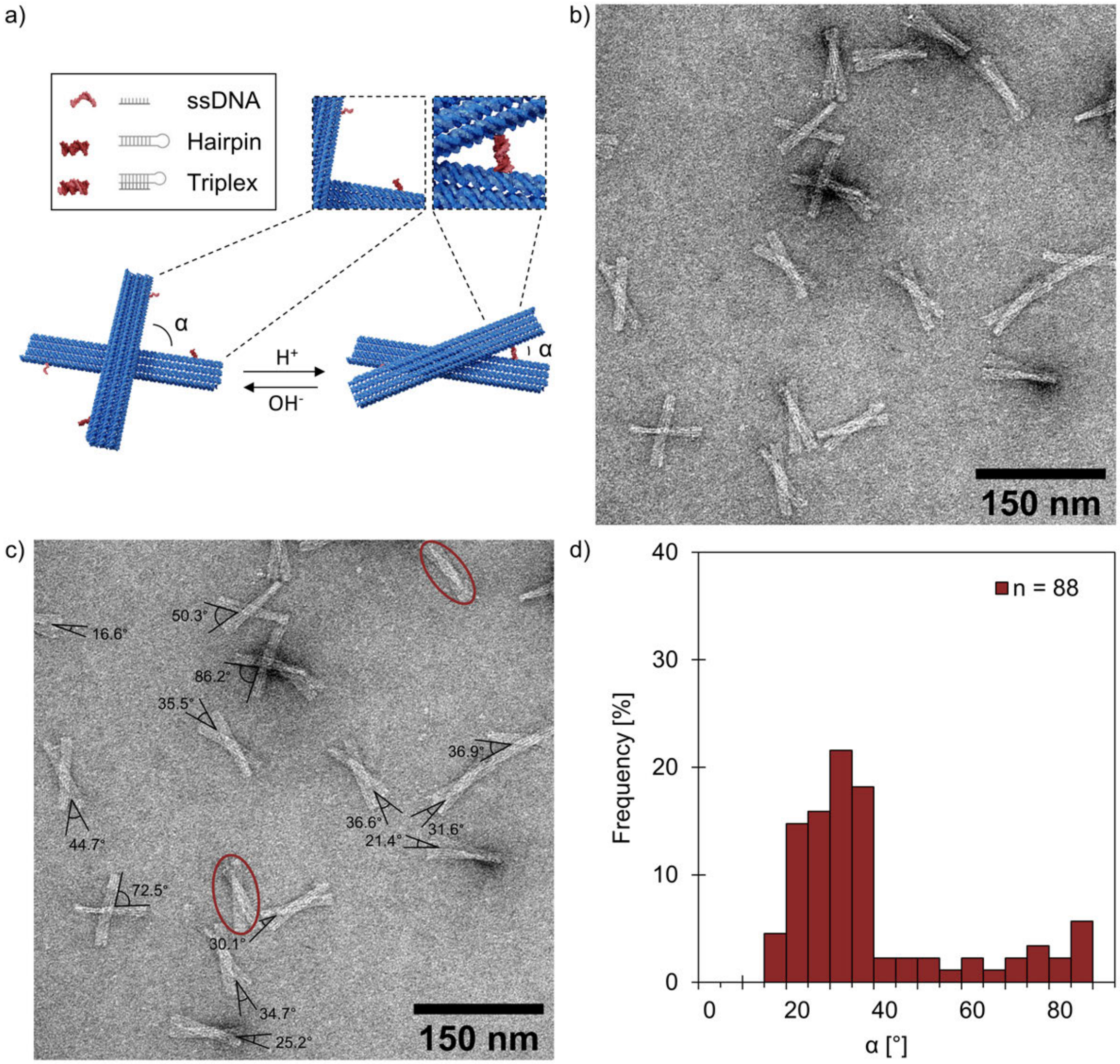


**Figure S8.** Characterization of the DNA origami unit with reverse pH latches (poly-T passivated) at pH 6.0. (a) The pH latches are at different positions in the unit with reverse pH latches and therefore the arms of the unit will be locked in a reverse direction (clockwise instead of counterclockwise). (b) TEM images of the unit ($c$ = 5.0 nM) in 1× FOB (1× TAE, 20 mM $MgCl_2$, 5 mM NaCl) at pH 6.0. (c) Examples of determined angles, $\alpha$, between the two arms of the unit. The encircled structure was excluded from the analysis. (d) The distribution of the angle, $\alpha$, between the two arms of the unit. The number of individual structures analyzed for the sample is $n$ = 88. The TEM images are negatively stained with uranyl formate (2 % (w/v)).

#### 4. pH-Responsive DNA Origami Dimers

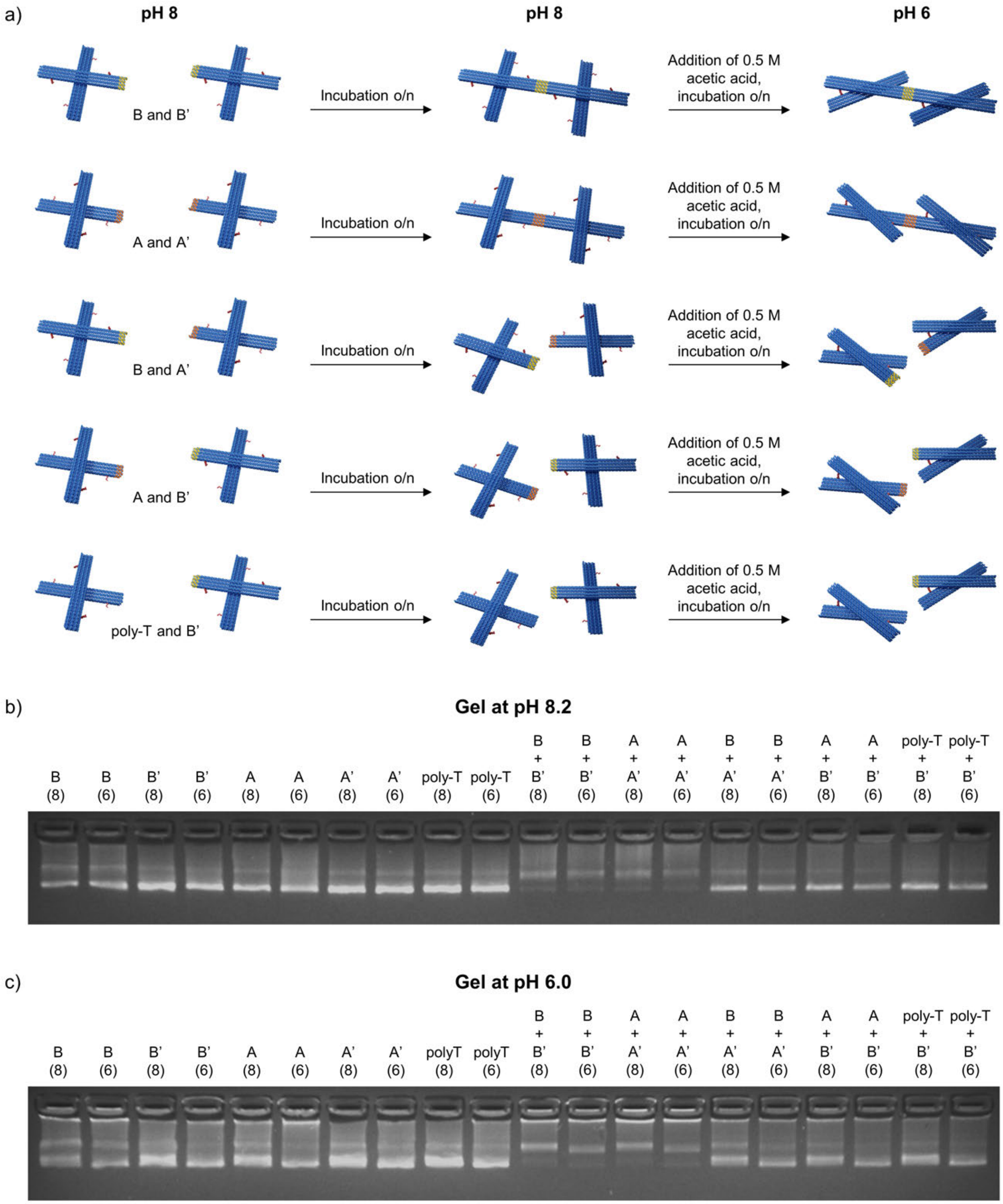


**Figure S9.** Characterization of the dimer formation at pH 8.2 by agarose gel electrophoresis. (a) Conceptual illustration of the dimerization when different DNA origami units are used. Dimers are selectively formed only for units with shape complementarity and appropriate connector oligonucleotides (mixture of B and B' units or mixture of A and A' units). The dimers formed at pH 8.2 are analyzed by agarose gel electrophoresis at (b) pH 8.2 and (c) pH 6.0 both before (samples marked '8') and after the pH has been decreased to pH 6 by 0.5 M acetic acid (samples marked '6'). The DNA origami concentrations in the gel are 15.0 nM for individual units and 5.4 nM (pH 6) or 5.9 nM (pH 8) in unit mixtures. Selected parts of the gels are also shown in the article in Figure 2b.

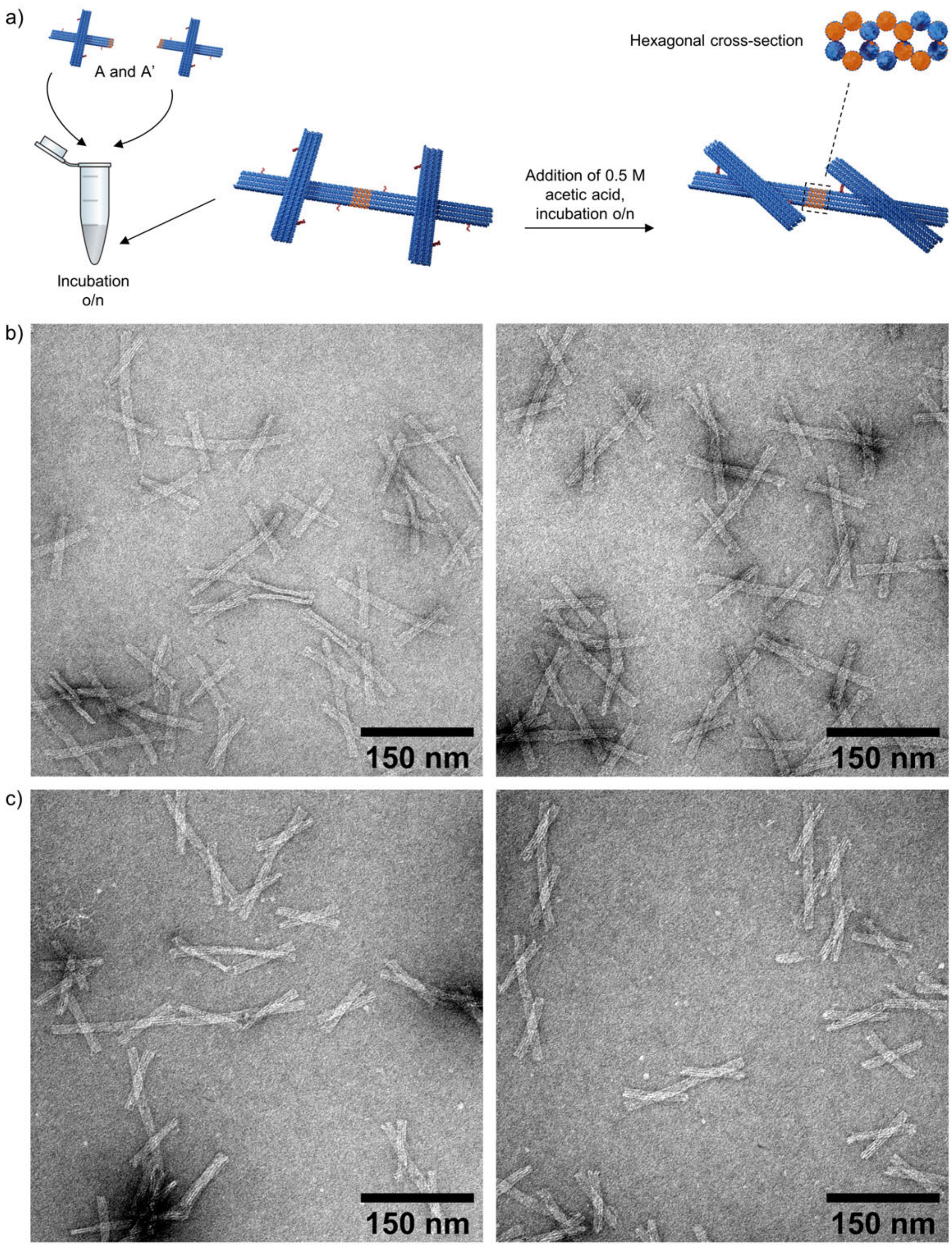


**Figure S10.** Dimer formation at pH 8.2 using A and A' DNA origami units. (a) Dimers are formed by mixing equimolar amounts of both units (A and A'). By adding acetic acid to the dimer solution, the arms could be locked into the closed configuration also after the dimerization. (b) TEM images of the dimers formed at pH 8.2 ($c_{\mathrm{dimer}}$ = 5.7 nM). (c) TEM images of the same dimer solution as in b) after the pH has been decreased to 6 ($c_{\mathrm{dimer}}$ = 5.4 nM). The TEM images are negatively stained with 2% (w/v) uranyl formate.

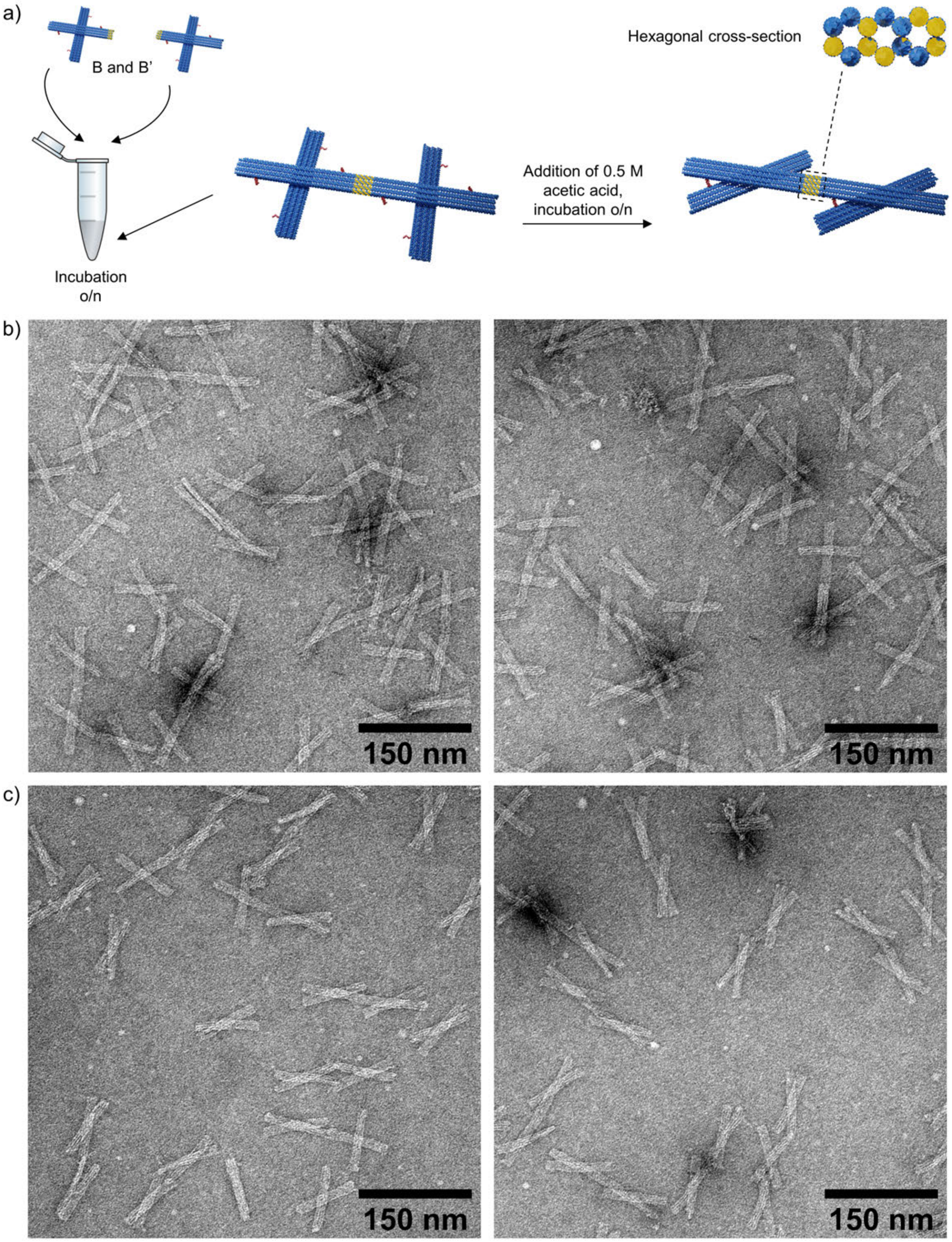


**Figure S11.** Dimer formation at pH 8.2 using B and B' DNA origami units. (a) Dimers are formed by mixing equimolar amounts of both units (B and B'). By adding acetic acid to the dimer solution, the arms could be locked into the closed configuration also after the dimerization. (b) TEM images of the dimers formed at pH 8.2 ($c_{dimer}$ = 5.7 nM). (c) TEM images of the same dimer solution as in b) after the pH has been decreased to 6 ($c_{dimer}$ = 5.4 nM). The TEM images are negatively stained with 2% (w/v) uranyl formate.

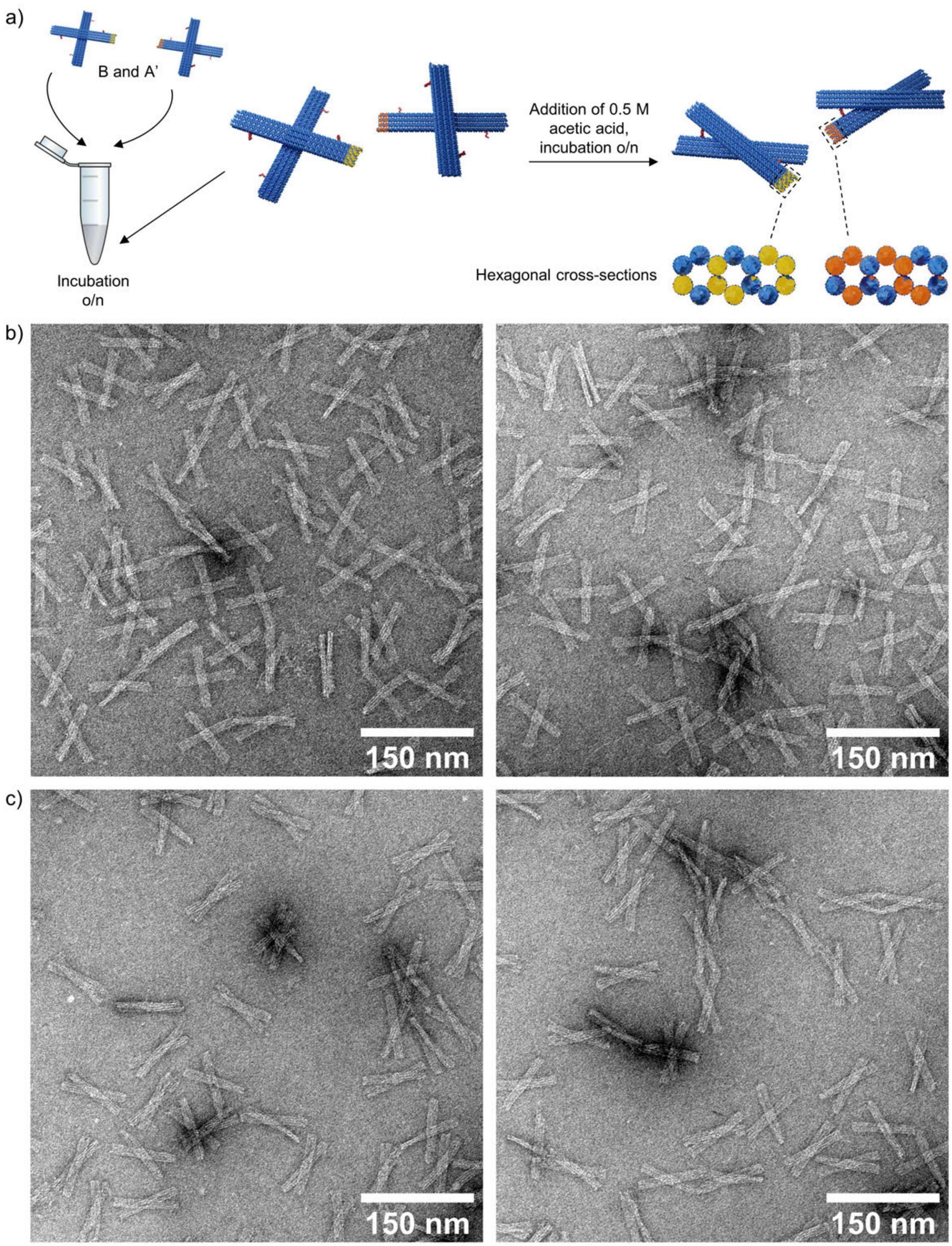


**Figure S12.** Mixture of B and A' DNA origami units at pH 8.2. (a) No dimers are formed by mixing equimolar amounts of units B and A'. By adding NaOH to the DNA origami mixture, the units will adopt the open configuration. (b) TEM images of the DNA origami mixture at pH 8.2 ($c_{\text{unit}}$ = 5.7 nM). (c) TEM images of the same DNA origami mixture as in b) after the pH has been decreased to 6 ($c_{\text{unit}}$ = 5.4 nM). The TEM images are negatively stained with 2% (w/v) uranyl formate.

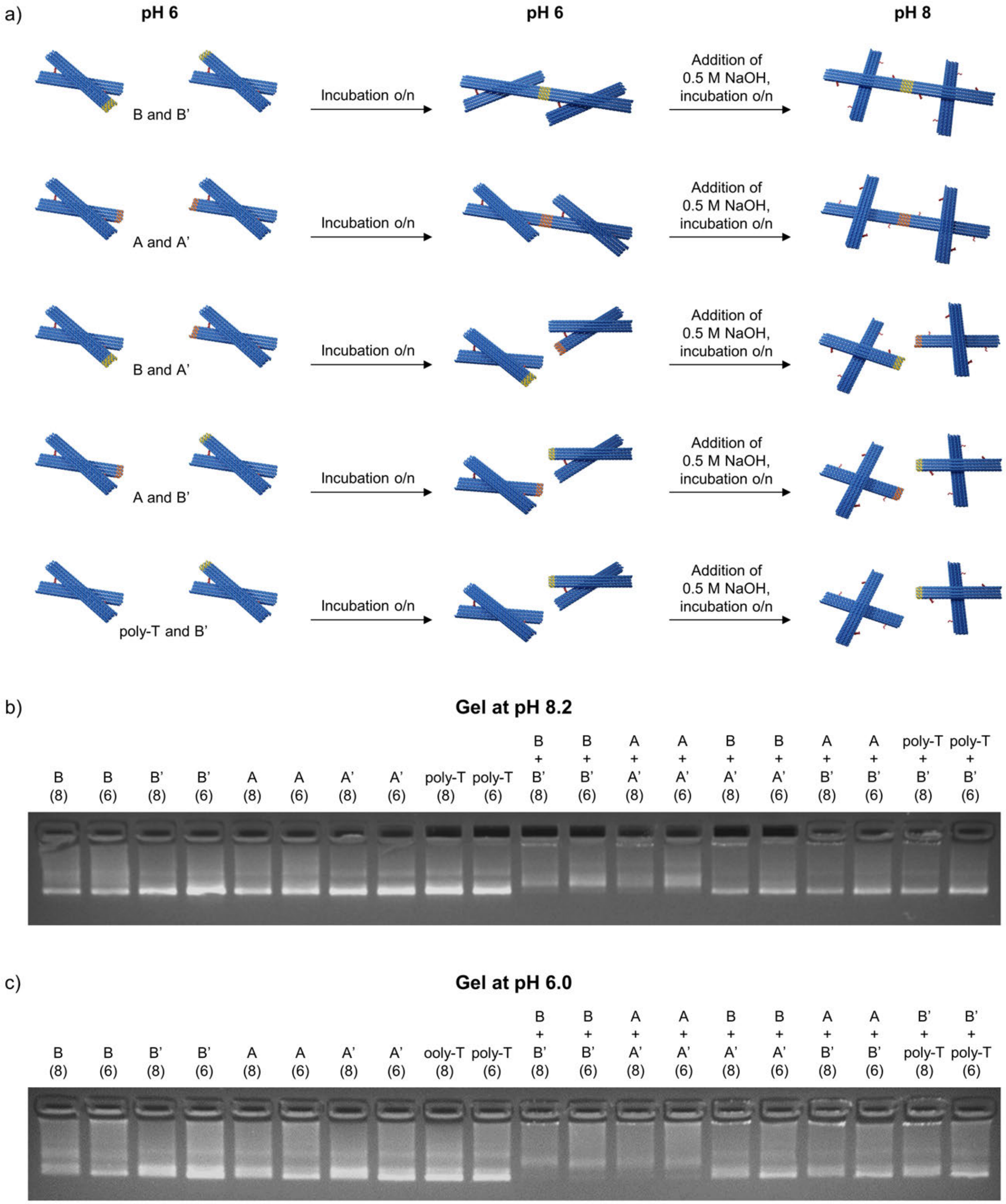


**Figure S13.** Characterization of the dimer formation at pH 6.0 by agarose gel electrophoresis. (a) Conceptual illustration of the dimerization when different DNA origami units are used. Dimers are selectively formed only for units with shape complementarity and appropriate connector oligonucleotides (mixture of B and B' units or mixture of A and A' units). The dimers formed at pH 6.0 are analyzed by agarose gel electrophoresis at (b) pH 8.2 and (c) pH 6.0 both before (samples marked '6') and after the pH has been increased to pH 8 by 0.5 M NaOH (samples marked '8'). The DNA origami concentrations in the gel are 15.0 nM for individual units and 5.4 nM (pH 8) or 5.9 nM (pH 6) in unit mixtures.

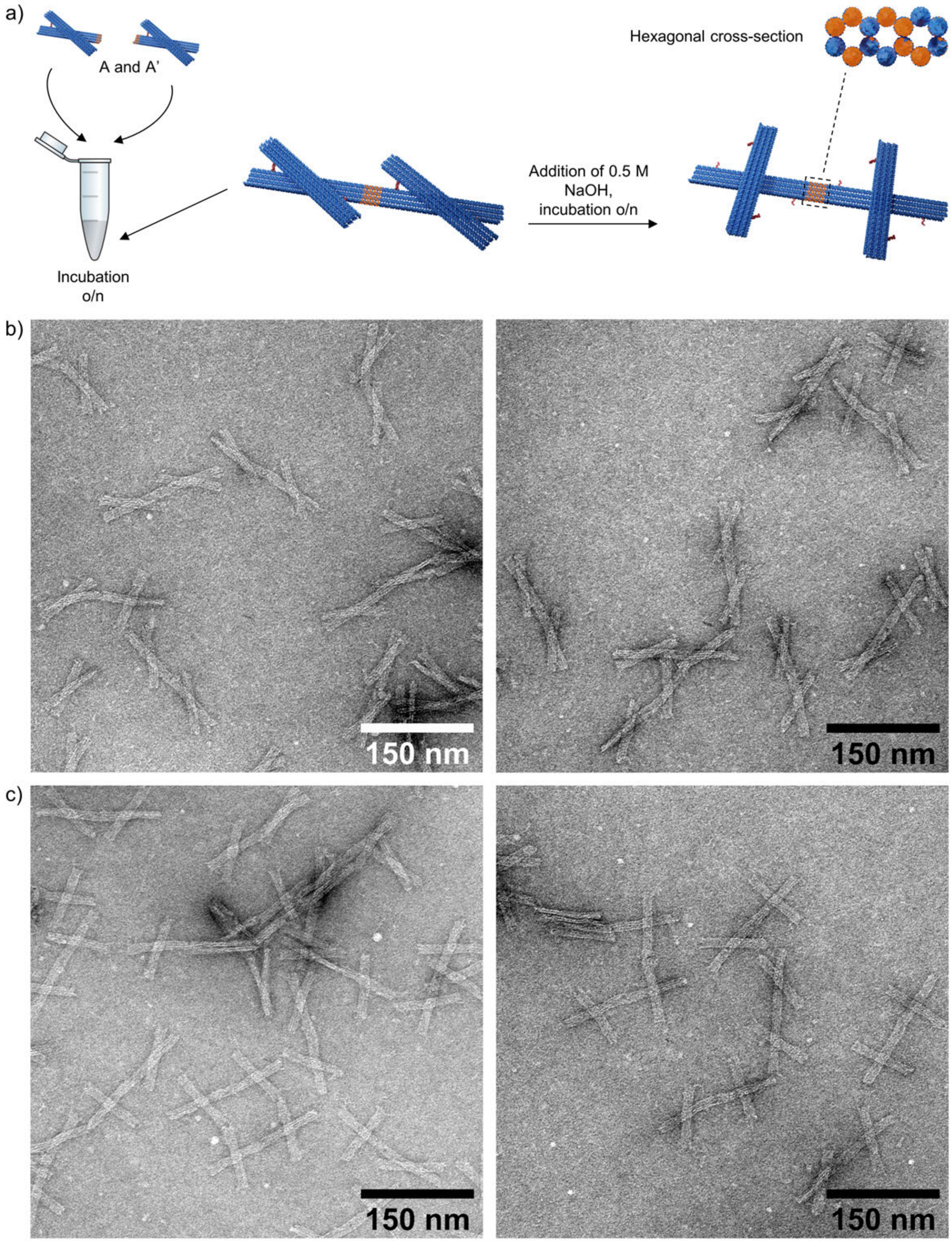


**Figure S14.** Dimer formation at pH 6.0 using A and A' DNA origami units. (a) Dimers are formed by mixing equimolar amounts of both units (A and A'). By adding NaOH to the dimer solution, the arms could be released again also after the dimerization. (b) TEM images of the dimers formed at pH 6.0 ($c_{\mathrm{dimer}}$ = 5.7 nM). (c) TEM images of the same dimer solution as in b) after the pH has been increased to 8 ($c_{\mathrm{dimer}}$ = 5.4 nM). The TEM images are negatively stained with 2% (w/v) uranyl formate.

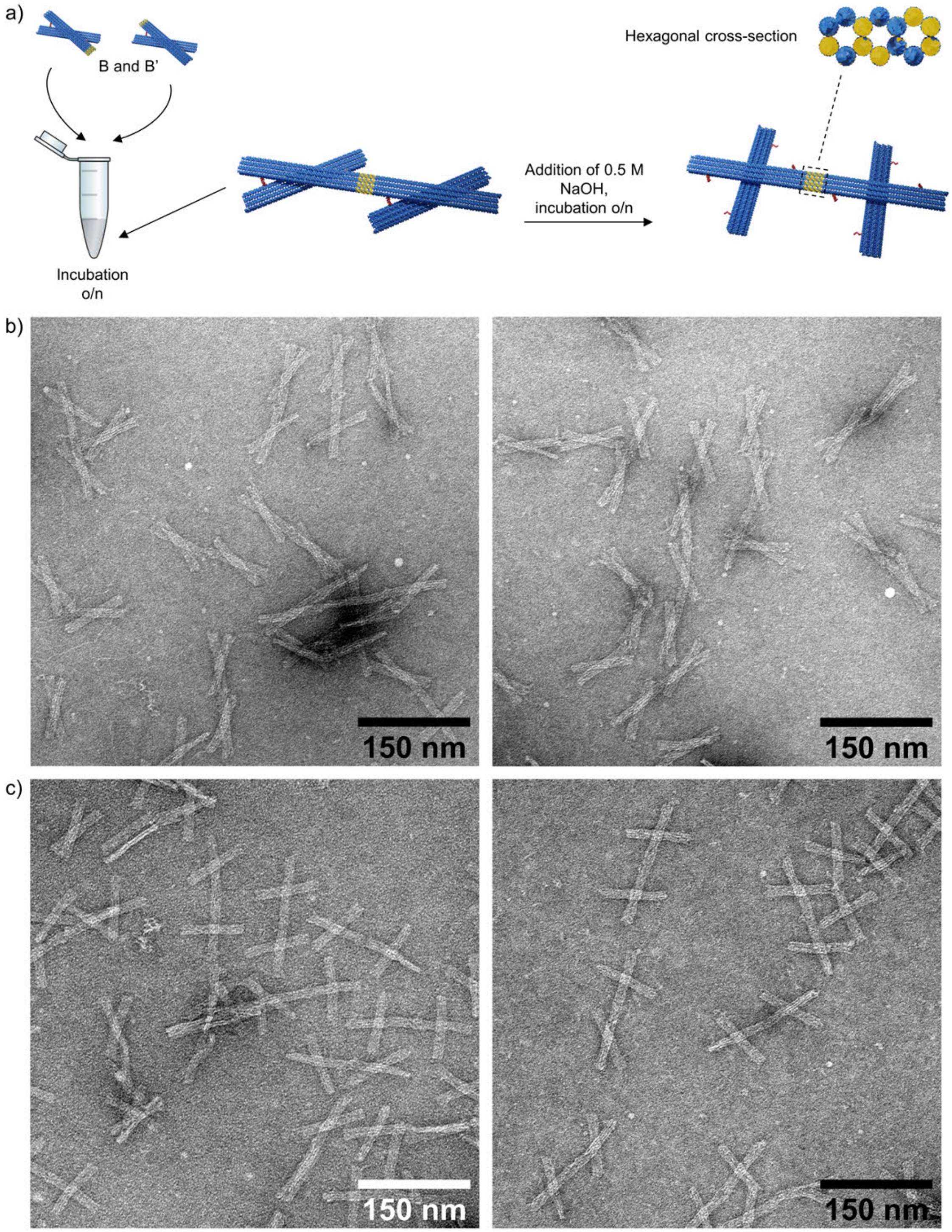


**Figure S15.** Dimer formation at pH 6.0 using B and B' DNA origami units. (a) Dimers are formed by mixing equimolar amounts of both units (B and B'). By adding NaOH to the dimer solution, the arms could be released again also after the dimerization. (b) TEM images of dimers formed at pH 6.0 ($c_{dimer}$ = 5.7 nM). (c) TEM images of the same dimer solution as in b) after the pH has been increased to 8 ($c_{dimer}$ = 5.4 nM). The TEM images are negatively stained with 2% (w/v) uranyl formate.

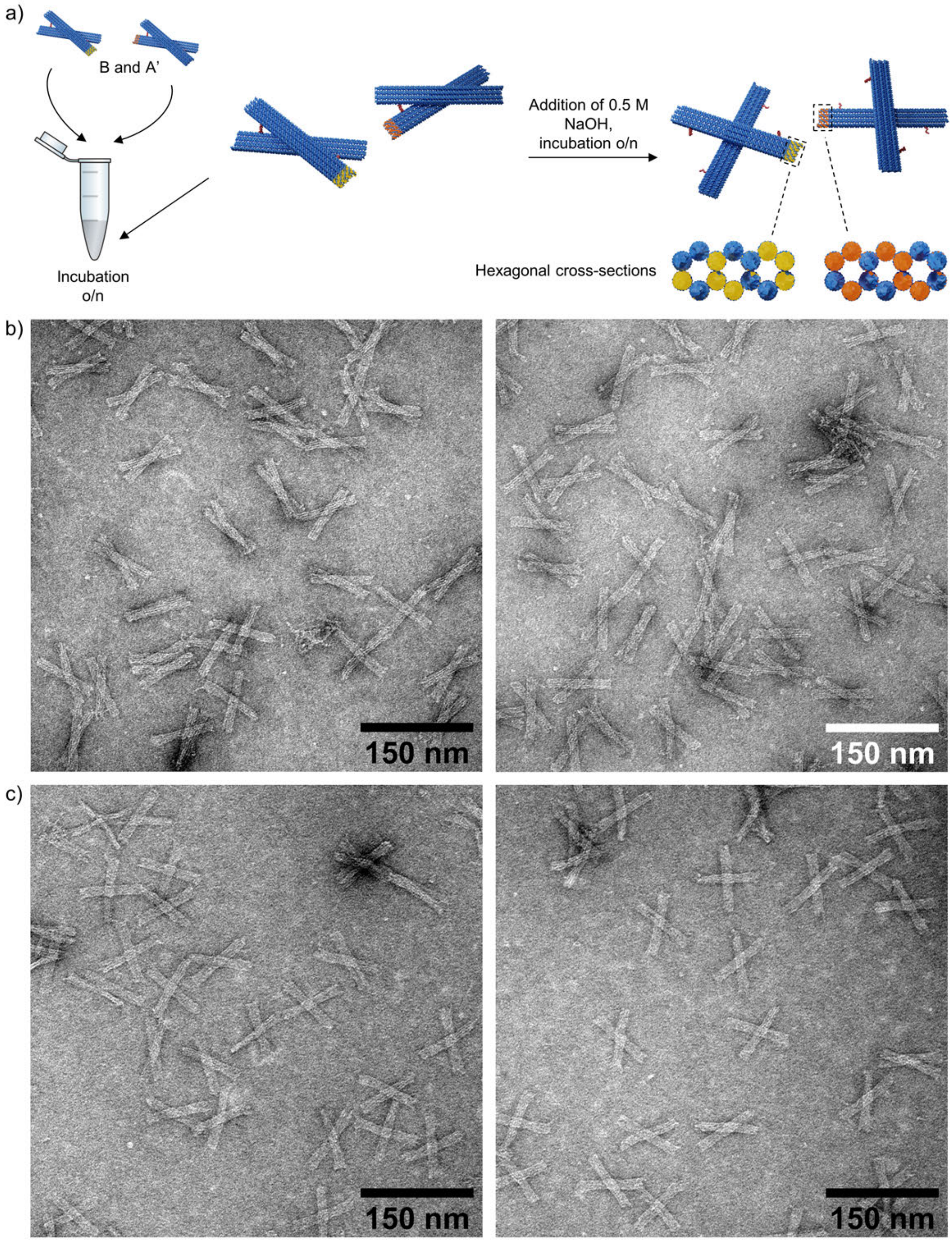


**Figure S16.** Mixture of B and A' DNA origami units at pH 6.0 (a) No dimers are formed by mixing equimolar amounts of units B and A'. By adding NaOH to the DNA origami mixture, the units will adopt the open configuration. (b) TEM images of the DNA origami mixture at pH 6.0 ($c_{\text{unit}}$ = 5.7 nM). (c) TEM images of the same DNA origami mixture as in b) after the pH has been increased to 6 ($c_{\text{unit}}$ = 5.4 nM). The TEM images are negatively stained with 2% (w/v) uranyl formate.

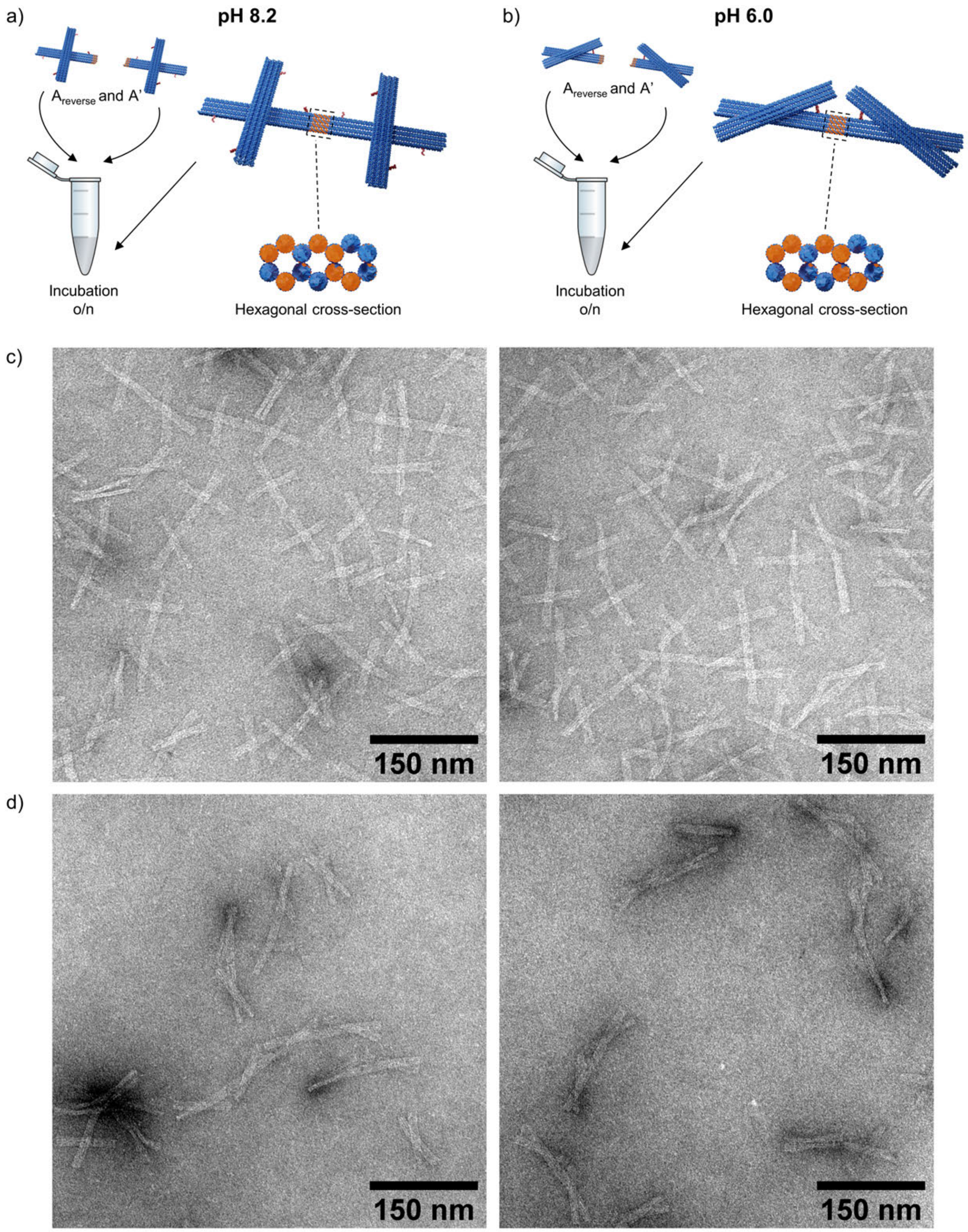


**Figure S17.** Dimer formation using units with different types of pH locks. Dimers are formed using equimolar amounts of units $A_{reverse}$ and A'. The $A_{reverse}$ unit has the reversed pH latches, whereas the A' unit have the regular pH latches. The dimers assembled at (a) pH 8.2 have their arms in an open configuration, whereas the dimers assembled at (b) pH 6.0 have arms that point towards each other. TEM images of the dimers formed at (c) pH 8.2 ($c_{dimer}$ = 5.7 nM) and (d) pH 6.0 ($c_{dimer}$ = 5.7 nM. The TEM images are negatively stained with 2% (w/v) uranyl formate.

#### 5. pH-Responsive 1D DNA Origami Arrays

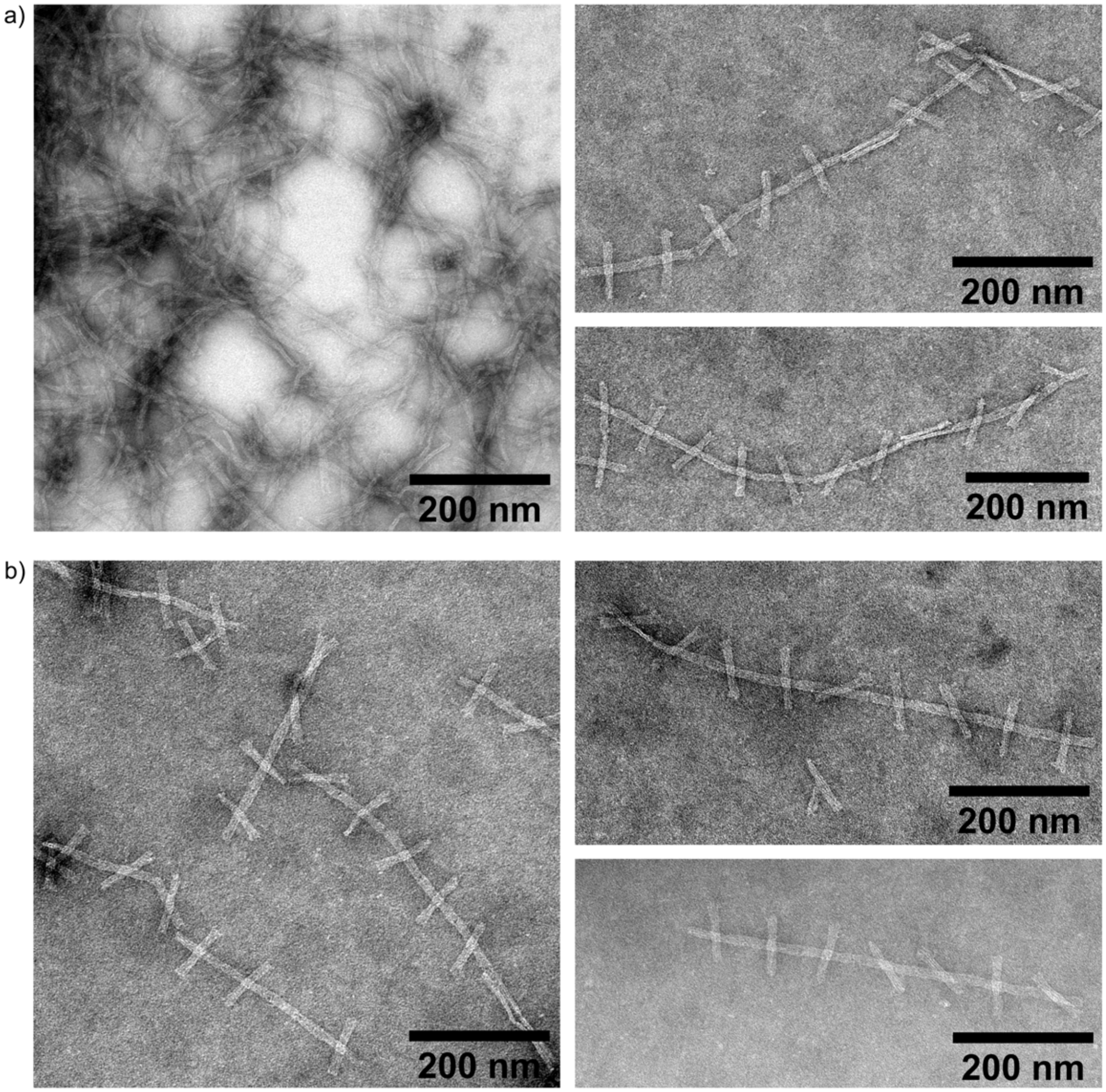


**Figure S18.** TEM images of linear 1D arrays formed in solution at pH 8.2 (25 h incubation at room temperature). The 1D chains were assembled using DNA origami unit concentrations of (a) 10.0 nM (diluted 1:2 in 1× FOB (1× TAE, 20 mM $MgCl_2$, 5 mM NaCl) before deposited onto the TEM grid) (b) 5.0 nM. The TEM images are negatively stained with 2% (w/v) uranyl formate.

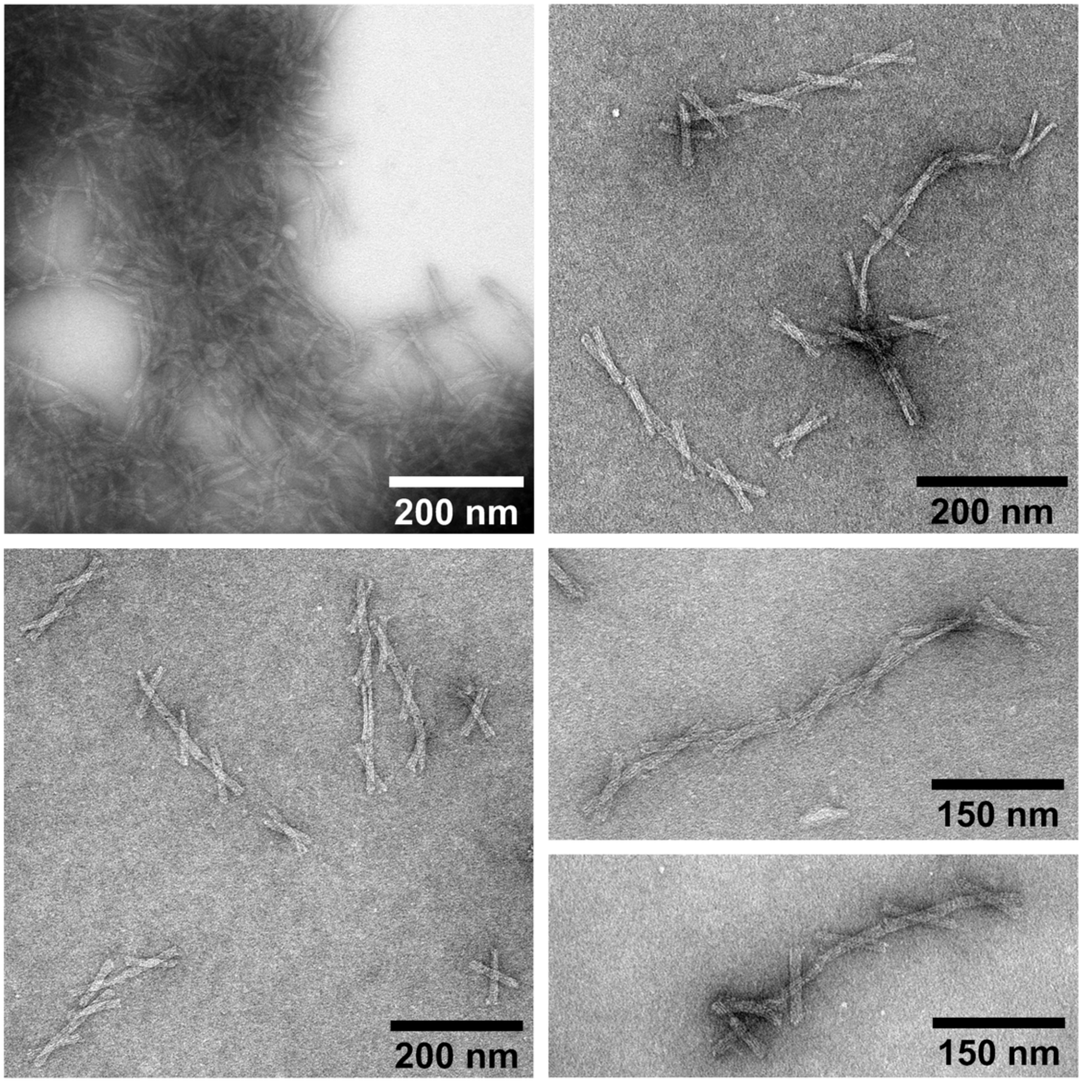


**Figure S19.** TEM images of linear 1D arrays formed in solution at pH 6.0 (24 h incubation at room temperature). The 1D chains were assembled using a DNA origami unit concentration of 2.0 nM. The TEM images are negatively stained with 2% (w/v) uranyl formate.

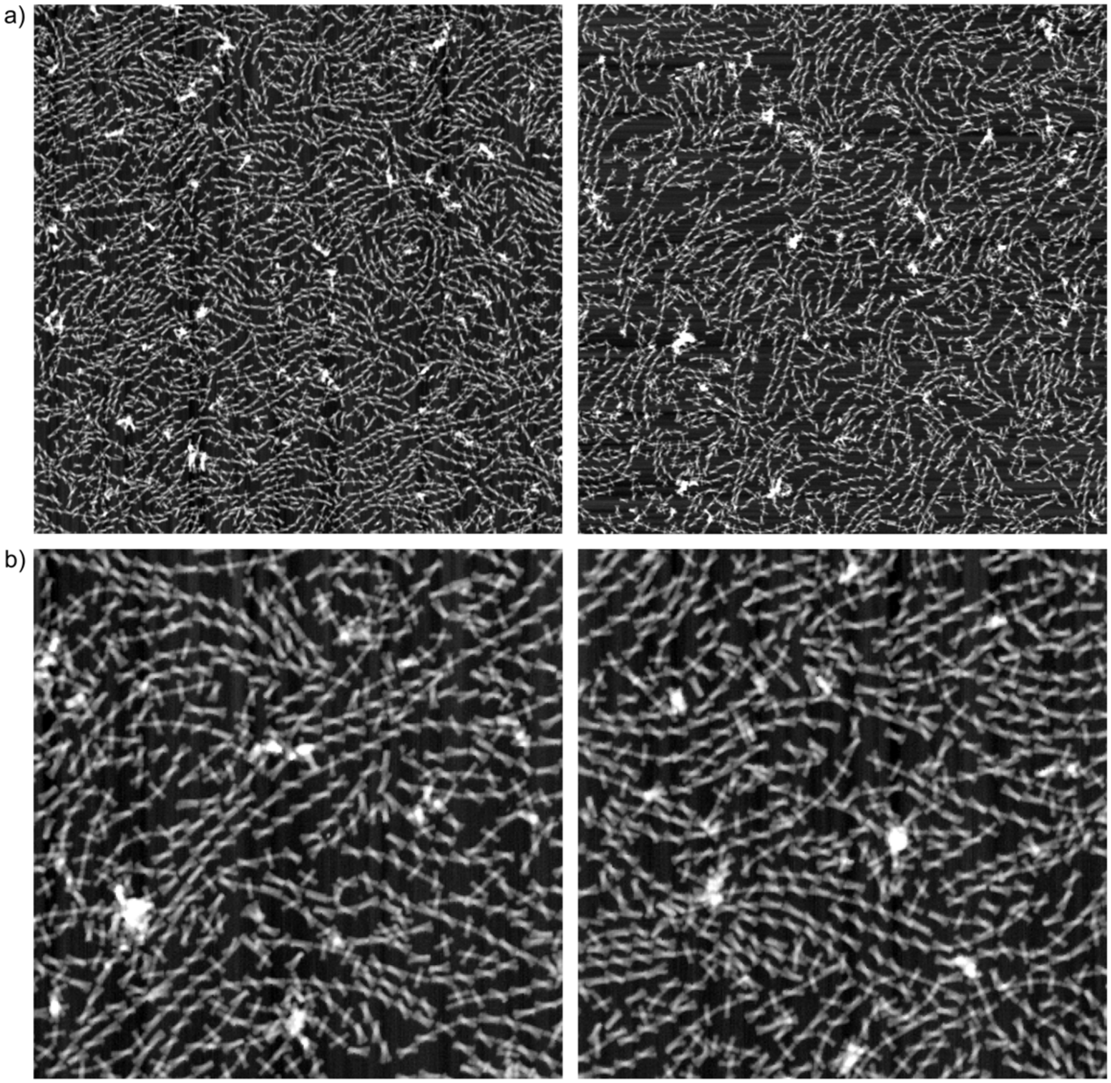


**Figure S20.** AFM images demonstrating the 1D array formation on the mica surface at pH 6.0. The DNA origami concentration in the sample is 2.0 nM. The DNA origami concentration in the sample is 2.0 nM and the assembly is done in 1× TAE containing 10 mM $MgCl_2$ and 75 mM NaCl. The lattice was formed on the mica surface during 3 h. (a) Large-scale AFM image (5 μm × 5 μm) of the formed linear chains. (b) Magnified AFM images (2 μm × 2 μm) of the assembled 1D arrays.

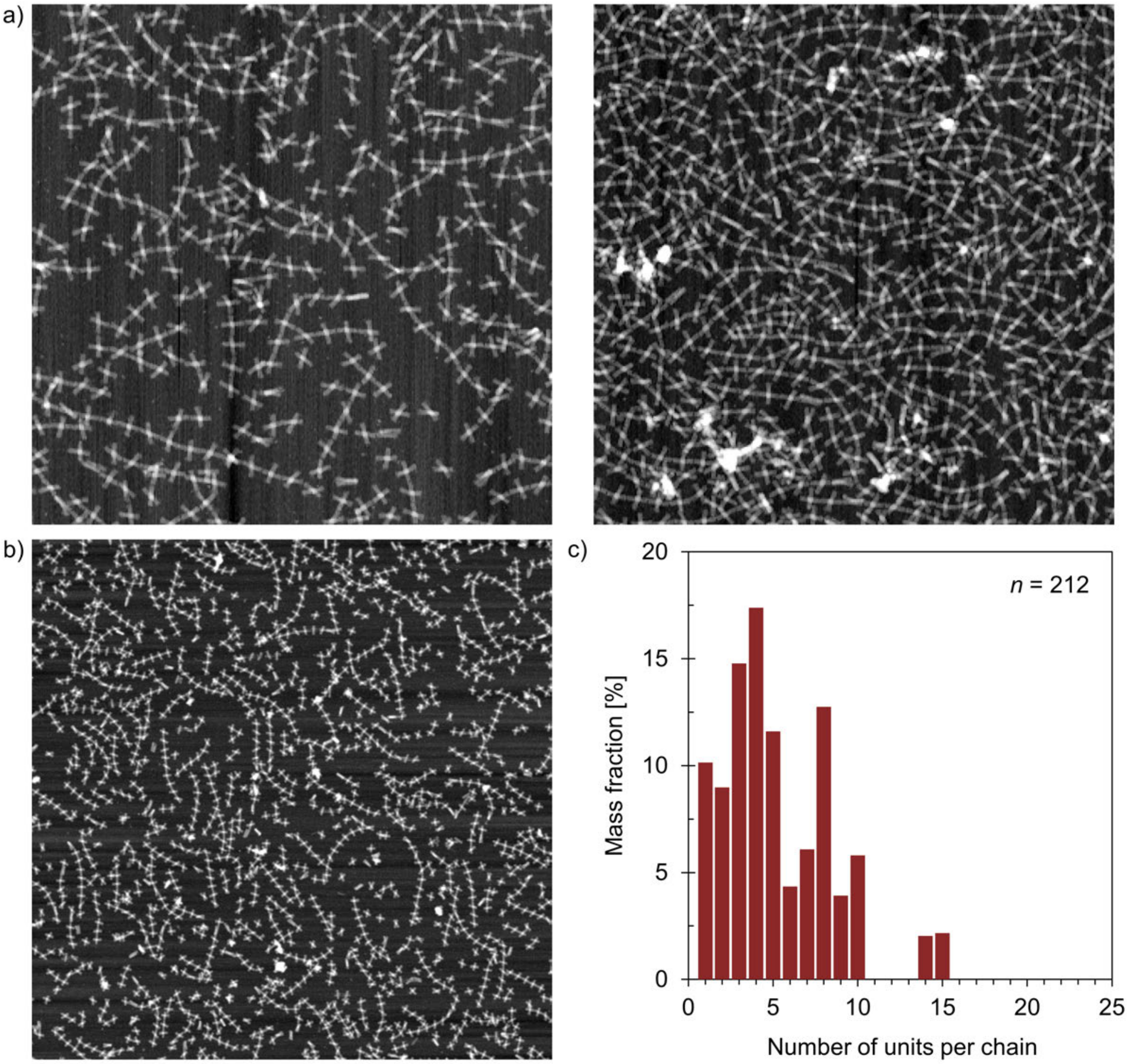


**Figure S21.** AFM images demonstrating the 1D array formation on the mica surface at pH 8.2 using the DNA origami unit with pH latches. The DNA origami concentration in the sample is 2.0 nM and the assembly is done in 1× TAE containing 10 mM $MgCl_2$ and 75 mM NaCl. The lattice was formed on the mica surface during 3 h. (a) AFM images (2 μm × 2 μm) of the assembled 1D arrays. (b) Large-scale AFM image (5 μm × 5 μm) of the formed linear chains. (c) Observed chain length distribution for the 1D arrays assembled at pH 8.2 (determined from AFM images).

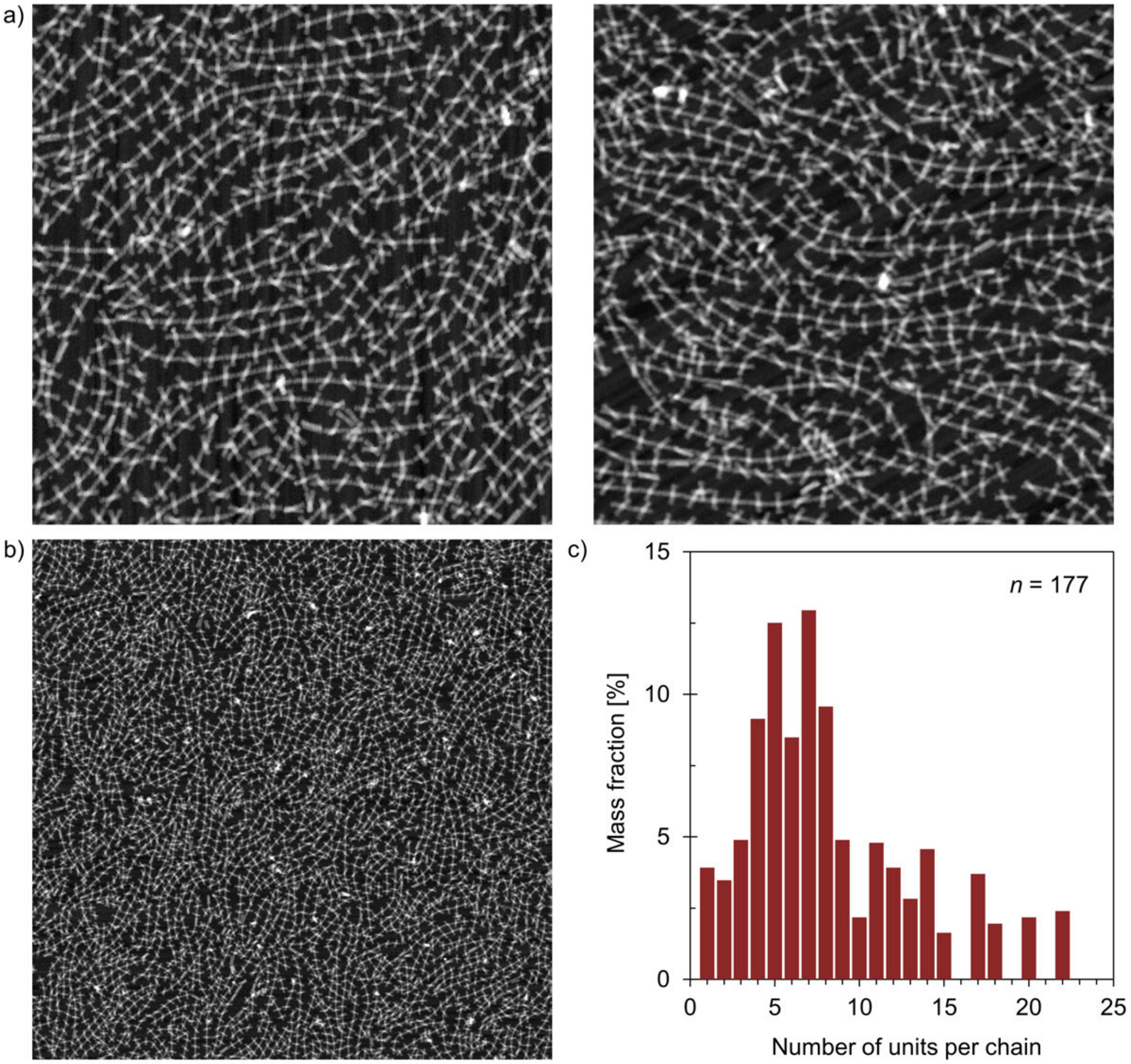


**Figure S22.** AFM images demonstrating the 1D array formation on the mica surface at pH 8.2 using the DNA origami unit with reverse pH latches. The DNA origami concentration in the sample is 2.0 nM and the assembly is done in 1× TAE containing 10 mM $MgCl_2$ and 75 mM NaCl. The lattice was formed on the mica surface during 5 h. (a) AFM images (2 μm × 2 μm) of the assembled 1D arrays. (b) Large-scale AFM image (5 μm × 5 μm) of the formed linear chains. (c) Observed chain length distribution for the 1D arrays assembled at pH 8.2 (determined from AFM images).

#### 6. pH-Responsive and Reconfigurable 2D DNA Origami Lattices

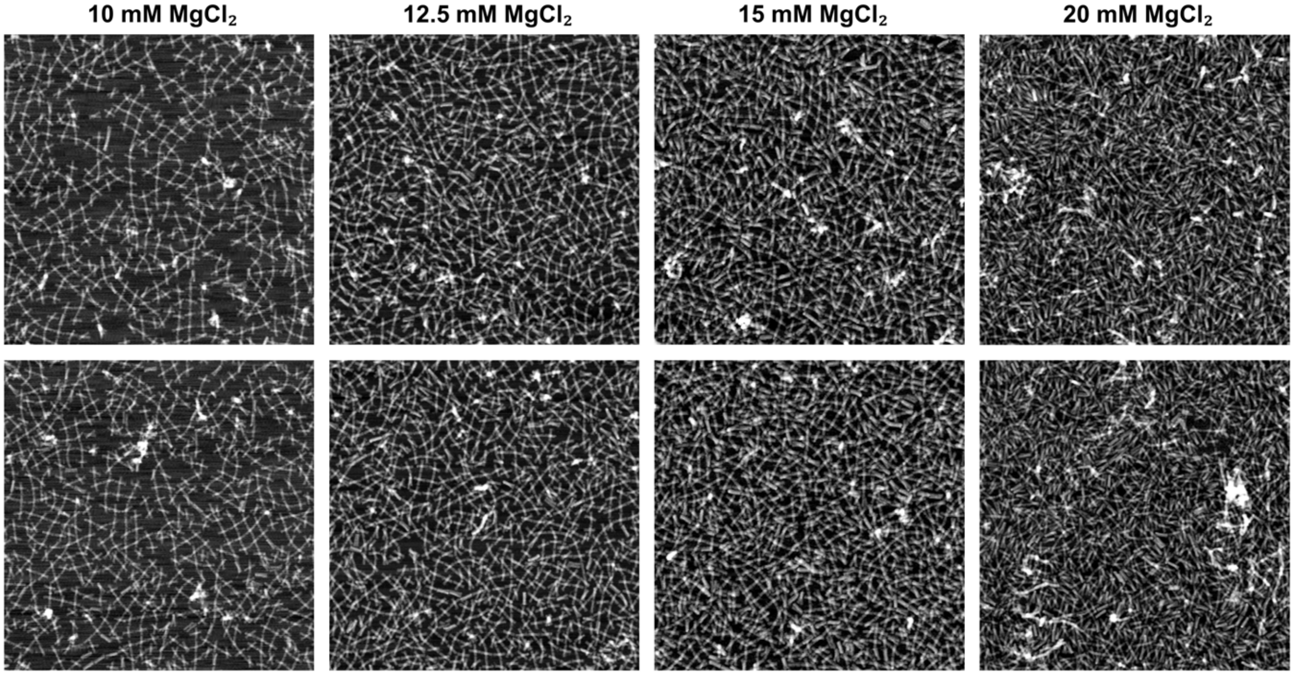


**Figure S23.** AFM images demonstrating the 2D lattice formation on the mica surface at pH 8.2 using the DNA origami unit with pH latches and different $MgCl_2$ concentrations. The DNA origami concentration in the sample is 2.0 nM and the assembly is done in 1× TAE containing $MgCl_2$ and 75 mM NaCl. The lattice was formed on the mica surface during 3 h. The size of the images is 2 μm × 2 μm.

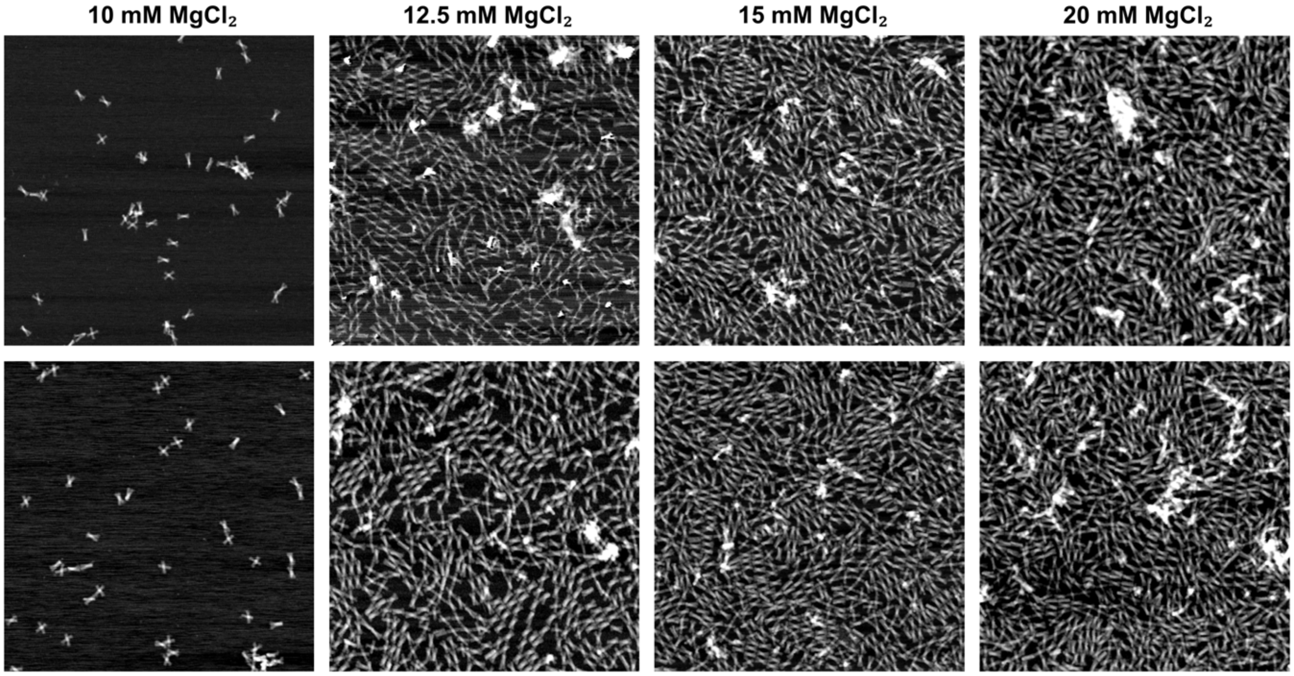


**Figure S24.** AFM images demonstrating the 2D lattice formation on the mica surface at pH 6.0 using the DNA origami unit with pH latches and different $MgCl_2$ concentrations. The DNA origami concentration in the sample is 2.0 nM and the assembly is done in 1× TAE containing $MgCl_2$ and 75 mM NaCl. The lattice was formed on the mica surface during 3 h. The size of the images is 2 $\mu$m × 2 $\mu$m.

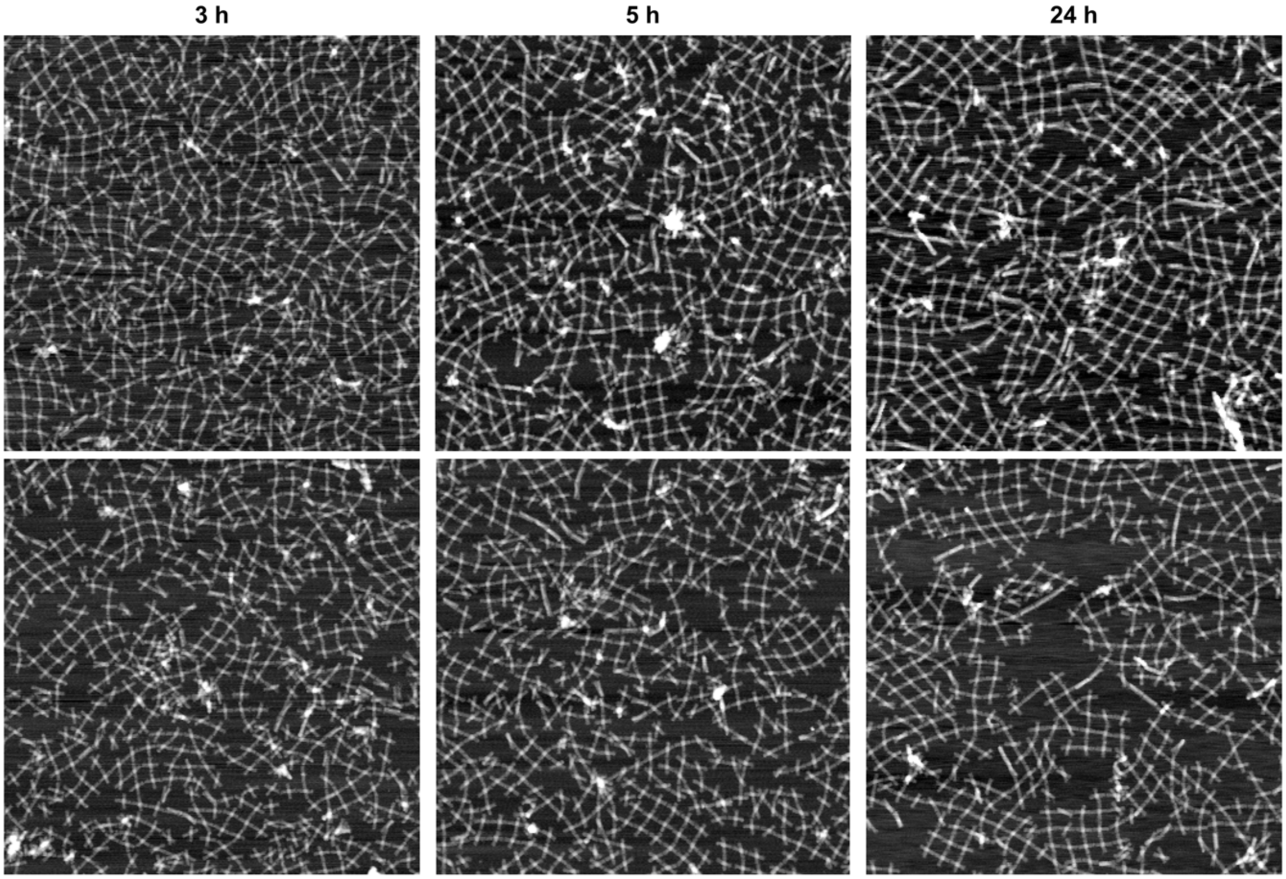


**Figure S25.** AFM images demonstrating the 2D lattice formation on mica at pH 8.2 using the DNA origami unit with pH latches and different incubation times. The DNA origami concentration in the sample is 2.0 nM and the assembly is done in 1× TAE containing 10 mM $MgCl_2$ and 75 mM NaCl. The size of the images is 2 μm × 2 μm.

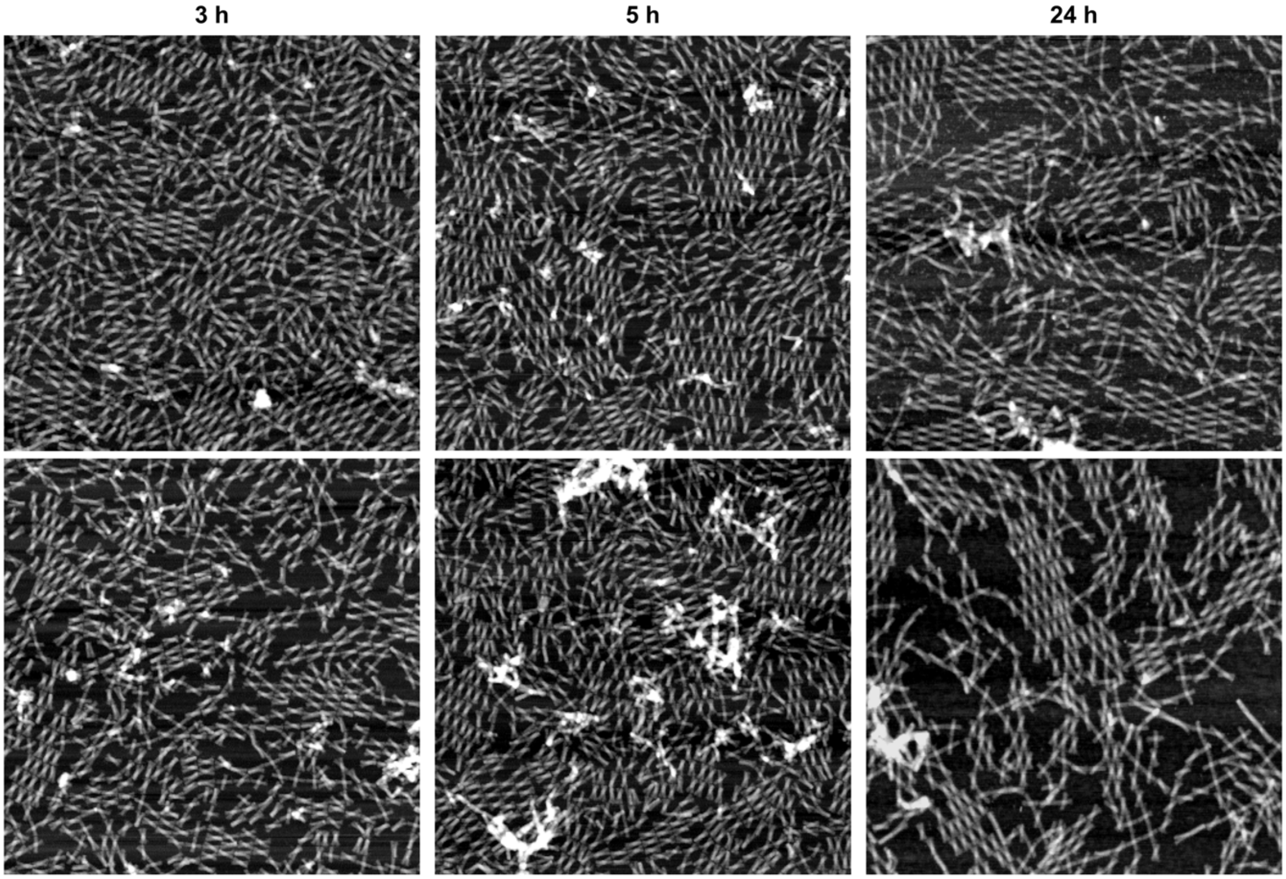


**Figure S26.** AFM images demonstrating the 2D lattice formation on mica at pH 6.0 using the DNA origami unit with pH latches and different incubation times. The DNA origami concentration in the sample is 2.0 nM and the assembly is done in 1× TAE containing 12.5 mM $MgCl_2$ and 75 mM NaCl. The size of the images is 2 μm × 2 μm.

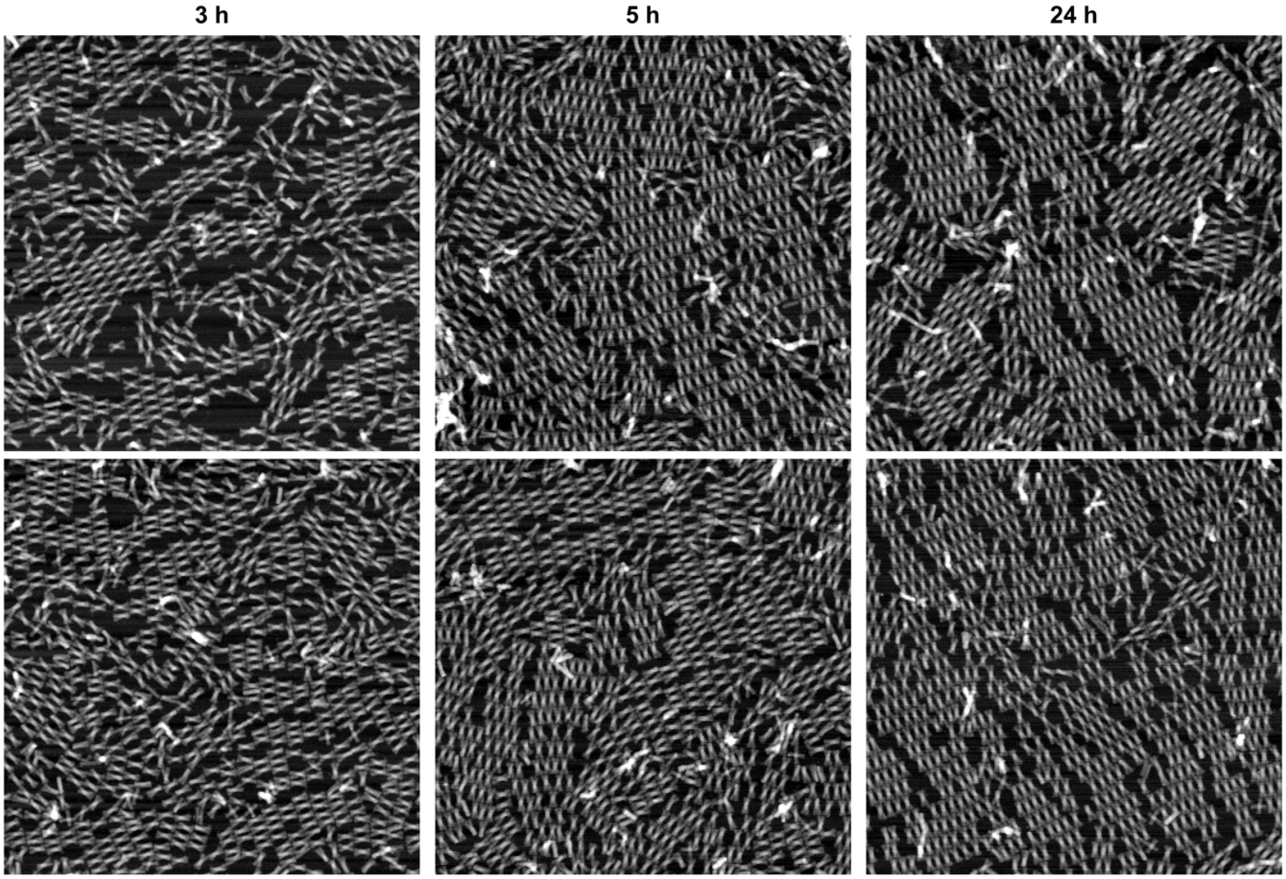


**Figure S27.** AFM images demonstrating the 2D lattice formation on the mica surface at pH 8.2 using the permanently closed DNA origami unit and different incubation times. The DNA origami concentration in the sample is 2.0 nM and the assembly is done in 1× TAE containing 10 mM $MgCl_2$ and 75 mM NaCl. The size of the images is 2 μm × 2 μm.

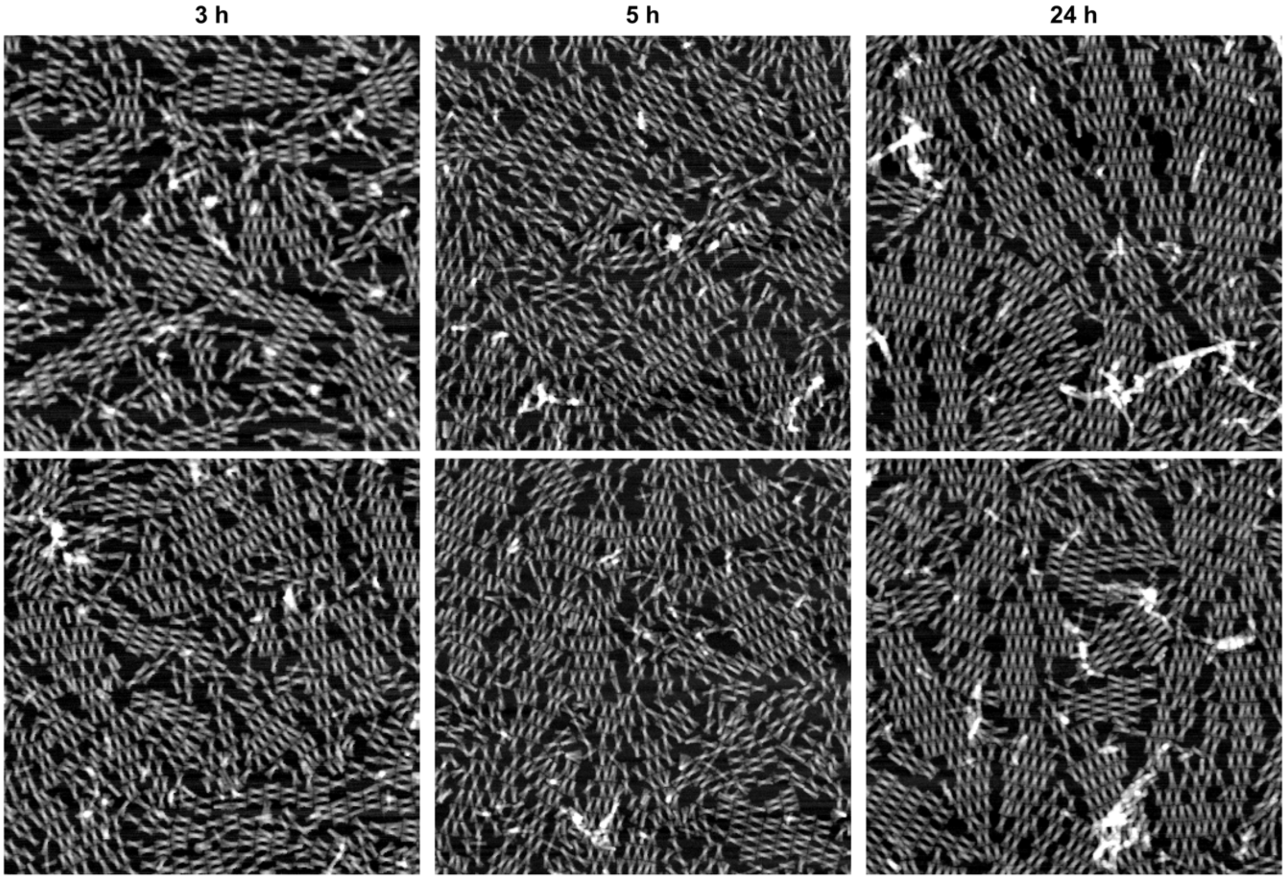


**Figure S28.** AFM images demonstrating the 2D lattice formation on the mica surface at pH 6.0 using the permanently closed DNA origami unit and different incubation times. The DNA origami concentration in the sample is 2.0 nM and the assembly is done in 1$\times$ TAE containing 12.5 mM $MgCl_2$ and 75 mM NaCl. The size of the images is 2 $\mu$m $\times$ 2 $\mu$m.

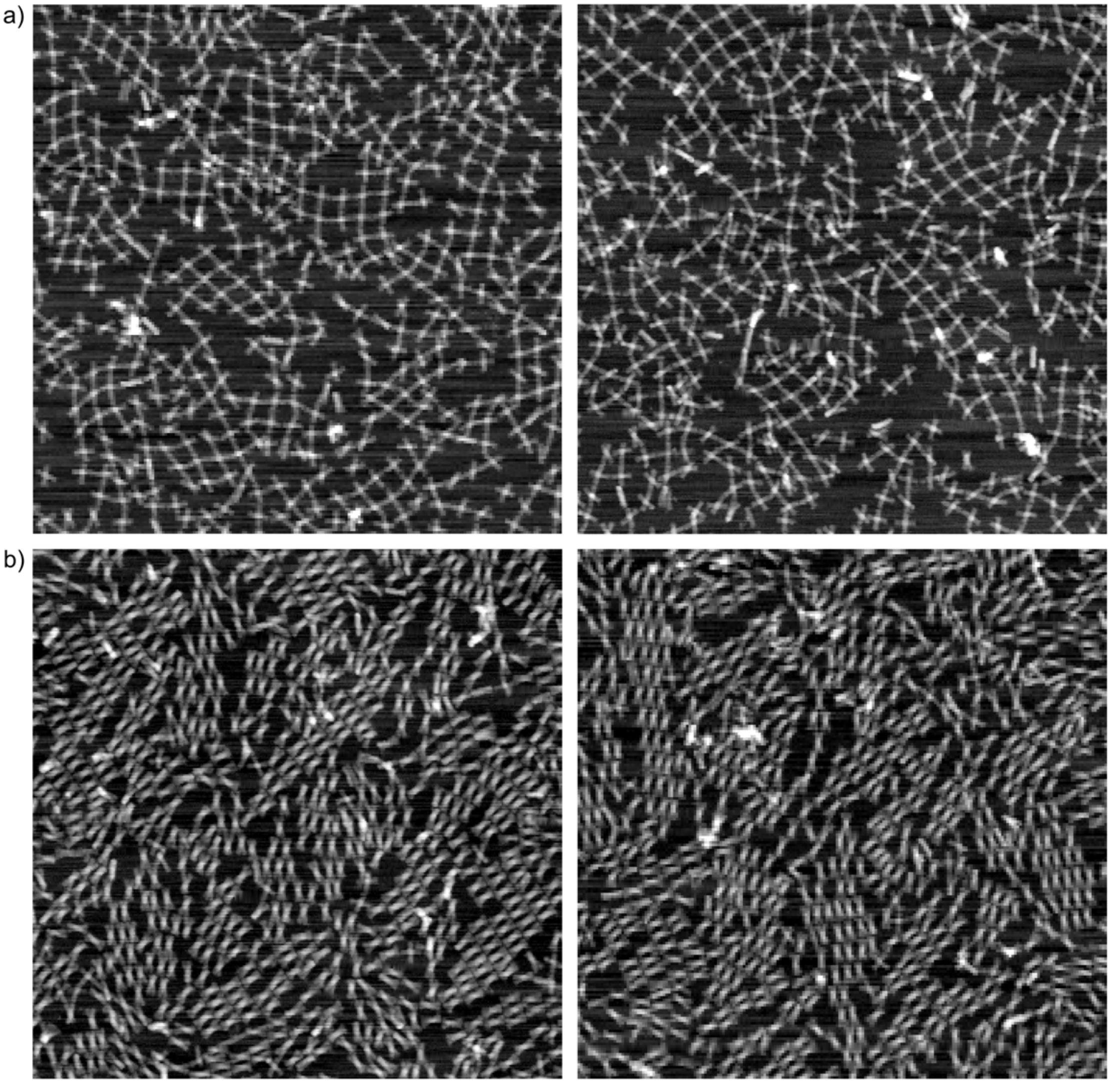


**Figure S29.** AFM images demonstrating the 2D lattice formation on mica using $CaCl_2$ instead of $MgCl_2$. (a) 2D lattice formation on mica at pH 8.2 using the DNA origami unit with pH latches. (b) 2D lattice formation on mica at pH 8.2 using the permanently closed DNA origami unit. The DNA origami concentration in the samples is 2.0 nM and the assembly is done in 1× TAE containing 10 mM $CaCl_2$ and 75 mM NaCl. The lattices were formed on the mica surface during 3 h. The size of the images is 2 μm × 2 μm.

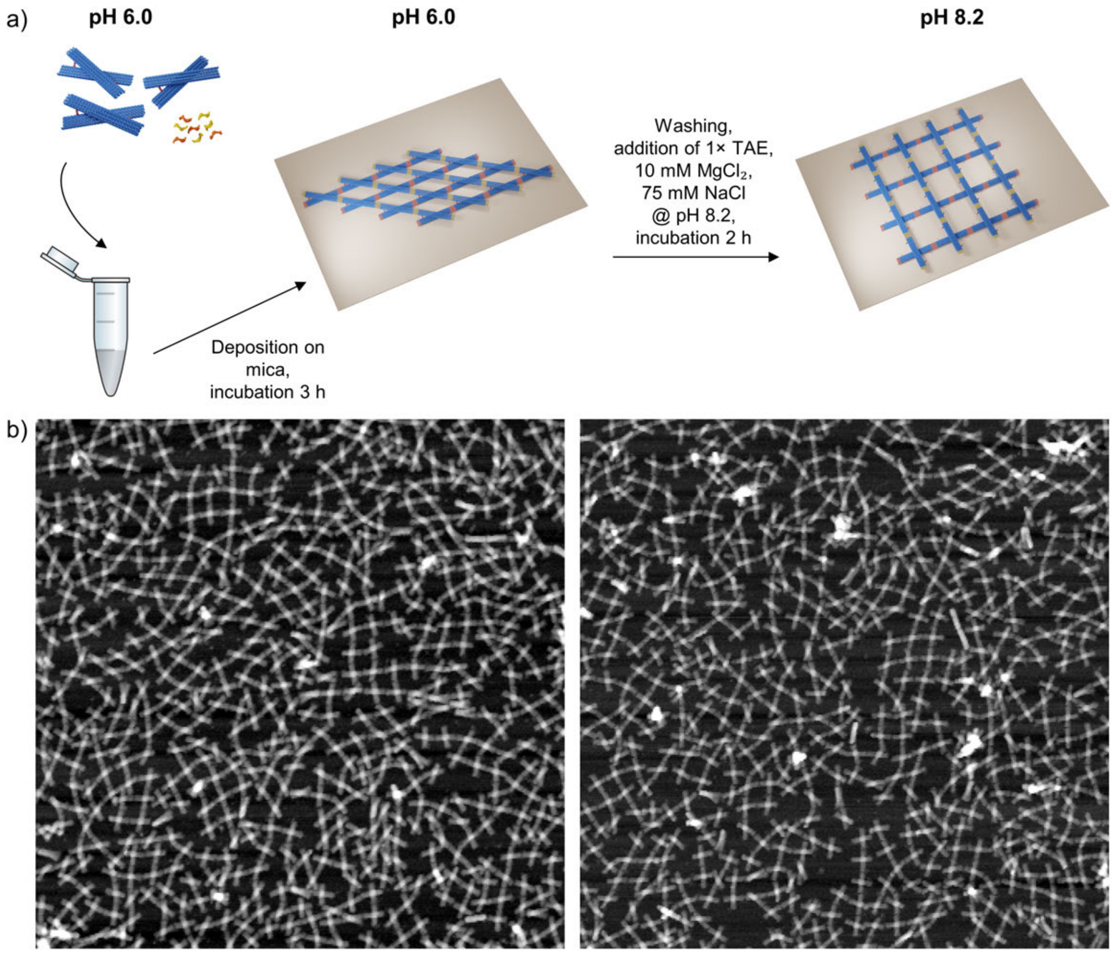


**Figure S30.** AFM images demonstrating the pH-responsiveness of the assembled lattices. (a) The lattices are assembled at pH 6.0 during 3 h. Weakly and unbound assemblies are washed away from the surface, after which 1× TAE, 10 mM $MgCl_2$ and 75 mM NaCl at pH 8.2 is deposited onto the surface to increase the pH to 8.2. To allow the lattice to rearrange, the sample is incubated for additional 2 h. (b) AFM images of the formed lattice after the pH has been increased to 8.2. The size of the images is 2 $\mu$m × 2 $\mu$m

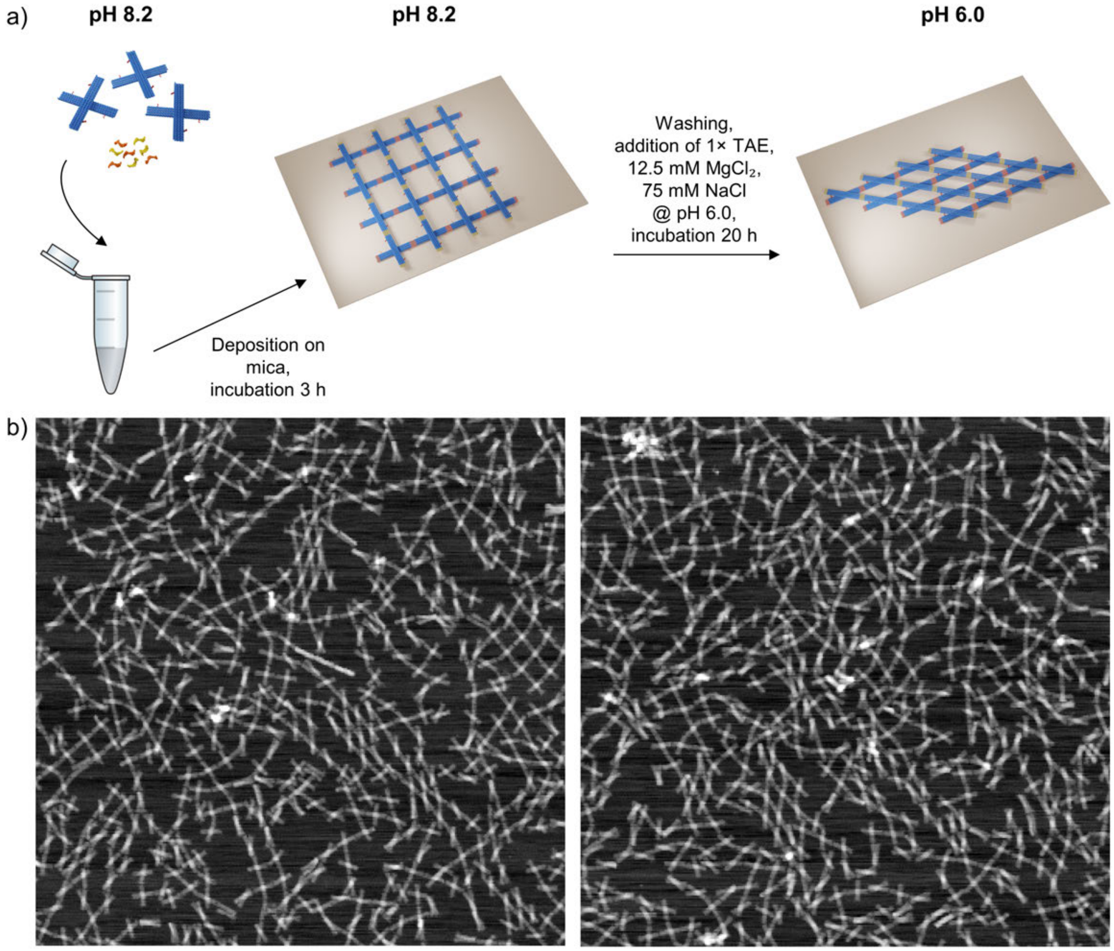


**Figure S31.** AFM images demonstrating the pH-responsiveness of the assembled lattices. (a) The lattices are assembled at pH 8.2 during 3 h. Weakly and unbound assemblies are washed away from the surface, after which $1\times$ TAE, 12.5 mM $MgCl_2$ and 75 mM NaCl at pH 6.0 is deposited onto the surface to decrease the pH to 6.0. To allow the lattice to rearrange, the sample is incubated for additional 20 h. (b) AFM images of the formed lattice after the pH has been decreased to 6.0. The size of the images is 2 $\mu$m $\times$ 2 $\mu$m.

#### 7. DNA-Templated, pH-Responsive, and Reconfigurable 2D AuNP Lattices

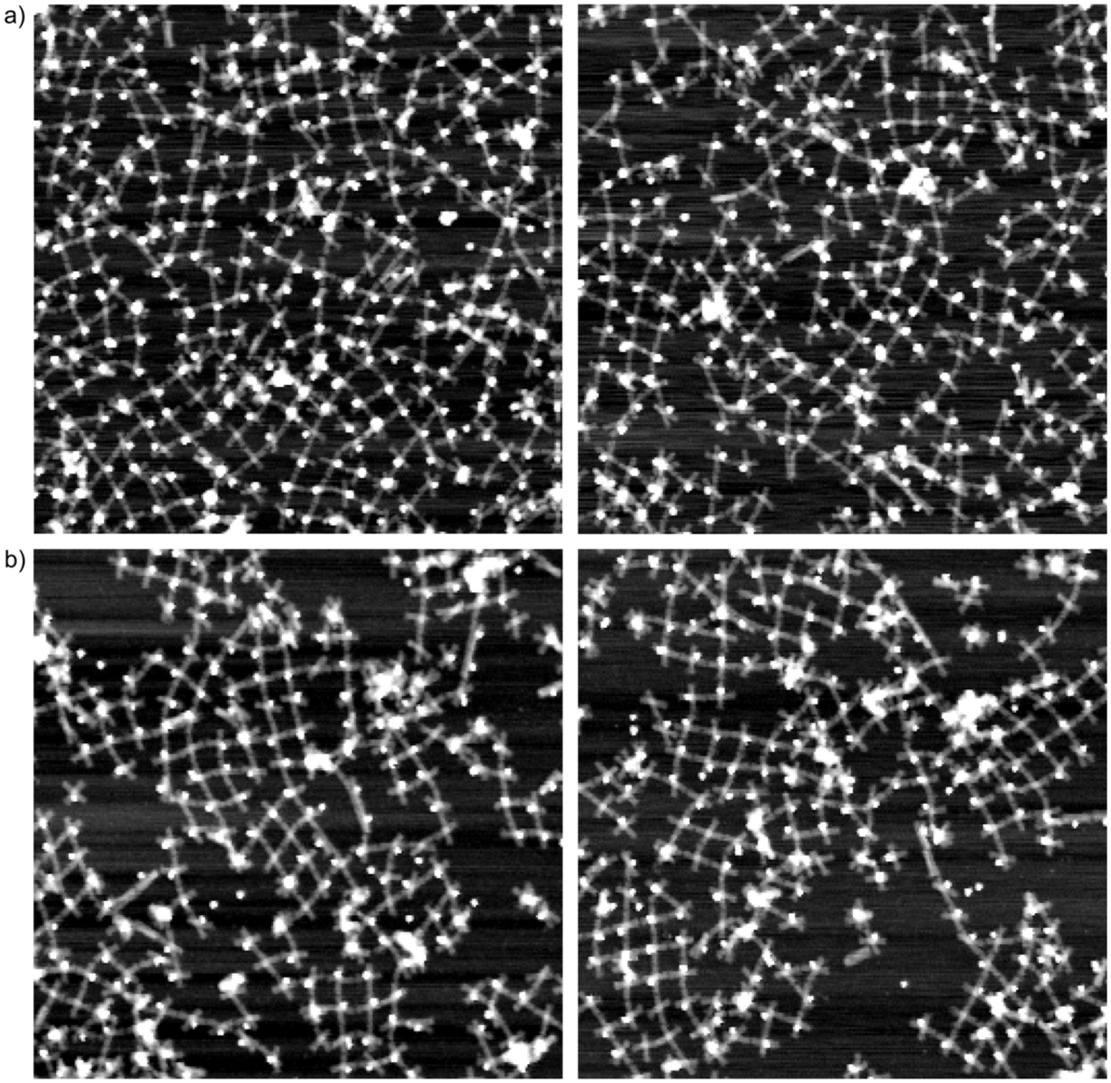


**Figure S32.** AFM images of AuNP lattices formed on a mica substrate at pH 8.2 using the DNA origami unit with pH latches. The assembly is done in 1× TAE containing 10 mM $MgCl_2$ and 75 mM NaCl. The size of the images is 1.5 μm × 1.5 μm. The DNA origami concentration in the samples is (a) 2.0 nM and (b) 3.0 nM (calculated based on the concentration in the conjugation step, assuming no loss during the PEG purification).

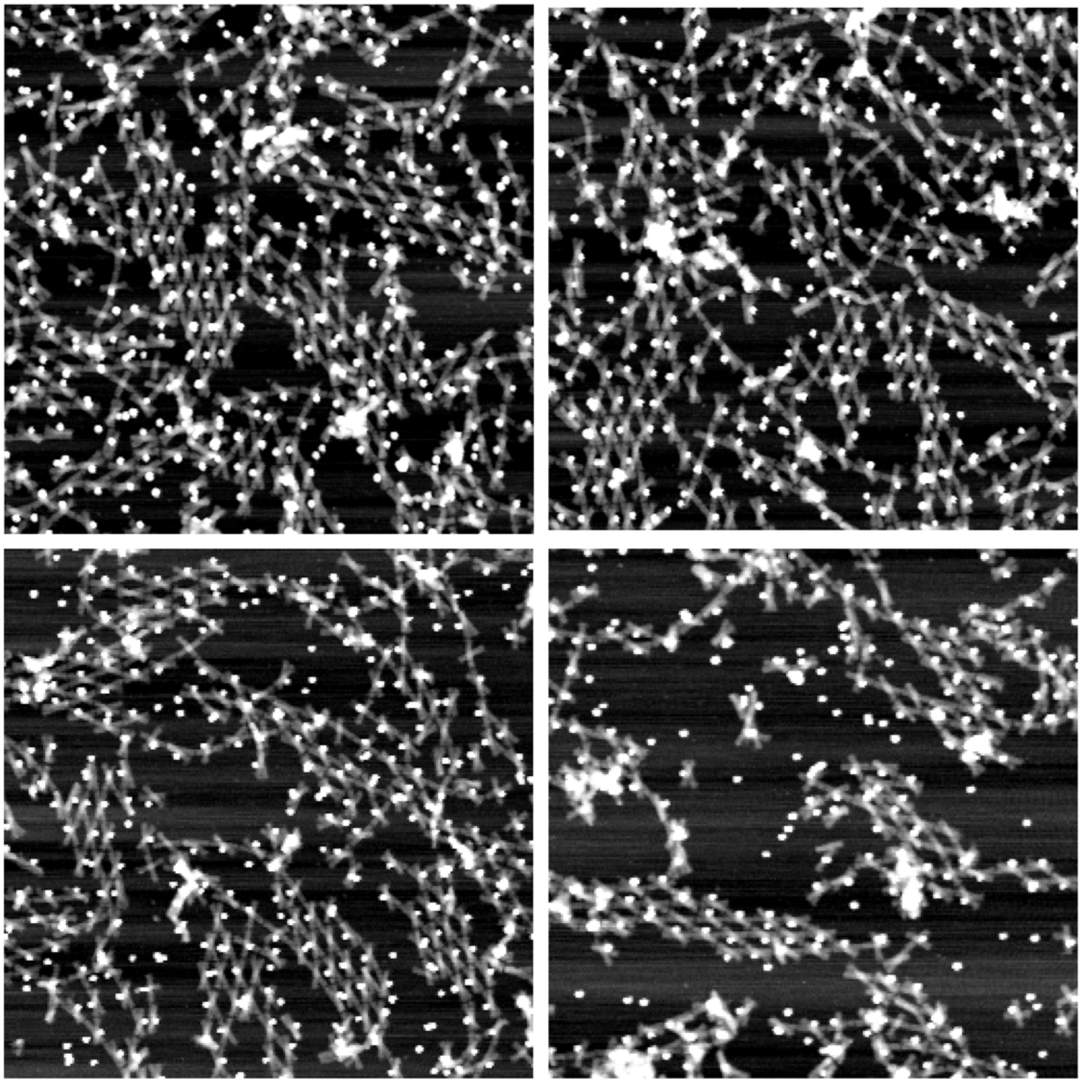

**Figure S33.** AFM images of AuNP lattices formed on a mica substrate at pH 6.0 using the DNA origami unit with pH latches. The assembly is done in 1$\times$ TAE containing 12.5 mM $MgCl_2$ and 75 mM NaCl. The size of the images is 1.5 $\mu$m $\times$ 1.5 $\mu$m. The DNA origami concentration is 3.0 nM (calculated based on the concentration in the conjugation step, assuming no loss during the PEG purification).

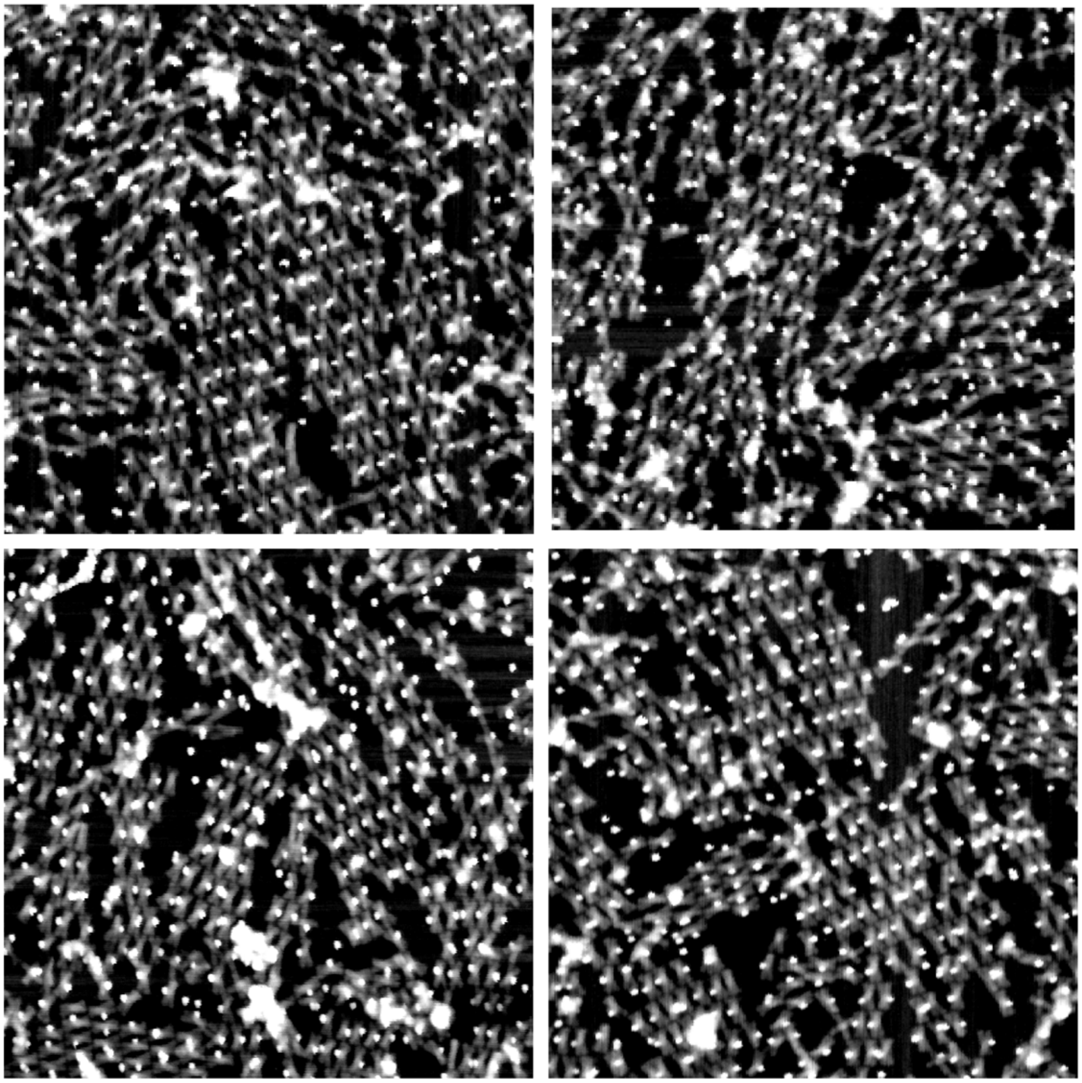

**Figure S34.** AFM images of AuNP lattices formed on a mica substrate at pH 8.2 using the permanently closed unit. The assembly is done in 1× TAE containing 10 mM $MgCl_2$ and 75 mM NaCl. The size of the images is 1.5 µm × 1.5 µm. The DNA origami concentration is 3.0 nM (calculated based on the concentration in the conjugation step, assuming no loss during the PEG purification).

#### 8. DNA origami Unit Design

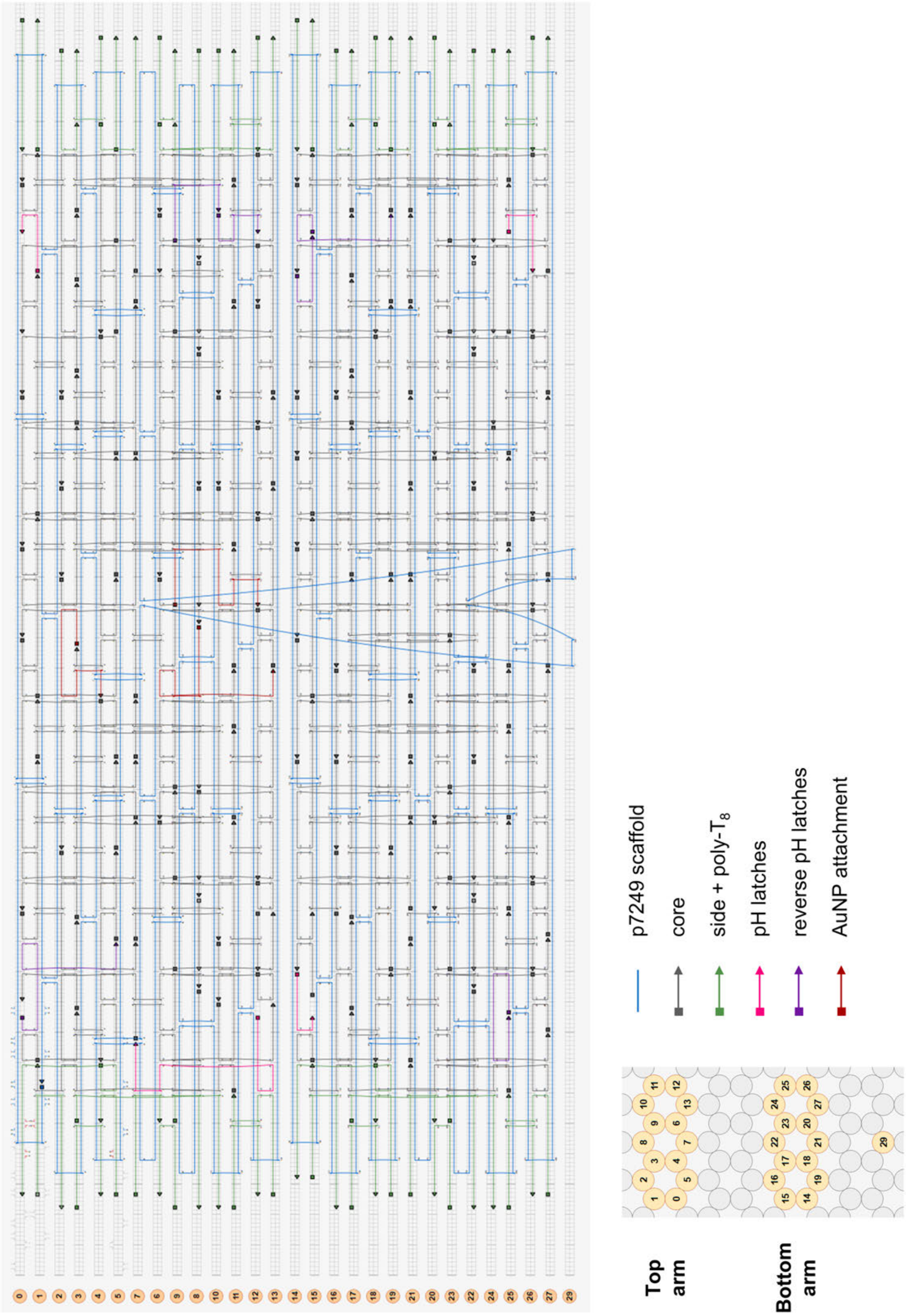


**Figure S35.** The caDNAno design for the DNA origami unit. The unit shown is the poly-T passivated (8-nt polythymine extensions at each helix to avoid end-to-end stacking) version and the staple strands are colored based on their type and/or function.

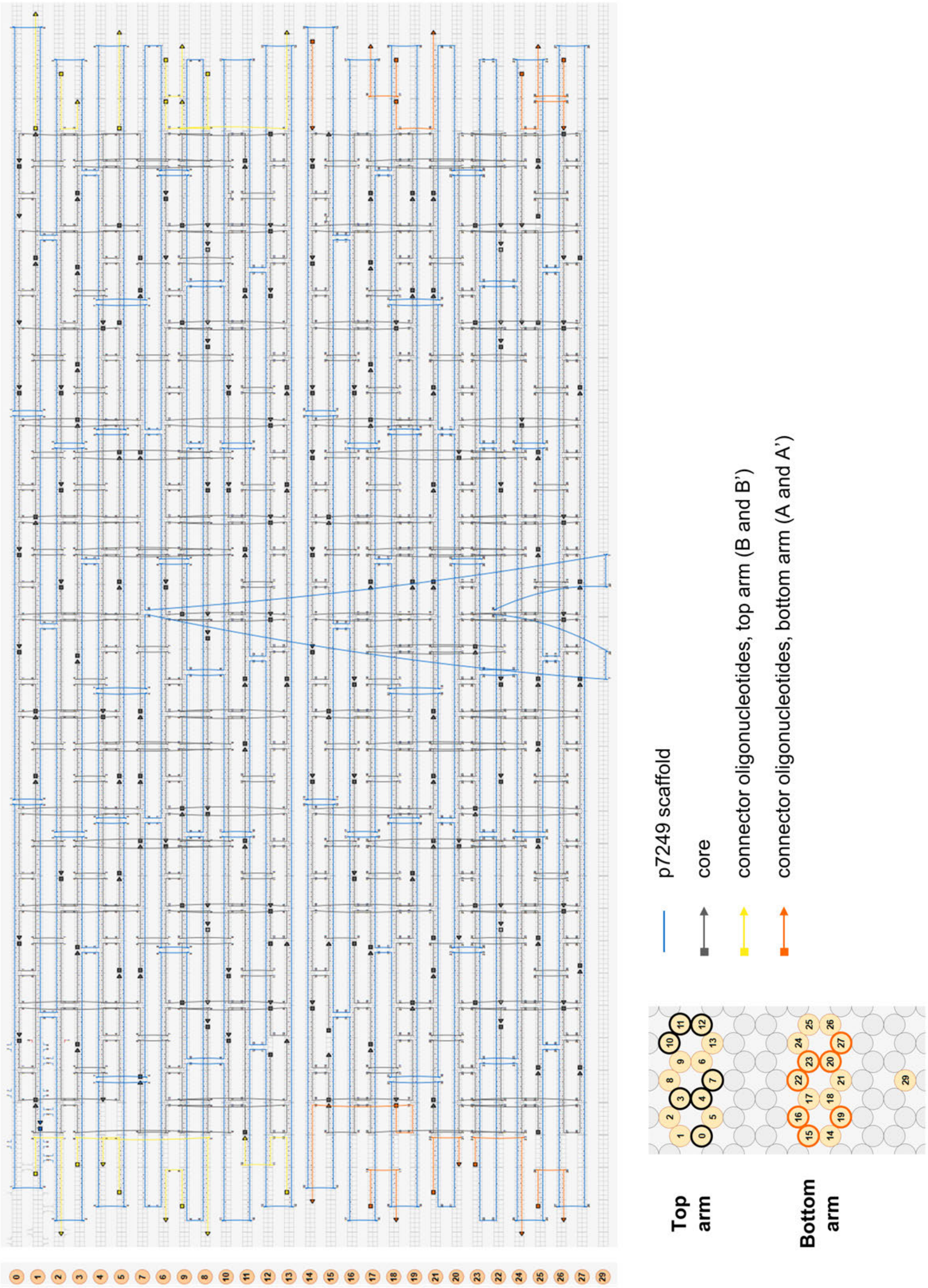


**Figure S36.** The design and position of the connector oligonucleotides used to selectively link the units together. Each of the two arms have connector oligonucleotides on seven of the helices and these helices are indicated in the helix map shown in the bottom-right part of the figure. 14 (7 per arm) of the connector oligonucleotides have a 3-nt overhang (in the 3' end) complementary to the scaffold sequence on the opposite end of the arm.

All versions of the DNA origami unit contain the core staple strands listed in **Table S2**. Depending on the unit type and wanted functionality, some of the core staple strands in **Table S3** could be replaced with staple strands in **Table S4** (pH latches), **Table S5** (sequences for permanently closing the unit), **Table S6** (reverse pH latches) or **Table S7** (AuNP attachment strands). The pH latches and the sequences for permanently closing the unit has been adapted from Ijäs et al. (3). The side strands with the poly-$T_8$ overhangs are listed in **Table S8**. For selectively connecting several units together, these side strands could be replaced with the connector oligonucleotides in **Table S9**. The replacement is always done for the whole set of staple strands of the same type, for example, the whole set "A of poly-T passivated side-strands" would always be replaced with the whole set "bottom arm, A".

**Table S2.** The sequences for the core staple strands. The staple strands are written in the 5' to 3' direction and the Start – End location refers to the position in the caDNAno design.

| Type | Start - end | Sequence |
|---|---|---|
| core | 0[146] – 5[160] | CGATTTTAAACAGGGAAGCCCTGAATCTATCCAAAGCAAATC |
| core | 0[167] – 2[161] | TATCCCATACCAACCCAGCTA |
| core | 0[202] – 3[207] | TTAACGCTTCATCAATATAATATTATCAATTTT |
| core | 0[251] – 3[244] | GCAAAAGGCCTGATCATCGGGAGAAACACTTAGAT |
| core | 0[83] – 3[81] | GTTTGGATTCCGAAAGGCGAAAATCCTGAATCAGAGCGGG |
| core | 1[119] – 1[118] | TTTACAGACCCTGAACAAAGTCAGAGGGTCAAAAGAACAGCC |
| core | 1[133] – 3[144] | AACATAATTGTTTACATTACCGCGCCCAATAGCAAACAAG |
| core | 1[175] – 5[188] | AGCGTCTACAGCCATATCCGGTATTCTA |
| core | 1[49] – 3[60] | TGTAGCGACGTGGACGCGGGGAGAGGCGTAAAGGGGCTTT |
| core | 10[181] – 12[175] | TCAGCTACAGACGAGCAGAGG |
| core | 10[202] – 13[202] | TATGCGTTATACACCGCGTTAAATAAGAGCGGAACCAGTGCC |
| core | 10[62] – 10[63] | TGTTAACCTACATTACACGACCAGTAAGCTTGCATGTGAAAT |
| core | 10[76] – 12[70] | TTCCTGTGCCTGCAACGACGG |
| core | 11[105] – 6[105] | GAATCGGTGCGGGCAGTTGGGTAACGCCAGCCCTATACACCG |
| core | 11[126] – 1[132] | CTATTACAGGCTGCCCAGGCAATCGCACCAGAGAGACTGAACAGAGAAT |
| core | 11[140] – 11[139] | GGCGAAGTACCGACCAACAATAGATATCGCCATTCGCCAGCT |
| core | 11[168] – 1[174] | TTCTGTCATGCAGAGGTCACGCTGCCAGCGGGAGGTTTGCACGCTAACG |
| core | 11[224] – 8[234] | GTAGGTCTAAAATACTTTACACAACTCGTGAC |
| core | 11[252] – 1[258] | AACAACTTGAGGATATCGCAAATTTTAGTTTTCAGCCTTTTATGCTTTG |
| core | 11[42] – 1[48] | ACCAGTCTTGACGCAAACGCTTGGTAATAGCACGTCGCTTAACAGCAAG |
| core | 12[153] – 0[147] | AGAATATGCAGAAGTACGAGCCCGAAGCCAATGAAATAGCAATAAGAAA |
| core | 12[174] – 0[168] | CATTTTCAGGCGGTGTAATGGTAACAACCTCCCGAGAAGGCTTATTATT |
| core | 12[195] – 10[182] | AACATGTCCTGCAAAAAACAAACGGCGGAGATGGGAACATGT |
| core | 12[223] – 7[223] | ATTGAGAATCGCCAGAGCCAGAACATTAAGCGGAATTATCCA |
| core | 12[237] – 6[231] | GGAAGGTTCTAAAGATTAATT |
| core | 12[258] – 0[252] | TCAACAGACCTCAAATGCAAATAACCTCAATTGCGAACCTACTACCTGA |
| core | 12[69] – 0[63] | CCAGTGCAATATTTCATAAAGGAACGGTGGCCGATGTTTGCGTCCACTA |
| core | 12[83] – 8[87] | TTGTAAAGGTCGACTCTAGAGATGGTCAAATTAACCATC |
| core | 12[90] – 0[84] | CACGACGGTCTTTATCCATCAAGAAGTGTTCACCGGTGGTTTTGTTCCA |
| core | 13[140] – 7[132] | ACCACCAAAAAGGGGGATGTGCTGCAAGAAAATACCAATAATTATTAAA |
| core | 13[182] – 12[196] | AACACCGAATTTAGCGACAATAAAAATTCTTACCACAACGCC |
| core | 13[203] – 8[213] | ACGCTGATATTTAAGTATAAAGCCAACGCCTGTTTCCGGAATAAAT |
| core | 14[118] – 22[119] | CTAGCGGTGGCATCAATTCTAGGGCGCGTGAACGGTACCAGA |
| core | 14[146] – 19[160] | ATCACCACTATTTTTGAGAGATAAAAATCCGGAGAAATCATA |
| core | 14[167] – 17[160] | AGAAAGGTTTTAGATTTATTTCAACGCATGTACCC |
| core | 14[188] – 21[181] | AGGTAAATAAATGCAATACTTTTGCGGGAAAACATGCATAAATATTTTG |
| core | 14[202] – 17[207] | ATTTGTCCGCGACCTGCTCCGATAAATTCATC |
| core | 14[251] – 17[244] | CTTGAGACAAAGCTATATTCATTACCCAAATGCAG |
| core | 15[133] – 22[140] | AGGGTAGTCAATATTAGTAGCATTAACAAAAACAGGCATGTCAAGCAGA |
| core | 15[175] – 14[189] | TATATTTGATTCAAAATTAGCAAAATTAAGCAAATAATGTGT |
| core | 15[49] – 17[60] | TACAGGATCTGAATGTCATAGCCCCCTTGAAGGATTTATT |

| | | |
|---|---|---|
| core | 15[64] – 14[70] | GTAAGAGCCGCCGGCCAGAA |
| core | 16[118] – 16[119] | GTCTGGAGCAAAGACAGACGATTTGATAAATTAATCCTGAGA |
| core | 16[139] – 14[119] | CTATCAGAAAACTAGATATTTTCATTTGCTAATAGGATATTCAACCGTT |
| core | 16[202] – 16[203] | CAACATTATTACGTAATGCCTGAAAGAACTGGCTCAACGGAA |
| core | 16[83] – 17[81] | ACCAGAACCACCACGAAAGTATTAAG |
| core | 16[97] – 21[104] | CCCTCAGAGAACCGCGCCTCCTGCCGTCGAGAGGG |
| core | 17[161] – 27[160] | CGGTTGTTAGCCAGCTTTCAACATTCCAGATTCCCCGGTTTA |
| core | 17[196] – 21[202] | GAAAGATTGTGTCGCGCGAAA |
| core | 17[208] – 19[223] | AGTTGAGATACGAACTATTATACCAGTCAGCATTGTGTTAGCCG |
| core | 17[230] – 18[217] | ATTCAACTAAAAAAATCTACGTTAATAAATTAGGAAAGGTATC |
| core | 17[245] – 22[238] | ATACACGCGGAATCGTCATAA |
| core | 17[61] – 18[49] | CTGAAACATCATAAGTTTTAACGGGGTCAGGAACCTATATCAAG |
| core | 17[82] – 19[97] | GCCACCCTCAGCCGCCTGACAGGAGGTTGAAACAAATAAATCAC |
| core | 18[216] – 14[203] | ATCGCCTATGTTACAATTACCTTATGCG |
| core | 19[161] – 21[160] | CAGGCAACGGATAATCAGAAATATTTAA |
| core | 19[224] – 17[229] | GAACGAGGAGGCATTACCAC |
| core | 19[245] – 21[244] | TCATAAGGGTAACGCCAAAAGATAACCC |
| core | 19[70] – 21[76] | TTTGCCATCTTTTCCCAGGCTGAGACTCGCTCAGT |
| core | 19[98] – 16[98] | CGGAACCAGAGCGGCCTTGATATTCACAGGCAGGTGCCACCA |
| core | 2[160] – 8[154] | CAATTTTAAGCCCAATAATTGTTTGAGGGGACGAC |
| core | 2[181] – 7[188] | TTGCTATTTTTGAAAGCGAACCCGTCGGATTCTCC |
| core | 2[202] – 2[203] | TCTGTAAATCGTCAGCCTAATTTAATTTCATTTGACCTTGCT |
| core | 2[97] – 7[104] | TGGTTTGGAGAGTTATTGCCCTTTTTATAATCAGT |
| core | 20[104] – 26[91] | TATAGCCTAGTACCAATTATTCATTAAAAAGGTAAATAGAAA |
| core | 20[188] – 26[175] | TTTTTGTCCATCAAATAATGCTGTAGCTAGAGCTTACAGGTC |
| core | 21[105] – 22[91] | TTGATCGAACACCACCGGAACCCACCCTCCCTCAGAACCGCC |
| core | 21[119] – 23[111] | TCAATAAACCATTAAACGGAATATTACGCAGTATGTTAGCGAGCAAACG |
| core | 21[133] – 23[146] | AGCTAAGATTGTATCGAACGAGGGTCTG |
| core | 21[161] – 26[154] | ATTGTAATTCAGTTTATAATTTTTTAAATATGCAAGGTCATTCCTTTAA |
| core | 21[182] – 22[182] | TTAAAATTCGCAAGAATAAAGCCTCAGATATGACCAATTCGC |
| core | 21[203] – 24[196] | CAAAGTACCCAGCGATCAAAAAGCAAAGCGGATTG |
| core | 21[224] – 22[234] | AGAGCAACTAAAACCTCTACGAAGGCACATAT |
| core | 21[245] – 25[251] | TCGTTTAGCAAAGAGGCAAAAACGCATAACCGATACACCCTCCAACGGC |
| core | 21[49] – 22[66] | CAGAAGGATTAGCGTAGCAAGAATGGGAATTAGAGAGAG |
| core | 21[77] – 27[77] | ACCAGGCCACCAGTAGCACCAAACAACTA |
| core | 21[84] – 15[83] | GGATAAGCTCAGAGATAATCAAAATCCTCATTAAACCAGCAT |
| core | 22[118] – 21[118] | AGGAAACCGAGGCACACAAGAGAATCGAAGCTGAACAAATGG |
| core | 22[139] – 27[139] | TAGCCGAAAGAACTGTAGATTAAAAGGA |
| core | 22[181] – 25[188] | GTCTGGCAGGAACGTAAATCAAGGTGAATTTCTTAAACTCCAAATTGCT |
| core | 22[212] – 17[195] | CTTTAAACAGTTCAGAAAACGAATCTAGGTA |
| core | 22[233] – 22[217] | TCATTGAATCCCCCTCA |
| core | 22[65] – 27[55] | CCACCACCCTCATTTTCCCAGTACGCCGGAAAGCGGAG |
| core | 22[86] – 22[70] | TCAGAACCGCCACCCTC |
| core | 23[112] – 25[125] | CAATAATGATACATAATCTCCAAAAAAAACGGAATTACATAC |
| core | 23[147] – 16[140] | GAAGTTTGAAAAGTAATCATAAGGATCTACAAAGG |
| core | 23[217] – 26[217] | CTTTACCACTCATCCAATGACAACAACCGCCGTTTTAATTCG |
| core | 23[238] – 24[238] | CAACCTAAAACGCATTAAACGGGTAAAA |
| core | 23[70] – 26[70] | CCAGCAAAATCACCTATCACCGTCACCGATTCAACCCAGCGC |
| core | 23[91] – 25[104] | GGAGGTTCGGAATATAATAATTTTTTCAACAATCAATATTGA |
| core | 24[195] – 27[202] | CATCAATAAGAATGACCATAAATTATACTGATACC |
| core | 25[105] – 20[105] | CGAAACGTAGAAAAAAGTTTATTTTGTCCGTTGAATTATAAG |
| core | 25[126] – 15[132] | ATAAAGGGACTCCTTACCCAAACAAAGTTAATCGTGTCATTGGCCGGAG |
| core | 25[140] – 25[139] | ATATATTGATAAGACTAAAGTACGGTCATGATTAATGGCAAC |
| core | 25[168] – 15[174] | ATGGCTTCAACATGAACCAATCTTCCTGTGTACCAAGAAGCCACCCTCA |
| core | 25[189] – 20[189] | GAAAAAGATTAAGACCAGACCGGAAGCAAACAGCTCATTAAA |
| core | 25[217] – 24[217] | CTTCAAATATCGATCGGAACGTACGTAATGCCAGACTATTAT |
| core | 25[252] – 15[258] | TACAGAGGTTTCCAGAGGGGGAATACTGCATAGGCGAACCGGGCTCATT |

| | | |
|---|---|---|
| core | 25[42] – 15[48] | AAAGTTTCCTGTAGTTCGTCAAGGGATATATTTCGTGCCTTGTTGATGA |
| core | 25[84] – 22[87] | GGGAGGGGGTGAATGTACTCAACCC |
| core | 26[153] – 14[147] | TTGCTCCATTGTATAATTCTGAAGCAAAAGCCCCATCCAATACAGTCAA |
| core | 26[174] – 14[168] | AGGATTAGCTTTCGGCTCATTACGTTAAGCTAAATGGCAAAGAAGGGTG |
| core | 26[216] – 21[223] | AGCTTCAGCGCCGATTTGACCCAACGGAGATTTTA |
| core | 26[258] – 14[252] | GGATCGTTATTCGGAAGTTTTCCAGACGCAGACCAGGAACCGAATTGGG |
| core | 26[69] – 19[69] | CAAAGACAGAAAGGTTACCATGGGTTTTCTCAAGAATTAGCG |
| core | 26[90] – 20[84] | ATTCATATTGCGAAGGTGTAT |
| core | 27[140] – 21[132] | GCCTTTATTAAAGAAACGCAAAGACACCAGGCTCCTAGTTTGCCTGTTT |
| core | 27[161] – 25[167] | TCAGCTTGAGAGTATTTGCGG |
| core | 27[203] – 22[213] | GATAGTTAAGCGAAGGAAGCCCGAAAGAAGTCAGAATCAGGTAATG |
| core | 27[231] – 14[231] | ATCGCCCGAATACACACTATCGAATTACGCGCAGAATTTCAA |
| core | 27[56] – 21[48] | TGAGAATAAAGACGTTAGTAAATGAATTCAGTTTCACGTCACGTAGCGA |
| core | 27[78] – 25[83] | AAGGAATGGTTTACGATTGA |
| core | 3[196] – 10[203] | TTAATTAGATGATGAAACCACCAGAAGGTCATTTTATAAACAAGTATCA |
| core | 3[208] – 0[217] | CCCTTAGAATGAATAAATTACCTTTTTAAAGAAAAC |
| core | 3[229] – 4[217] | TAGCGATAGATATAAATCAATATATGTGAGTCCTTGAATATCAT |
| core | 3[245] – 8[238] | TAAGAGATTAATTTCATCTTC |
| core | 3[61] – 4[49] | CCTCGTTAGTTGTAACCACCACACCCGCCGATAACGTATTTTCC |
| core | 3[82] – 5[97] | TGGCCCTGACCCCAGCATCGGCAAAATCCCTGAGTGTTTCTTTT |
| core | 4[132] – 5[118] | CACTCATTAGGAATACGTCAAAAATGAAAATAGTATTTTATT |
| core | 4[216] – 0[203] | ATTCCTGCCTGATTAAAATTAATTACAT |
| core | 5[119] – 8[112] | TTCATCGCGAGAACCTAATATTCCAGCCAGCTTTC |
| core | 5[161] – 6[161] | AGATATACTAAGAGCAAGAAACAAATGTGAGCGAGGACCCAT |
| core | 5[189] – 2[182] | AGAACAACAGTTACAAAATAATTCCAGAAGATTAG |
| core | 5[217] – 3[228] | GTTTGGATTATACTTGAATTTAAACA |
| core | 5[238] – 6[245] | AATGGAAGGGTTAGTACGCTGAGAAGAGAGAGACTACCTTTTTCTGCCC |
| core | 5[77] – 7[76] | CGCCAGGCCAGCTAAACAGGAACGCCAG |
| core | 5[98] – 2[98] | CACCAGTGAGACGTGCCCGAGATAGGGTTTATAAATCCACGC |
| core | 6[104] – 12[91] | AGTAAAAGCAATACTCGAATTCGTAATCGATCCCCTCCCAGT |
| core | 6[160] – 11[167] | CCTAATTATAAAACAGAGGTGGAGCCAGAAAGTAA |
| core | 6[244] – 11[251] | GAACGTTCATCACCTTGCTGATTGAAAGGAGCACT |
| core | 7[105] – 8[91] | GAGGCTCCCGGGCAACAGCTGGCAGCAATTGATTAGTAATAA |
| core | 7[119] – 9[111] | GAACGGGCGGCTGTCCGGAAAGCAACTGTTGGGAAGGGCGGCTTCCGCT |
| core | 7[133] – 8[150] | CCAAGGCTATCTTAATGTAGAATAGTCCTGAACAAGACA |
| core | 7[189] – 8[182] | GTGGGAGGCGCGAGGCGTTTTGCCTTAATCGTAAC |
| core | 7[224] – 11[223] | AAATCATTTAAAAGACTTCGAAACAATAGAAAAAGCTCAACA |
| core | 7[231] – 1[230] | AGGTCTGTCAATAGTCTGAATTGAAACAAACATCATGGAAAC |
| core | 7[54] – 8[66] | TTTAGACAGTGTAAAGATTCCGCTCACAATAACT |
| core | 7[77] – 13[83] | AATCCTGCGACGAGCCGGAAGTTGAATGGCTATTA |
| core | 8[111] – 7[118] | CGGCACTGCGGTAATTGAGCGAAGCAAGATTCCAA |
| core | 8[181] – 13[181] | CGTGCATTTGGTGTATTGACCCAGTATT |
| core | 8[212] – 3[195] | ACCGACCGTGTGATAAATAAGGCAATCGCTA |
| core | 8[233] – 8[217] | CTAAATTTAATGGTTTG |
| core | 8[65] – 13[62] | CAAACTATCGGCCTTGCCATGGAACCTGGGGTACGTGGCACAGAC |
| core | 8[86] – 8[70] | ACTTGCCTGAGTAGAAG |
| core | 9[112] – 11[125] | TCTGGTGCTTTCCTAAACATCGCCATTAGCGATTACTCTTCG |
| core | 9[217] – 12[224] | CATAATTTTTGAGTCAGCAAATGAAAAATAGCTTA |
| core | 9[70] – 10[77] | TCCACACAACATCATAGCTGT |
| core | 9[91] – 11[104] | CGTTGTAGAGTCTGATGCGCGAACTGATAGGGTTTGGGTACC |

**Table S3.** The sequences for the core strands that could be replaced to get a certain functionality. The staple strands are written in the 5' to 3' direction and the Start – End location refers to the position in the caDNAno design.

| Type | Start - end | Sequence |
|---|---|---|
| core, pH latch 1.1 | 12[59] – 7[53] | TAAAAGGGACATTCTGGCGTAAGAATGCCTAACTCACTGCCCG C |
| core, pH latch 1.2 | 14[69] – 15[59] | TGGAAAGCGCAGTCGTGT |
| core, pH latch 2.1 | 1[231] – 0[240] | AGTACAACGGATTCAAGAT |
| core, pH latch 2.2 | 25[240] – [26]231 | GGTAGAGCAGCGAAAGACA |
| core, reverse pH latch 1.1 | 0[62] – 5[76] | TTAAAGAGTCACGCTGCGCTGATGGTGGACAAGAGTATTGGG |
| core, reverse pH latch 1.2 | 25[56] – 25[55] | TTTCCAAGGGCGACACTTGAGCCATTACTACAACGTGTCGTC |
| core, reverse pH latch 2.1 | 9[238] – 12[238] | TATTAAATCCTTCATTAGAAGTATTAGATCTTTAGGAATTGA |
| core, reverse pH latch 2.2 | 14[230] – 19[244] | CTTTAATGACGTTGGGAAGTCAACGTAATGGTTTACGGTCAA |
| core, AuNP attachment | 3[145] – 4[133] | AATTGAGTTATGCATTAGACGGGAGAATTAATAACCCTATACCG |
| core, AuNP attachment | 8[149] – 13[139] | GTATCGGCCTCAGGAAGAAGCGCCAACCAATCGAACGA |
| core, AuNP attachment | 9[154] – 12[254] | GAAAAATAATATTAACGCGCCTGTTTATAAAAGGTTAATAAG |

**Table S4.** The sequences for the pH latches. The staple strands are written in the 5' to 3' direction and the Start – End location refers to the position in the caDNAno design. The hairpin-forming extensions and the ssDNA counterparts have been colored red, whereas the spacers are marked with lowercase letters.

| Type | Start - end | Sequence |
|---|---|---|
| pH latch 1.1, hairpin | 12[59] – 7[53] | AGAGAAGAAAAGAGGAAGGAttttTCCTTCCTCTTTTCTTCTCTttttTA AAAGGGACATTCTGGCGTAAGAATGCCTAACTCACTGCCCGC |
| pH latch 1.2, ssDNA | 14[69] – 15[59] | TGGAAAGCGCAGTCGTGTttttTCTCTTCTTTTCTCCTTCCT |
| pH latch 2.1, hairpin | 1[231] – 0[240] | AGTACAACGGATTCAAGATttttCCTTTCTTTCTTTCTTCCTCttttGAGG AAGAAAGAAAGAAAGG |
| pH latch 2.2, ssdNA | 25[240] – [26]231 | CTCCTTCTTTCTTTCTTTCCttttGGTAGAGCAGCGAAAGACA |

**Table S5.** The sequences for permanently closing the unit. The staple strands are written in the 5' to 3' direction and the Start – End location refers to the position in the caDNAno design. The ssDNA regions forming the duplex have been colored red, whereas the spacers are marked with lowercase letters.

| Type | Start - end | Sequence |
|---|---|---|
| latch 1.1 - closed, ssDNA | 12[59] – 7[53] | TCCTTCCTCTAAAGAAGAGAttttTAAAAGGGACATTCTGGCG TAAGAATGCCTAACTCACTGCCCGC |
| latch 1.1 - closed, ssDNA | 14[69] – 15[59] | TGGAAAGCGCAGTCGTGTttttTCTCTTCTTTAGAGGAAGGA |
| latch 2.1 - closed, ssDNA | 1[231] – 0[240] | AGTACAACGGATTCAAGATttttCCTTTCTTTCAAAGAAGGAG |
| latch 2.2 - closed, ssdNA | 25[240] – 26[231] | CTCCTTCTTTGAAAGAAAGGttttGGTAGAGCAGCGAAAGAC A |

**Table S6.** The sequences for the reverse pH latches. The staple strands are written in the 5' to 3' direction and the Start – End location refers to the position in the caDNAno design. The hairpin-forming extensions and the ssDNA counterparts have been colored red, whereas the spacers are marked with lowercase letters.

| Type | Start - end | Sequence |
|---|---|---|
| reverse pH latch 1.1, hairpin | 0[59] – 5[76] | AAAAGGGAGAAGAAAAGAGGttttCCTCTTTTCTTCTCCCTTTTttttAAGAGTCACGCTGCGCTGATGGTGGACAAGAGTATTGGG |
| reverse pH latch 1.2, ssDNA | 25[60] – 25[59] | CAAGGGCGACACTTGAGCCATTACTACAACGTGTCGTCTTTCttttTTTTCCCTCTTCTTTTCTCC |
| reverse pH latch 2.1, hairpin | 10[244] – 12 [240] | TATTAGATCTTTAGGAATTttttCCCCCTTTCTTTTTTCTTCTttttAGAAGAAAAAAGAAAGGGGG |
| reverse pH latch 2.2, ssdNA | 15[240] – 19[244] | TCTTCTTTTTTCTTTCCCCCttttCGTAATGGTTTACGGTCAA |
| core | 14[230]–15[239] | CTTTAATGACGTTGGGAAGTCAA |
| core | 9[238] – 10[245] | TATTAAATCCTTCATTAGAAG |

**Table S7.** The sequences for the AuNP attachment. The staple strands are written in the 5' to 3' direction and the Start – End location refers to the position in the caDNAno design. The part complementary to the thiol modified strand is colored red, whereas the spacers are marked with lowercase letters.

| Type | Start - end | Sequence |
|---|---|---|
| AuNP attachment | 3[145] – 4[133] | GAAGGGAGGAAAGTATCGGCCTCAGGAAGAAGCGCCAACCAATCGAACGA |
| AuNP attachment | 8[149] – 13[139] | GAAGGGAGGAAAttAATTGAGTTATGCATTAGACGGGAGAATTAATAACCCTATACCG |
| AuNP attachment | 9[154] – 12[254] | GAAGGGAGGAAAttGAAAAATAATATTAACGCGCCTGTTTATAAAAGGTTAATAAG |
| 3' thiol modified strand | – | TTTCCTCCCTTCTTT/3ThioMC3-D/ |

**Table S8.** The sequences for the side strands. The staple strands are written in the 5' to 3' direction and the Start – End location refers to the position in the caDNAno design. The poly-$T_8$ overhangs added to the side strands are written in lowercase letters.

| Type | Start - end | Sequence |
|---|---|---|
| side, polyT$_8$, B | 0[289] – 0[259] | ttttttttAGAGGCGAATTATTCATTTCAAT |
| side, polyT$_8$, B | 1[259] – 1[289] | AATACCAAGTTACAAAATCGCGCtttttttt |
| side, polyT$_8$, B | 2[282] – 3[265] | ttttttttATATACAGTAACAGTAGTTTAAC |
| side, polyT$_8$, B | 4[265] – 7[285] | ATAAAGACGGCTTAGGTTGGGTTATAtttttttt |
| side, polyT$_8$, B | 4[285] – 3[282] | ttttttttCGTAAAACAGAAGTCAGATGAtttttttt |
| side, polyT$_8$, B | 5[259] – 5[285] | CATATCAAAATTATTTGCAtttttttt |
| side, polyT$_8$, B | 6[265] – 13[285] | AATGCTGATATCAAACCCTCAATCAAtttttttt |
| side, polyT$_8$, B | 6[285] – 9[282] | ttttttttTAACTATATGTAAACGCGAGAtttttttt |
| side, polyT$_8$, B | 8[282] – 9[265] | ttttttttAAACTTTTTCAAATATGACAAAG |
| side, polyT$_8$, B | 10[282] – 12[259] | ttttttttCAATAGATAATACATTAATAGATTGGCAAA |
| side, polyT$_8$, B | 12[285] – 11[282] | ttttttttTATCTGGTCAGTTAGAGCCGTtttttttt |
| side, polyT$_8$, B' | 1[22] – 2[15] | ttttttttAAAACCGTCTATTGCGCCGCTACAGGGCGCGtttttttt |
| side, polyT$_8$, B' | 3[15] – 4[18] | ttttttttTACTATGGTTGCTCGTGCCAGtttttttt |
| side, polyT$_8$, B' | 3[35] – 8[15] | TTTGACGATCCAGAACAATATTACCGtttttttt |
| side, polyT$_8$, B' | 4[48] – 0[22] | AGTCGGGGCCAACGCTCCAACGTCAAAGGGCGAtttttttt |
| side, polyT$_8$, B' | 5[18] – 4[35] | ttttttttCTGCATTAATGAATCGAAACCTG |
| side, polyT$_8$, B' | 7[18] – 6[35] | ttttttttCATTAATTGCGTTGCGTGAGTGA |
| side, polyT$_8$, B' | 9[15] – 6[18] | ttttttttCCAGCCATTGCAGCTAACTCAtttttttt |
| side, polyT$_8$, B' | 9[35] – 10[15] | ACAGGAATCAATCGTCTGAAATGGATtttttttt |
| side, polyT$_8$, B' | 11[15] – 12[18] | ttttttttTATTTACATTGGGATAGAACCtttttttt |
| side, polyT$_8$, B' | 13[18] – 11[41] | ttttttttCTTCTGACCTGAAAGCCAACAGACAGATTC |
| side, polyT$_8$, A | 14[289] – 14[259] | ttttttttGAGAAACACCAGAACGAGTAGTA |
| side, polyT$_8$, A | 15[259] – 15[289] | CAGTGAATAAGGCTTGCCCTGACtttttttt |
| side, polyT$_8$, A | 16[282] – 17[265] | ttttttttAGAGTAATCTTGACAATGGCTGA |
| side, polyT$_8$, A | 18[265] – 21[285] | CGGTGTAACGATAAAAACCAAAATAGtttttttt |
| side, polyT$_8$, A | 18[285] – 17[282] | ttttttttAGGACAGATGAACCTTCATCAtttttttt |
| side, polyT$_8$, A | 19[259] – 19[285] | AACTGACCAACTTTGAAAGtttttttt |
| side, polyT$_8$, A | 20[265] – 27[285] | GCAAAAGTCGCTGAGGCTTGCAGGGAtttttttt |
| side, polyT$_8$, A | 20[285] – 23[282] | ttttttttCGAGAGGCTTTTAAAATGTTTtttttttt |
| side, polyT$_8$, A | 22[282] – 23[265] | ttttttttAGACTGGATAGCGTCCTAATAGT |
| side, polyT$_8$, A | 24[282] – 26[259] | ttttttttACTTTTTCATGAGGAAGCTTTGATTTTGCG |
| side, polyT$_8$, A | 26[285] – 25[282] | ttttttttGTTAAAGGCCGCGGACTAAAGtttttttt |
| side, polyT$_8$, A' | 15[22] – 16[15] | ttttttttCATACATGGCTTAGTAACAGTGCCCGTATAAtttttttt |
| side, polyT$_8$, A' | 17[15] – 18[18] | ttttttttACAGTTAATGCCCAGACTGTAtttttttt |
| side, polyT$_8$, A' | 17[35] – 22[15] | CCCTGCCGCAAGCCCAATAGGAACCCtttttttt |
| side, polyT$_8$, A' | 18[48] – 14[22] | TTTGCCTATTTTCGTTACCGTTCCAGTAAGCGTtttttttt |
| side, polyT$_8$, A' | 19[18] – 18[35] | ttttttttGCGCGTTTTCATCGGCTTAGCGT |
| side, polyT$_8$, A' | 21[18] – 20[35] | ttttttttAGCAGCACCGTAATCACAATGAA |
| side, polyT$_8$, A' | 23[15] – 20[18] | ttttttttATGTACCGTAACACCATCGATtttttttt |
| side, polyT$_8$, A' | 23[35] – 24[15] | ACTGAGTCATTCCACAGACAGCCCTCtttttttt |
| side, polyT$_8$, A' | 25[15] – 26[18] | ttttttttATAGTTAGCGTATGGGATTTTtttttttt |
| side, polyT$_8$, A' | 27[18] – 25[41] | ttttttttGCTAAACAACTTTCAATTCTGTAACGATCT |

**Table S9.** The sequences for connector oligonucleotides. The staple strands are written in the 5' to 3' direction and the Start – End location refers to the position in the caDNAno design. The 3-nt long overhangs (in the 3' end) complementary to the scaffold sequence on the opposite end of the arm are written in lowercase letters.

| Type | Start - end | Sequence |
|---|---|---|
| top arm, B | 1[259] – 1[284] | AATACCAAGTTACAAAATCGCGCaaa |
| top arm, B | 2[271] – 3[265] | TACAGTAACAGTAGTTTAAC |
| top arm, B | 5[259] – 5[280] | CATATCAAAATTATTTGCActg |
| top arm, B | 6[265] – 13[280] | AATGCTGATATCAAACCCTCAATCAActt |
| top arm, B | 6[274] – 9[277] | CTATATGTAAACGCGAGAcca |
| top arm, B | 8[271] – 9[265] | CTTTTTCAAATATGACAAAG |
| top arm, B' | 1[33] – 2[20] | ACCGTCTATTGCGCCGCTACAGGGCGCGata |
| top arm, B' | 3[35] – 8[20] | TTTGACGATCCAGAACAATATTACCGaaa |
| top arm, B' | 5[29] – 4[35] | CATTAATGAATCGAAACCTG |
| top arm, B' | 9[26] – 6[23] | GCCATTGCAGCTAACTCAtaa |
| top arm, B' | 13[29] – 11[41] | CTGACCTGAAAGCCAACAGACAGATTC |
| bottom arm, A | 14[278] – 14[259] | AAACACCAGAACGAGTAGTA |
| bottom arm, A | 18[265] – 21[280] | CGGTGTAACGATAAAAACCAAAATAGagc |
| bottom arm, A | 18[274] – 17[277] | ACAGATGAACCTTCATCAaca |
| bottom arm, A | 24[271] – 26[259] | TTTTCATGAGGAAGCTTTGATTTTGCG |
| bottom arm, A | 26[274] – 25[277] | AAAGGCCGCGGACTAAAGata |
| bottom arm, A' | 17[26] – 18[23] | GTTAATGCCCAGACTGTAagg |
| bottom arm, A' | 18[48] – 14[27] | TTTGCCTATTTTCGTTACCGTTCCAGTAAGCGTgag |
| bottom arm, A' | 21[29] – 20[35] | AGCACCGTAATCACAATGAA |
| bottom arm, A' | 23[35] – 24[20] | ACTGAGTCATTCCACAGACAGCCCTCact |
| bottom arm, A' | 25[26] – 26[23] | GTTAGCGTATGGGATTTTgtt |
